# Genome organisation and evolution of a eukaryotic nicotinate co-inducible pathway

**DOI:** 10.1101/2021.04.19.440407

**Authors:** Eszter Bokor, Michel Flipphi, Sándor Kocsubé, Judit Ámon, Csaba Vágvölgyi, Claudio Scazzocchio, Zsuzsanna Hamari

**Affiliations:** University of Szeged Faculty of Science and Informatics, Department of Microbiology, Szeged, Hungary; Institute de Génétique et Microbiologie, Université Paris-Sud, Orsay, France; Department of Microbiology, Imperial College, London, United Kingdom and Université Paris-Saclay, CEA, CNRS, Institute for Integrative Biology of the Cell (I2BC), 91198, Gif-sur-Yvette, France

## Abstract

In *Aspergillus nidulans* a regulon including 11 *hxn* genes (*hxnS, T, R, P, Y, Z, X, W, V, M* and *N*) is inducible by a nicotinate metabolic derivative, repressible by ammonium and under stringent control of the nitrogen-state sensitive GATA factor AreA and the specific transcription factor HxnR. This is the first report in a eukaryote of the genomic organisation of a possibly complete pathway of nicotinate utilisation. In *A. nidulans* the regulon is organised in three distinct clusters, this organisation is variable in the *Ascomycota*. In some *Pezizomycotina* species all the 11 genes map in a single cluster, in other in two clusters. This variable organisation sheds light on cluster evolution. Instances of gene duplication, followed by, or simultaneous with, integration in the cluster; partial or total cluster loss; horizontal gene transfer of several genes, including an example of whole cluster re-acquisition in *Aspergillus* of section *Flavi* were detected, together with the incorporation in some clusters of genes not found in the *A. nidulans* co-regulated regulon, which underlie both the plasticity and the reticulate character of metabolic cluster evolution. This study provides a comprehensive phylogeny of six members of the cluster across representatives of all *Ascomycota* classes.

## Introduction

Nicotinic acid (niacin, vitamin B3), a precursor of NAD and NADP, can be utilised by some bacteria as sole nitrogen and carbon source. The common first step in all investigated prokaryotes is the hydroxylation of nicotinic acid (NA) to 6-hydroxynicotinic acid (6-NA). The further fate of 6-NA is variable; in *Pseudomonas sp.* [1, 2] it is converted to 2,5-dihydroxypyridine (2,5-DP), in *Bacillus* sp. to 2,6-dihydroxynicotinic acid (2,6-NA) [3] and anaerobically to 1,4,5,6-tetrahydro-6-oxonicotinic acid in *Eubacterium barkeri* (formerly *Clostridium barkeri*) [4]. The detailed and variable further bacterial metabolic steps, whether aerobic or anaerobic have been reviewed in [5].

The ascomycete fungus *Aspergillus nidulans* can utilise NA as sole nitrogen source. In common with bacteria, a molybdenum cofactor (MOCO)-containing flavoprotein catalyses the conversion of NA to 6-NA (Purine hydroxylase II, previously called xanthine dehydrogenase II, HxnS [6–9]. The *hxnS* gene is a paralogue of *hxA,* encoding a canonical xanthine dehydrogenase (HxA, Purine hydroxylase I, [10, 11]) which is co-regulated with most other genes of the purine utilisation pathway ([12, 13] and references therein). The substrate specificities of HxA and HxnS have been studied in detail ([11] and references therein). In *A. nidulans* a NA-inducible co-regulated gene cluster is extant (*hxn*1/VI cluster, for cluster I in chromosome VI) comprising six genes, namely, *hxnS, hxnR* (encoding the pathway-specific transcription factor), *hxnP* and *hxnZ* (encoding transporters of the Major Facilitator Superfamily, which could play a role in the uptake of NA and/or NA-derivatives), a putative flavin oxidoreductase (*hxnT)* and a α-ketoglutarate-dependent dioxygenase (*hxnY*) both which may be involved in the further metabolism of 6-NA [11]. In the 1970s, NA non-utilizer mutants were isolated and genetically characterised [6]. These map in *hxnS* and *hxnR*, but also in a second gene cluster in chromosome VI (see below).

The *hxn1/VI* genes are specifically induced by a metabolite of NA catabolism but also by nitrogen starvation ([11] and RNASeq data [14] available at FungiDB). Expression of the *hxn* genes requires both the pathway-specific Zn-finger factor HxnR and the wide-domain GATA transcription factor AreA [11]. The latter mediates de-repression of a wide range of genes in the absence of preferred nitrogen sources (such as ammonium, L-glutamate and L-glutamine) [15–17]. The *hxnR* gene is defined by loss of function mutations which are non-inducible for the six genes of the cluster (including *hxnR* itself) and by constitutive mutations where transcription of all *hxn1/VI* genes occurs in the absence of inducer compounds [11]. The physiological involvement of the hxn1/VI cluster in nicotinate metabolism is further shown by the phenotype of null mutations in the *hxnR* gene, which result in inability to utilise nicotinate, and two of its downstream metabolic derivatives as nitrogen sources [11].

Hereby we complete the description of the genomic organisation of the nicotinate-inducible *hxn* genes by the identification of five additional, HxnR-dependent genes in *A. nidulans*, and we describe variations in the genomic organisation of the eleven *hxn* genes throughout the *Ascomycota* phylum.

The evolution of gene clustering in primary metabolism has been a subject of discussion. Specifically, we do not know which are the factors, which lead to clustering of previously unclustered genes, those involved in clustering maintenance and those eventually leading to de-clustering [18]. Rokas and co-workers have proposed that clustering confers a specific advantage when, in a given metabolic pathway one or more intermediates are toxic as single gene loss, leading to accumulation of a toxic metabolism will be minimised [19, 20]. Toxic intermediates, such as 2,5-DP have been identified in the nicotinate degradation pathway of number of bacteria. Our own work to be published elsewhere (Bokor E., Amon J., Flipphi M., Vagvolgyi C., Scazzocchio C. and Hamari Z.) indicates that these intermediates also occur in *A. nidulans*. Investigating the diverse organisation and evolution of the nicotinate regulon may contribute to this debate.

## Results and Discussion

### Three HxnR dependent, co-inducible gene clusters are extant in *A. nidulans*

In order to search for additional genes involved in nicotinate metabolism we investigated the cluster structure in available ascomycete genomes (see below for a thorough description). Strikingly, in *Cyphellophora europaea* (*Pezizomycotina*, *Eurotiomycetes*, *Chaetothyriales*), five additional genes (to be called *hxnV*, *hxnW*, *hxnX, hxnM* and *hxnN* see below) are positioned between *hxnP* and *hxnR* orthologues, forming a single, 11-gene cluster that includes all orthologues of the *A. nidulans hxnZ, hxnY, hxnP, hxnR, hxnT* and *hxnS* genes [11] (Fig 1, *A. nidulans* cluster 1/VI, table 1). In *A. terreus* (and several other Aspergilli, see below) *hxnV*, *hxnW* and *hxnX* are directly adjacent to *hxnS* (Fig 1). In *A. nidulans* a cluster including *hxnX*, *hxnW* and *hxnV* (cluster 2/VI) is ∼40 kb distant from *hxnZ* (deduced from the re-assembled genomic sequences [21]) while *hxnM* and *hxnN* are adjacent to each other in chromosome I (cluster 3/I). While this article was being written, Martins et al. [22] suggested the clustered organisation we described for *A. terreus* and *A. nidulans* and drew comparisons with a number of other species. However, these authors did not investigate the co-regulation by nicotinate or its metabolites of the putative new *hxn* genes.

**Fig 1.**
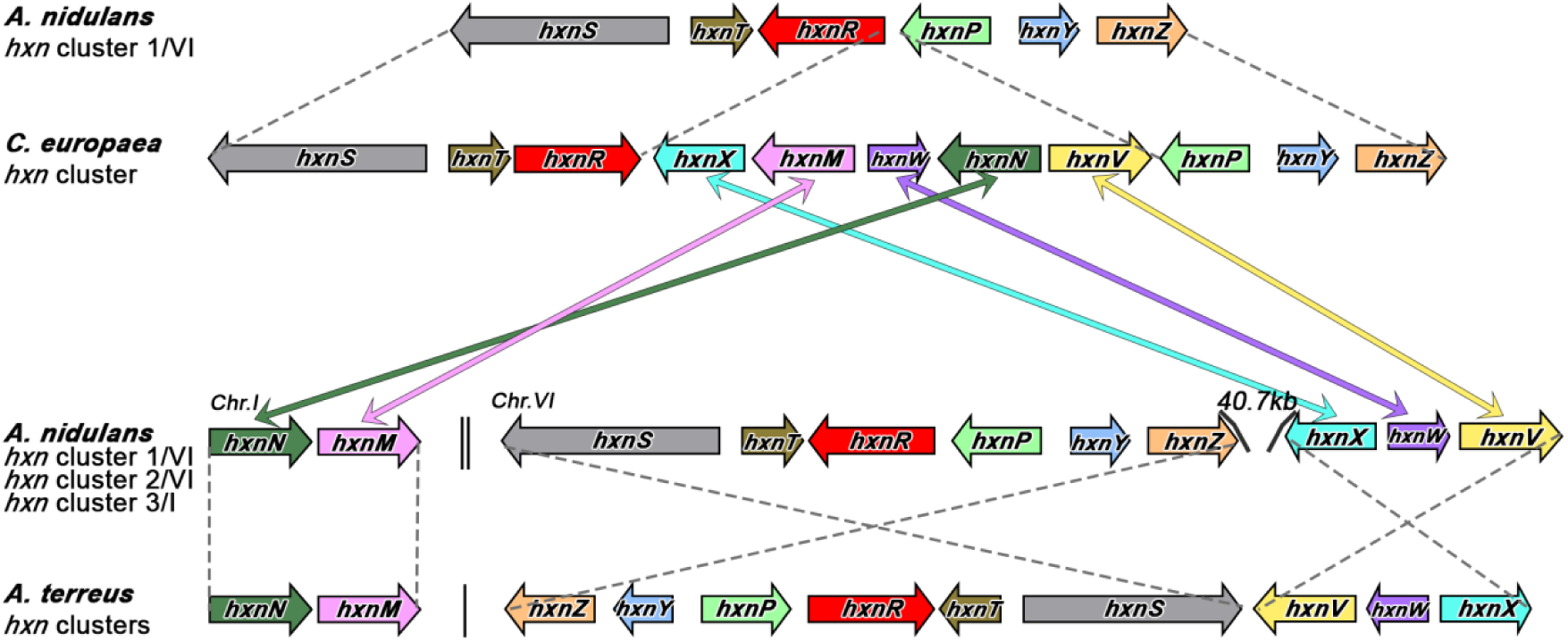
Expanded clusters in *Eurotiomycetes* uncover new *hxn* genes. Comparison of the organisation of known [11] and putative novel *hxn* genes in three species: *A. nidulans*, *A. terreus* and *Cyphellophora europaea.* Each orthologous gene is symbolised by a thick arrow of a different colour, which also indicates relative orientation. Colour-coded double headed arrows connect the 5 new putative *C. europaea hxn* genes to orthologues in the *A. nidulans* genome. Dashed lines connect similarly arranged cluster segments in the three species. For *A. nidulans,* a double vertical line indicates separation of clusters in different chromosomes (Super-scaffold BN001306 for Chromosome VI, BN001301 for Chromosome I). For *A. terreus*, a single vertical line separates two distinct contigs (Contig AAJN01000215 for the 9-gene cluster, AAJN01000156 for the 2-gene cluster). In *C. europaea*, the 11-gene cluster is contained in contig AOBU01000059.

**Table 1.**
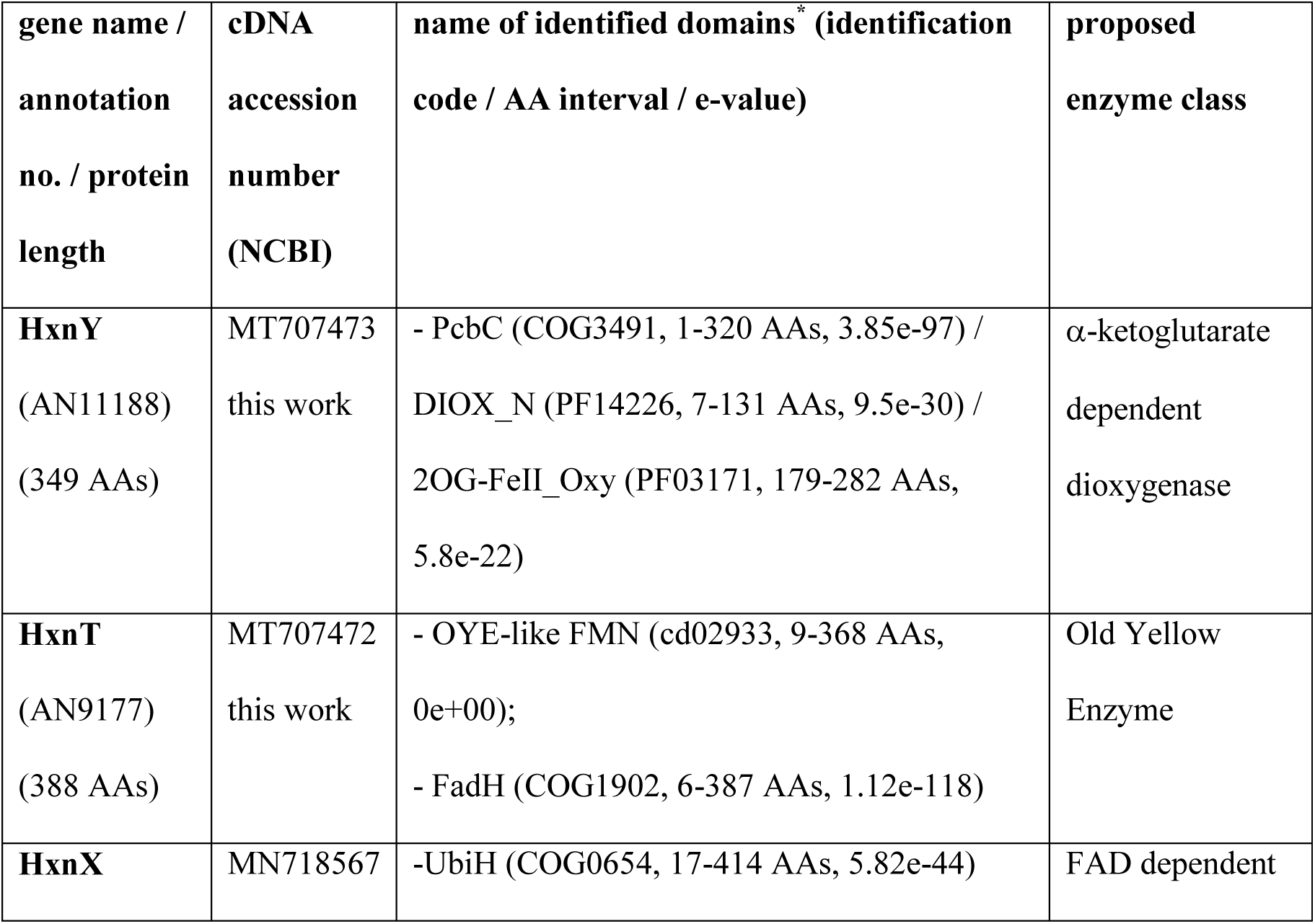
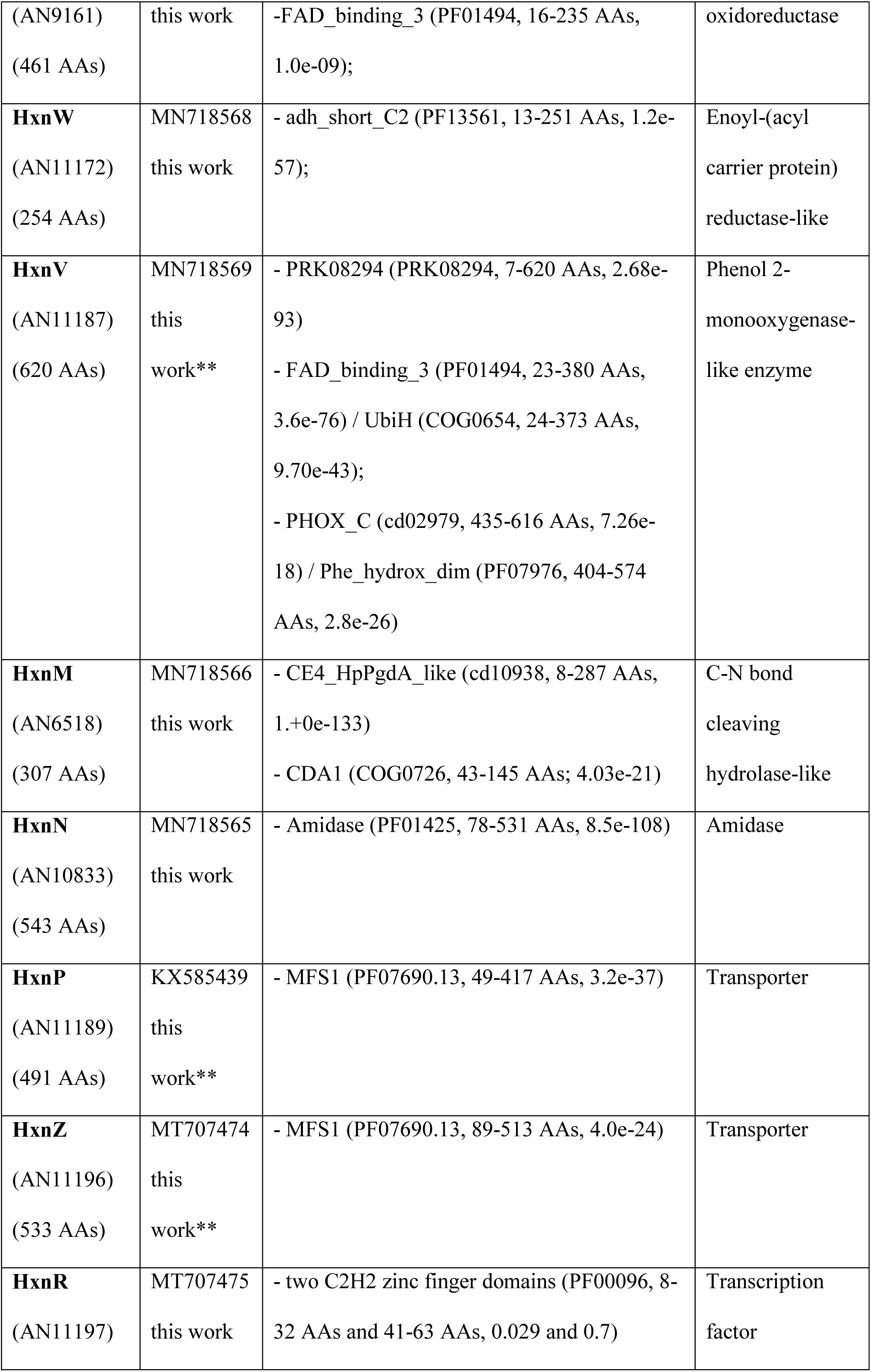
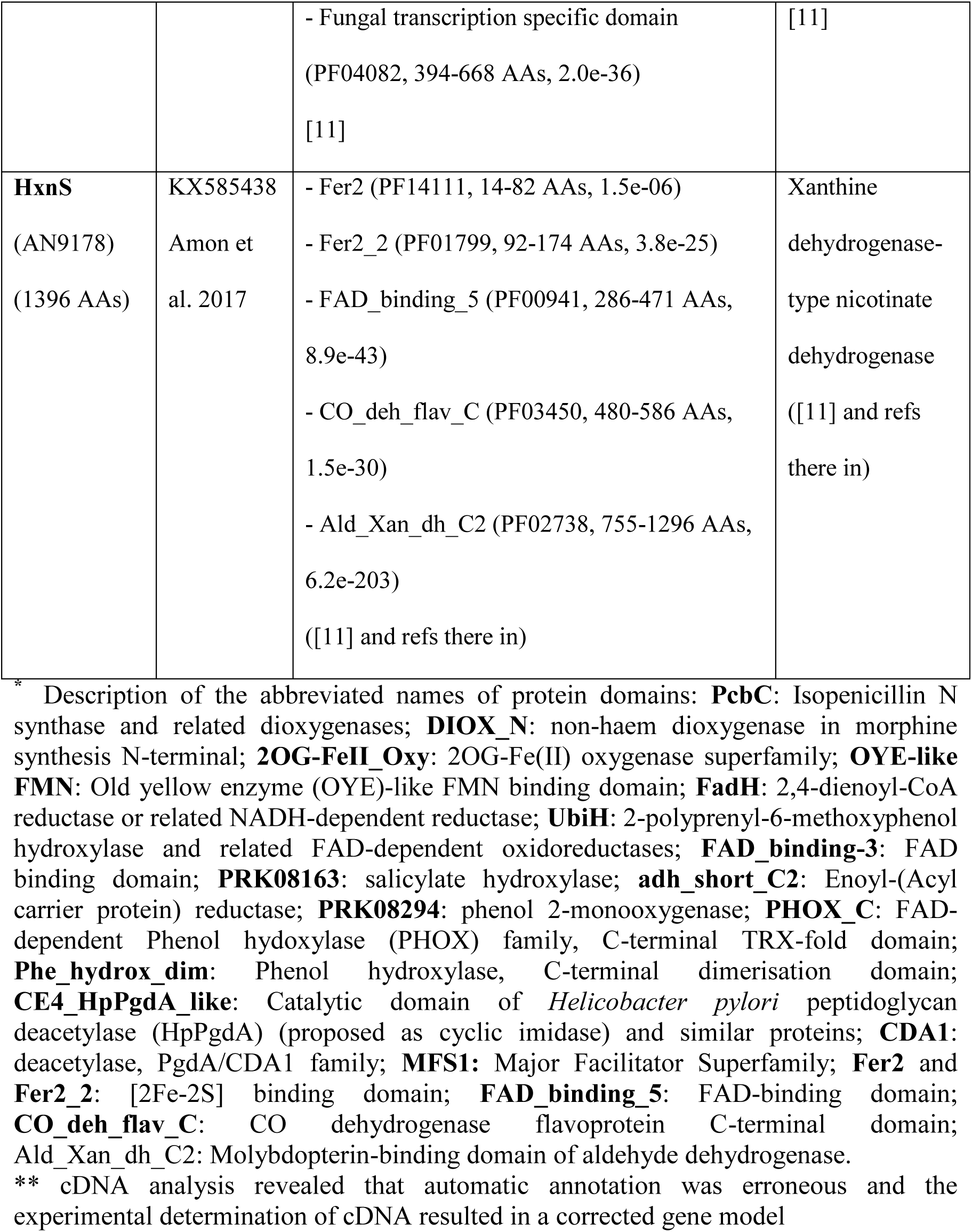
Results of i*n silico* domain analysis of modelled Hxn enzymes

The genomic organisation in Chromosome VI confirms data obtained with a mutagenic screen, which yielded besides mutations in *hxnS* and *hxnR* [11] additional mutants unable to grow on either NA or 6-NA as sole nitrogen sources. A number of tightly linked mutations, of which only two (*hxn6* and *hxn7*) are still available, mapped in chromosome VI at about ≈10 cM from mutations in the *hxnS* and *hxnR* genes, which is coherent with the genomic organisation described above (Kelly and Scazzocchio, personal communication).

We isolated from a genomic DNA library [23] a plasmid able to complement *hxn6* for growth on 6-NA as sole nitrogen source. The 8256 bp insert comprises *hxnV*, *hxnW*, *hxnX* and partial flanking sequences of the AN9159 and AN9162 loci. The *hxn6* mutation is a G1171A transition within the *hxnV* ORF (see below for correction of the *hxnV* gene model in S1 Fig) resulting in W296STOP (amber). Southern blots showed *hxn7* to be a chromosomal aberration (possibly an insertion) interrupting the *hxnV* open reading frame (S2 Fig). In *A. nidulans*, the *hxnX* gene (cluster 2/VI) is at 40,748 bps from *hxnZ* (based on re-assembly data [21], while *hxnN* and *hxnM* are adjacent to each other and transcribed from the same strand in Chromosome I (Cluster 3/I) (Fig 1). We obtained cDNAs of the genes in the three clusters, and confirmed that, as gathered by manual inspection and comparative genomics, the database gene models (proposed by automated annotation) for *hxnP, hxnZ* and *hxnV* are erroneous (S1, S3 and S4 Figs for the correct gene models, Table 1 for accession numbers). HxnX, HxnW, HxnV are oxidoreductases, while HxnM and HxnN are hydrolases. A summary of the predicted activities of all the encoded Hxn proteins are shown in table 1. All the genes in cluster 2/VI and 3/I show an HxnR-dependent induction by 6-NA. In an *hxnR^c^7* strain, the genes in clusters 2/VI and 3/I show variable levels of constitutive expression (Fig 2A), as shown before for cluster 1/VI [11]. The limits of the newly detected clusters are demarcated by the completely different pattern of expression of the flanking genes (loci AN9159 and AN9162 for cluster 2/VI, and loci AN6517 and AN10825 for cluster 3/I; Fig 2B).

**Fig 2.**
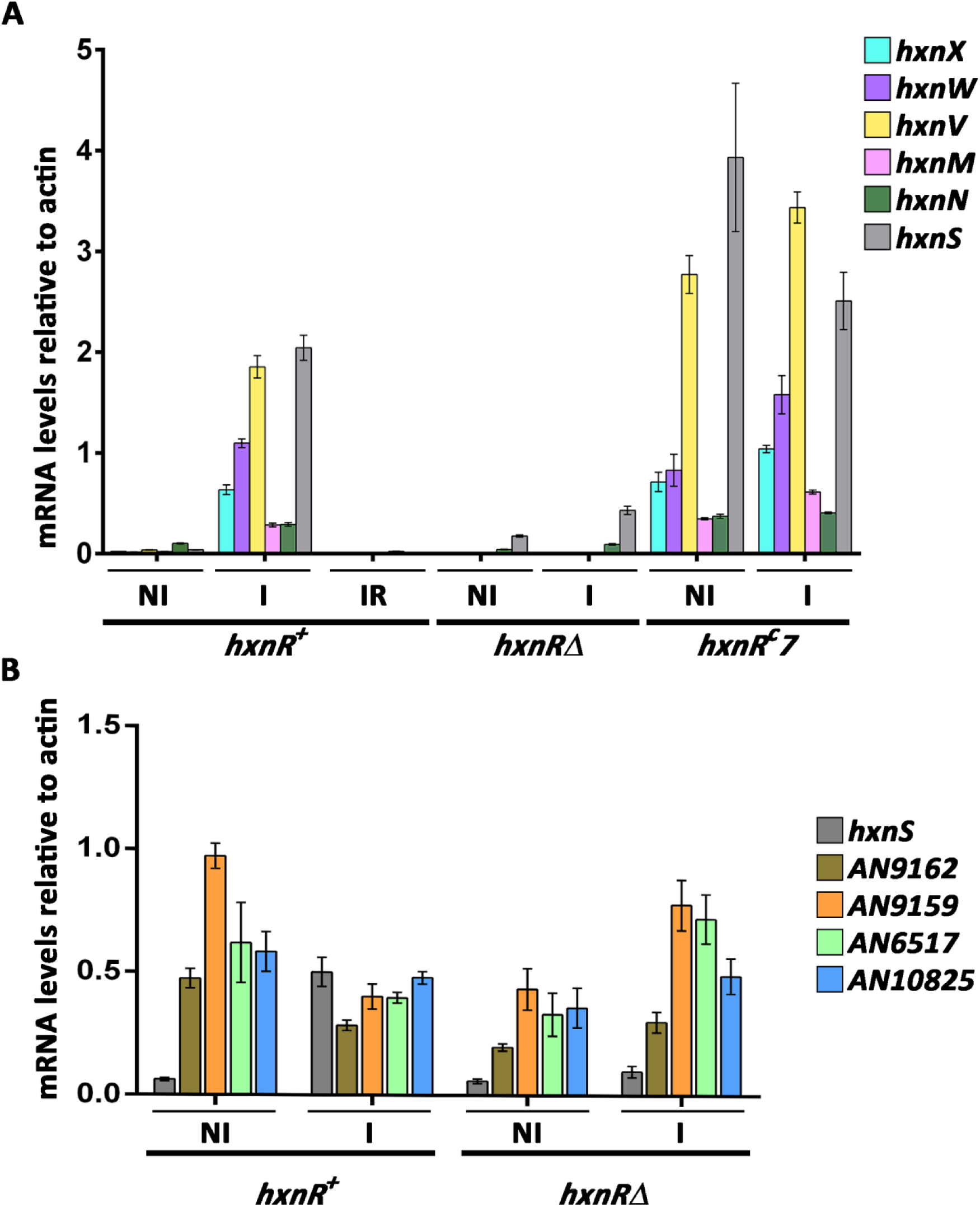
HxnR dependent co-induction by 6-NA and ammonium repression of genes in clusters 2/VI and 3/I. All genes in clusters 2/VI and 3/I (Panel A) and the cognate cluster-flanking genes (Panel B) were tested together with *hxnS* (in cluster VI/1), which was included as a positive control of expression. The relative mRNA levels were measured by RT-qPCR and data were processed according to the relative standard curve method [24] with the γ-actin transcript (*actA/*AN6542) as reference. Mycelia were grown on 10 mM acetamide as sole N-source for 8 h at 37 °C. They were either kept on the same medium for a further 2 h (non-induced, NI) or induced with 1 mM 6-NA (as the sodium salt, I) or induced as above together with 5 mM of L-(+)di-ammonium-tartrate (induced-repressed, IR), also for 2 h. Strains used were *hxnR^+^* (FGSC A26), *hxnR*Δ (HZS.136) and *hxnR^c^7* (FGSC A872) (S1 Table). Standard deviations of three independent experiments are shown. Primers are listed in S2 Table.

As previously shown for the genes in cluster 1/VI, these five newly identified *hxn* genes are strongly ammonium repressible (Fig 2A) and with one exception (see below), strictly dependent on the AreA GATA factor, mediating nitrogen metabolite de-repression (Fig 3).

**Fig 3.**
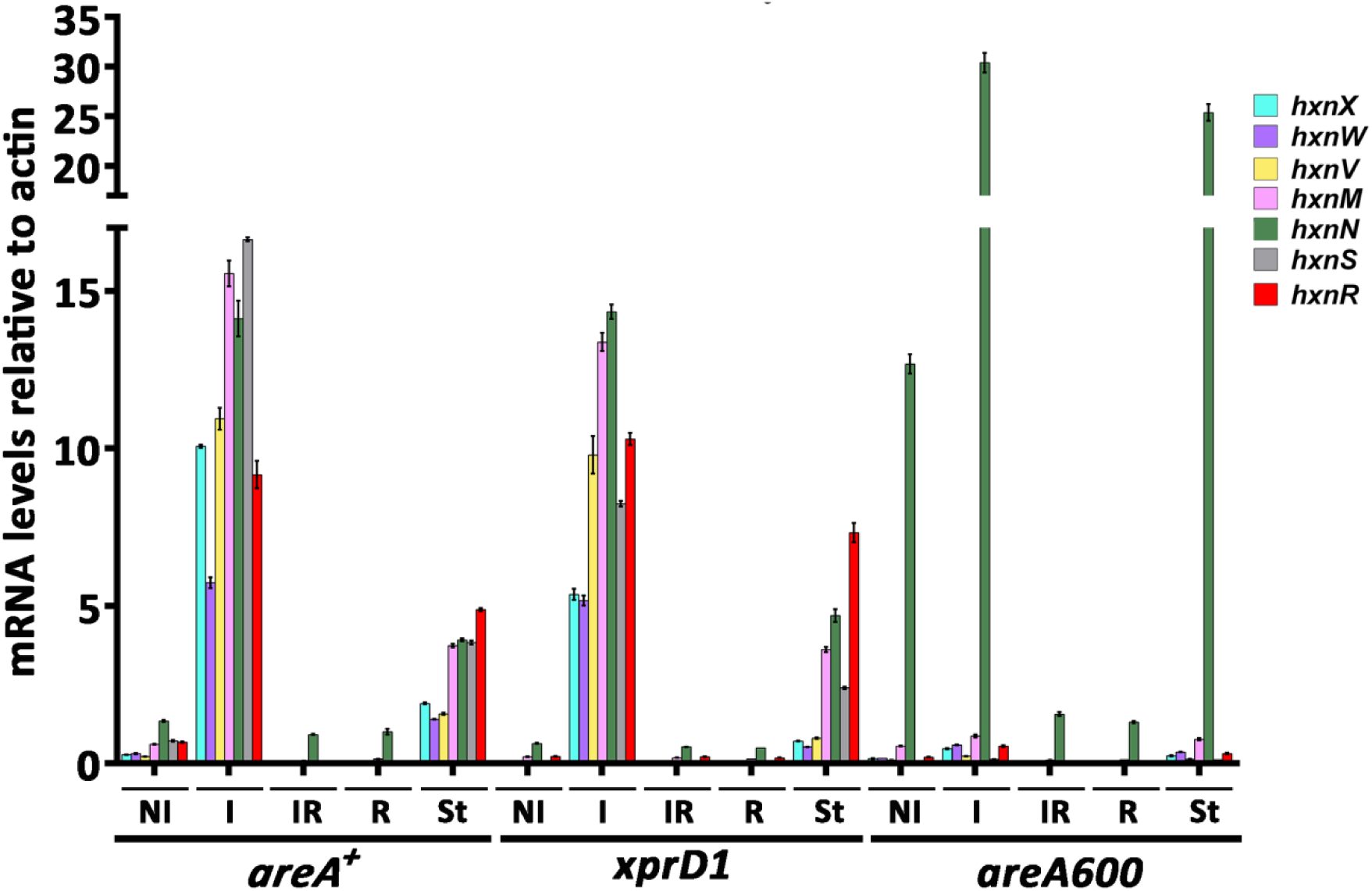
The GATA factor AreA is essential for expression of all *hxn* genes with the exception of *hxnN*. Relative mRNA levels in *areA^+^* (FGSC A26), a supposedly derepressed *areA* mutant (*xprD1*, HZS.216) and an *areA* null mutant (*areA600*, CS3095) strains were determined (S1 Table). Non-induced conditions (NI): Strains were grown on MM media with 5 mM L-(+)di-ammonium-tartrate as sole N-source for 8 h, then the mycelia were transferred to MM with 10 mM acetamide for further 2 h. Induced conditions (I): as above but transferred to 10 mM nicotinic acid as sole N-source. Induced repressed conditions (IR): transferred to 10 mM nicotinic acid and 5 mM L-(+)di-ammonium tartrate. N-starvation conditions (St): transferred to nitrogen-source-free medium. RT-qPCR data were processed according to the standard curve method [24] with the γ-actin transcript (*actA/*AN6542) as reference. Standard deviations based on three biological replicates are shown. Primers are listed in S2 Table.

The *xprD1* is usually considered to be the most extreme de-repressed allele of the *areA* regulatory gene [25], however, it did not behave as a de-repressed allele for the expression of any *hxn* gene but rather as a partial loss of function allele for *hxnS* and *hxnP* expression [11] while being variable in its effects on the genes in clusters 2/VI and 3/I (Fig 3). A similar behaviour was reported for *ureA* (a urea transporter gene) expression [26], which strongly suggests that the phenotypes resulting from this specific mutation are promoter-dependent. The amidase-encoding *hxnN* gene shows a paradoxical pattern of expression. While it is clearly subject to repression by ammonium, it is drastically over-expressed in *areA600* background under neutral (non-induced, non-repressed conditions, see legend to Fig 3), as well as under induced and nitrogen starvation conditions (Fig 3). As *areA600* is a null mutation due to a chain termination mutation upstream of the DNA binding domain [27], we must conclude that AreA does not act as a transcriptional activator but as a repressor for *hxnN*. The apparently paradoxical susceptibility of *hxnN* to ammonium repression is most probably due to its complete dependence on HxnR, whose expression is drastically repressed by ammonium ([11] and Figs 2 and 3). We searched the genes in the three clusters for the consensus AreA 5’HGATAR DNA binding sites [28] (Fig 4). The *hxnV* gene upstream sequence does not feature canonical AreA sites; nevertheless its expression is repressible by ammonium, most likely due to indirect repression via repression of *hxnR* transcription. The *hxnR* upstream region shows both canonical AreA sites and one putative HxnR binding site (see below). This is consistent with this gene being inducible, self-regulated and subject to nitrogen metabolite repression ([11] and Fig 2). The negative effect of AreA on *hxnN* expression may be due to the presence of a canonical GATA binding motif (5’AGATAA on the non-coding strand at position -14 to -19), interfering with the start or progress of transcription.

**Fig 4.**
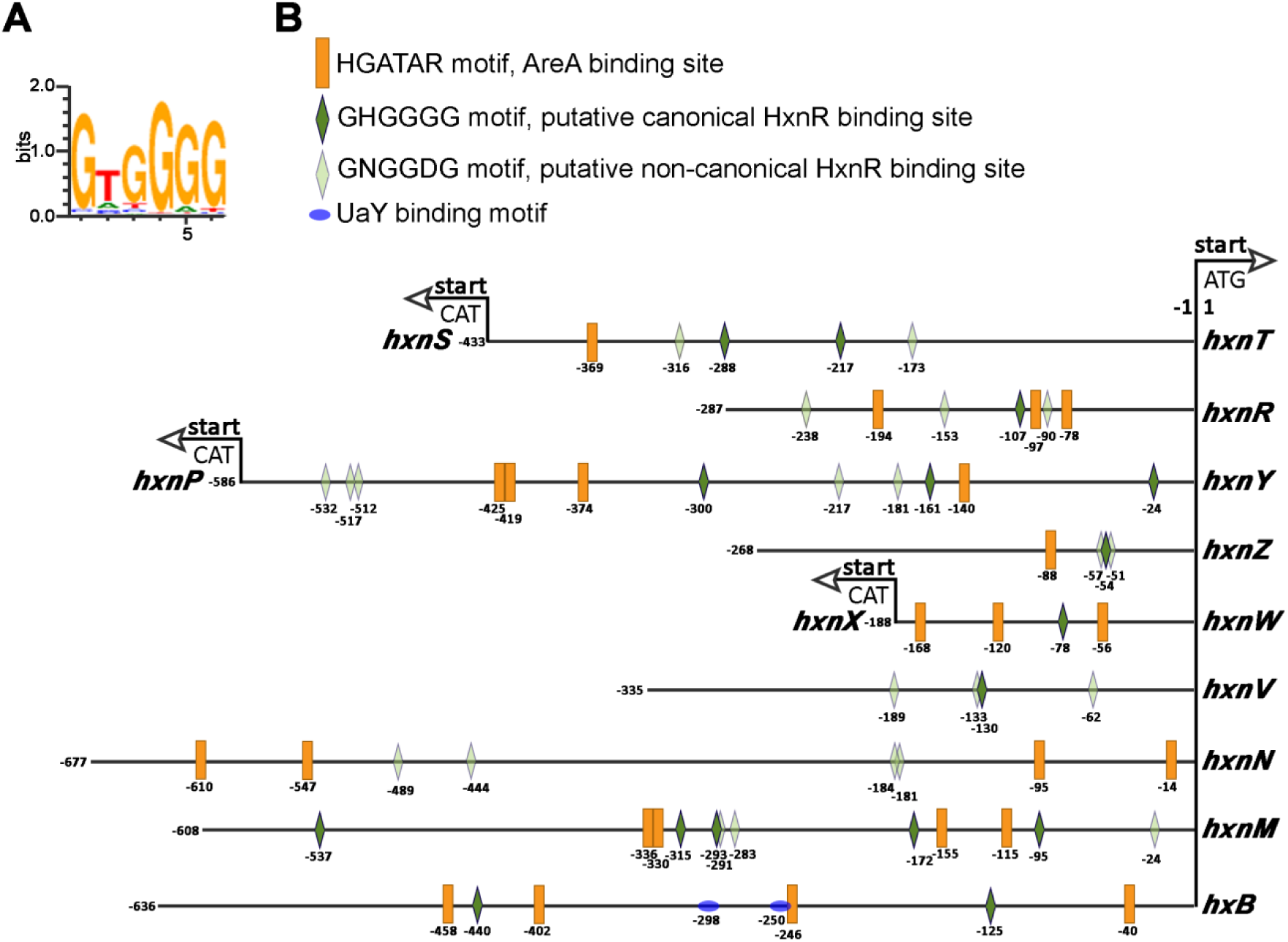
AreA and putative HxnR binding sites are extant in the 11 genes of the *hxn* regulon. (A) Sequence logo of the DNA binding motif of the HxnR transcription factor generated by the “DNA binding site predictor for Cys2His2 Zinc Finger Proteins” application (http://zf.princeton.edu/) [29]. (B) Distribution of 5’HGATAR AreA binding sites (orange boxes) [28] and putative canonical 5’GHGGGG HxnR binding sites (dark green lozenges) in *hxn* gene promoters and also in the promoter of the *hxB* gene. The latter encodes a trans-sulphurylase necessary for the activity of the MOCO cofactor in enzymes of the xanthine oxidoreductase group (including HxnS and HxA). UaY binding sites on the *hxB* promoter are marked by blue coloured ovals [30]. Sequences conforming to the consensus 5’GHGGGG sequence are present in all HxnR-regulated genes, except *hxnN*. Nevertheless, Fig 2 shows clearly that *hxnN* is under the control of HxnR. Thus, the physiological binding sites may have a more relaxed consensus sequence. We propose 5’GNGGDG motif as a non-canonical consensus binding site that can be found in *hxnN* as well as in other *hxn* promoters. Light green lozenges indicate the location of the more relaxed consensus 5’GNGGDG motif. Note that the *hxnT*/*hxnS*, *hxnP*/*hxnY* and *hxnX*/*hxnW* gene couples share bidirectional promoters.

The binding sites of HxnR have not been experimentally determined, however, they could be predicted with reasonable probability [29]. Besides the consensus 5’HGATAR AreA binding sites, Fig 4 shows also the distribution of the putative canonical and non-canonical HxnR binding sites (5’GHGGGG and 5’GNGGDG, respectively) in all 11 *hxn* genes as well as in the *hxB* gene (AN1637), encoding a MOCO sulphurylase ([31] for review) necessary for the enzymatic activity of both HxA and HxnS [30]. Two putative canonical HxnR binding sites are extant in the *hxB* promoter (Fig 4). This gene is under the independent and additive control of UaY (the transcription factor regulating the purine utilisation pathway) and HxnR [30].

### Chromosome rearrangements lead to separation of clusters 1/VI and 2/VI in *A. nidulans* and other Aspergilli

The organisation described above for *A. terreus* (section *Terrei*) is most probably ancestral to the Aspergilli, as is it seen in species belonging to diverging sections of this genus (Fig 5 and S5 Fig) namely in *A. carbonarius* (section *Nigri*) and in *A. unguis,* an early diverging species of section *Nidulantes* (Fig 5 and S5 Fig).

**Fig 5.**
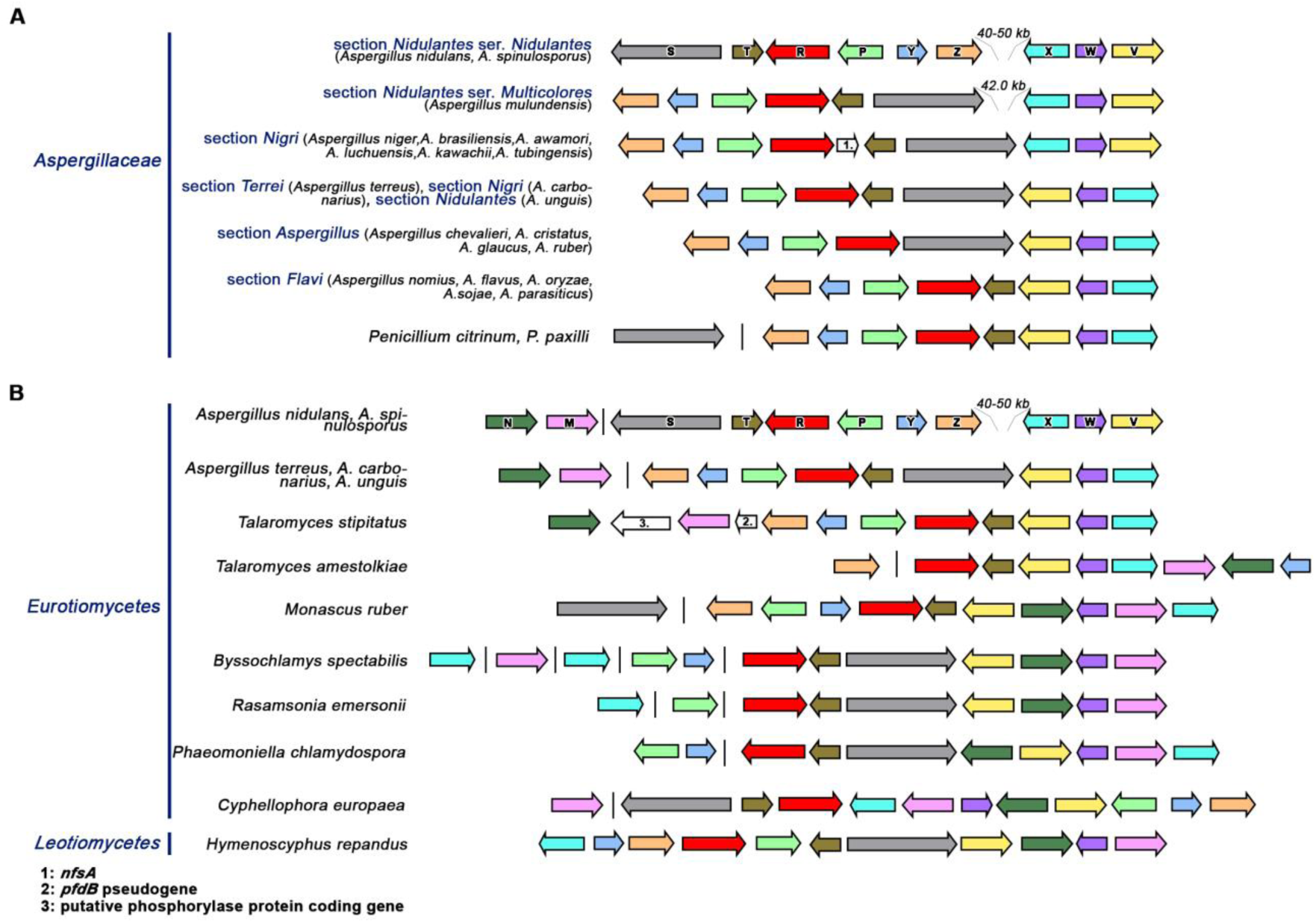
Genomic arrangement of the *hxn* gene clusters. (A) *Aspergillaceae* (B) Other *Eurotiomycetes* compared with *Hymenoscyphus repandus* (*Leotiomycetes*, *Helotiales*). Orthologues found in different species are indicated by arrows of the same colour as in Fig 1. A single vertical line symbolises physical separation of genes on different contigs.

This organisation closely resembles the one seen in species of the basal section *Aspergillus* (*A. ruber*, *A. glaucus*, *A. chevalieri* and *A. cristatus*), where, however, the *hxnT* gene is absent. Within section *Nidulantes*, a first chromosomal inversion resulted in the separation of the clusters 1/VI and 2/VI, inverting the position and orientation of *hxnX, hxnW,* and *hxnV* (in cluster 2/VI) in relation to *hxnS* (in cluster 1/VI). This results in *A. mulundensis* in a gap of 41,992 bp between the two clusters (shown in Fig 5, section *Nidulantes*, series *Multicolores* [32]). A second inversion within this taxon led to the configuration seen in *A. nidulans* and its closest sequenced relative *A. spinulosporus* (section *Nidulantes*, series *Nidulantes*) [32] leaving respectively gaps of 40,876 bp and 49,972 bp between *hxnZ* and *hxnX. A. sydowii* and *A. versicolor* (section *Nidulantes*, series *Versicolores*) [32] (S5A Fig), also show separation of clusters 1/VI and 2/VI, however, the relative gene orientation and phylogenetic position of the latter two species strongly suggest that this organisation arose from events independent to those described above for *A. nidulans* and *A. spinulosporus*. Two distinct independent inversions, like the one described above for *A. mulundensis* must have occurred within section *Nigri,* leading to the organisation seen in *A. aculeatus* and the *A. niger* clade (Fig 5 and S5A Fig); in *A*. *niger* and allied species, *hxnS* and *hxnX* are abutting neighbours; in *A. aculeatus* (also in section *Nigri*), where *hxnT* is absent there is a ∼32 kb gap between these genes.

In two species (*A. steynii* and *A. westerdijkiae*) of two closely related series (ser. *Steyniorum* and ser. *Circumdati*, respectively), clusters 1 and 2 are separated without any relative change of gene orientation (S5A Fig). This could be formally described as an insertion, however partial DNA identity and gene synteny in the inter-cluster sequence rather suggest two successive inversions. In *A. wentii* (section *Cremei*) a rearrangement associated with the loss of *hxnS* separates from the original cluster, a sub-cluster including *hxnZ*, *hxnY*, and a pseudogenised *hxnP*; while *hxnV*, *hxnW* and *hxnX* are still included in the main cluster together with the neighbouring *hxnT* and *hxnR*.

### In the *Pezizomycotina*, with the exception of the Aspergilli, the *hxnN* and *hxnM* genes are included in the *hxn* cluster

The enzymes encoded in clusters 1/VI and 2/VI are all oxidoreductase enzymes, however to release ammonium from NA-derived metabolites, hydrolytic enzymes are necessary [2]. Within the putative *hxn* clusters of many *Pezizomycotina* species, two genes encoding respectively a putative cyclic-imide hydrolase (*hxnM* EC 3.5.2.16, >60% identity with AAY98498, the cyclic imide hydrolase from *Pseudomonas putida*, [33]) and a putative amidase (*hxnN* EC 3.5.1.4) are extant. The cognate genes of *A. nidulans* have been described above. In *C. europaea, hxnN* and *hxnM* lie in between *hxnX* and *hxnV*, and are separated by *hxnW* constituting two neighbouring, divergently transcribed gene couples, *hxnV*-*hxnN* and *hxnW-hxnM* within the cluster (Fig 1). It should be stressed that these two divergently transcribed couples are conserved across different classes of the *Pezizomycotina* (S5B Fig), however, not in the genus *Aspergillus*, where with the exception of section *Flavi*, *hxnN* and *hxnM* are separated from the main cluster.

In *Monascus ruber*, (*Eurotiomycetes*, *Eurotiales*, *Aspergillaceae* - same family as *Aspergillus*) where *hxnS* is not included in a 10-strong gene cluster (see below), the two divergently transcribed couples are conserved. In *Talaromyces stipitatus* and *T. islandicus* (*Eurotiales*, *Trichocomaceae*), where *hxnS* is altogether missing) *hxnM* and *hxnN* are at one terminus of the 10-strong gene cluster. Fig 5 and S5B Fig show a variety of cluster organisations in species of the *Pezizomycotina* and *Saccharomycotina* subphyla, with the *hxnN* and *hxnM* genes showing different patterns of integration within the *hxn* cluster, with however a remarkable conservation of the <*hxnN-hxnV*> and <*hxnM-hxnW*> divergently transcribed couples in classes of *Pezizomycotina*.

In the genome of *C. europaea*, besides the divergently transcribed couples mentioned above, two other couples are extant, <*hxnS-hxnT*> and <*hxnP-hxnY*>. These couples are mostly conserved in the *Pezizomycotina*, irrespective of whether all eleven genes are included in a single cluster. Noticeably, in *A. nidulans*, cluster 1/VI comprises <*hxnS-hxnT*> and <*hxnP-hxnY*>. In *Hymenoscyphus repandus* (*Leotiomycetes*, *Helotiales*), similarly to *C. europaea*, all 11 genes are included in a single mega-cluster, albeit in a different arrangement; nevertheless, two divergent couples are conserved (<*hxnS-hxnT*> and <*hxnM-hxnW*>). A similar conservation of divergently transcribed genes is seen in other gene clusters, such as the DAL cluster of the *Saccharomycetales*, where the <*DAL4-DAL1*> pair is conserved between *S. cerevisiae* and *Naumovia castellii* in spite of two inversions affecting the budding yeast DAL cluster in chromosome IX [34], and in the biotin biosynthesis cluster of the *Pezizomycotina* (<*bioF-bioDA*> [35]. The persistence of these divergently transcribed couples could be due to the fact that they share a bi-directional promoter, as established for *GAL10* and *GAL1* in *S. cerevisiae* ([36, 37] and refs therein) and for *niiA* - *niaD* in *A. nidulans* [38, 39].

### Evolution of the *hxn* gene cluster(s) in the *Ascomycetes*

Previous work has shown that HxnS is restricted to the *Pezizomycotina* [11]. Thus, it is unlikely that other fungi could hydroxylate NA and thus utilise it as a nitrogen source. However, it is possible that an *hxnS* gene was incorporated into a pre-existent metabolic pathway, whether catabolic or detoxifying, whether or not organised as a cluster. We thus investigated the presence of putative *hxn* clustered genes throughout the fungal kingdom. No putative *hxn* clusters are present in any early divergent fungal lineages in the *Basidiomycota* or in the *Taphrynomycotina,* except that *hxnT*, *hxnN* and *hxnM* unlinked orthologues are present in the early diverging *Taphrinomycotina*, *Saitoella complicata* (see S5 Fig).

Clusters comprising *hxn* genes are present in several scattered species of *Saccharomycotina* (S5B Fig), however not in the *Saccharomycetaceae* and *Debaryomycetaceae* families. All species of *Lipomyces,* an early divergent genus of the *Saccahromycotina*, include *hxnN* and *hxnM* genes (see above). The genomes of fourteen scattered species of *Saccharomycotina* (S5B Fig) comprise clusters with the *hxn* gene complement, always including the transcription factor *hxnR* and never including *hxnS, hxnZ,* and *hxnN*, even if the latter gene could be found unlinked to the cluster in an early divergent species (*Trigonopsis variabilis*). A phylogeny of *hxnR* is shown in supplementary S6 Fig and is consistent with a monophyletic origin of this gene in the *Saccharomycotina* and *Pezizomycotina*. It seems most unlikely that the clusters of the *Saccharomycotina* have a single origin. The *Lipomyces hxnM*-*hxnN* divergent gene pair is found only in this genus from where all other *hxn* genes are absent. Among other families, the occurrence of clusters with variable organisations does not follow any obvious evolutionary pattern. In the fourteen species of *Saccharomycotina* where we found an *hxn* cluster, the *hxnT*, *hxnR* and *hxnV* genes, are monophyletic (S5-S8 Figs). Notwithstanding the above, the phylogeny of *hxnM* suggests several different origins of clustered *hxnM*s within the *Saccharomycetales* from an un-clustered paralogue, possibly acquired by HGT (see below, and S9 Fig). One clustering event occurred in the *Phaffomycetaceae*, possibly two in the *Pichiaceae*, while only one species of the CUG-Ala clade, *Pachysolen tannophilus* [40] includes an *hxn* cluster, with an *hxnM* gene. Among the *Pichiaceae*, in the genus *Ogatea*, the monophyletic origin of clustered and un-clustered *hxnM* genes is supported by their intron exon organisation (S9 Fig).

Several instances of gene loss, gene duplication and cluster reorganisation have occurred in the *Pezizomycotina*. In some *Aspergillus* species, *hxnT* (encoding an FMN dependent oxidoreductase) is missing from the cluster (S5A Fig) and indeed from the genome. In many taxa of *Sordariomycetes* duplication of *hxnV* and subsequent loss of the *hxn* cluster genes can be observed, leaving just the *hxnV* copy and *hxnM*.

It is striking that in the *Aspergillus* section *Flavi,* in *Talaromyces* species and in most species of *Penicillium* the *hxnS* gene is absent and the organisation of the whole cluster is completely identical in some species of *Talaromyces*, in most of Penicillia and in *Aspergillus* section *Flavi* (S5 Fig). This coincidence indicates possible HGTs between these taxons (see below, HGT between *Talaromyces* and *Aspergillus* section *Flavi*). As the transcription factor-encoding gene *hxnR* is conserved, the implication is that these organisms should be able to utilise 6-NA but not NA.

### Insertion of additional genes within the *hxn* clusters

We define as “additional genes” those that appear sporadically within the *hxn* clusters of some taxa. The insertion of a gene encoding a nitro reductase (*nfsA*) originally horizontally transmitted from a cyanobacterium has been discussed previously [11]; the insertion occurred after the divergence of *A. carbonarius* from other members of section *Nigri* [41] (Fig 5 and S5A Fig).

In the *hxn* cluster of *Aspergillu*s section *Flavi*, and in a number of *Penicillium* and *Talaromyces* species (S5, S10 and S11 Figs), a gene of unknown function, to be called *pfdB,* for putative **<u>p</u>**eroxisomal **<u>F</u>**MN-dependent **<u>d</u>**ehydrogenase (see below) lies between *hxnZ* and *hxnM*. This is a paralogue of *pfdA*, a gene universally present in the *Pezizomycotina*, which is never included in an *hxn* cluster. The encoded proteins include PF01070.18 (FMN-dependent dehydrogenase) and PF00173.28 (Cytochrome b5-like binding domain) domains and have a canonical PST1 (peroxisomal entry signal [42]. The phylogeny of PfdA and PfdB clearly suppports a scenario of gene duplication of *pfdA* in the ancestor of Penicillia with simultaneous of subsequent cluster integration (mean similarity between *A* and *B* paralogues 65% compared with 88% of *A* orthologues among themselves) (S11 Fig). *pfdA* has a second, distinct paralogue, *pfdC*, too, which however lost the PST1 signal in some cases and are only present in section *Flavi,* and in a number of *Talaromyces* and *Penicilium* species and in a few species of other clades (S10 and S11 Figs). The occurrence of PfdC in taxons is consistent with the duplication of the PfdA ancestor in an early diverging species followed by several episodes of loss completely unrelated to the evolution of the *hxn* cluster.

In *P. paxilli*, *P. citrinum* and *P. steckii,* a gene encoding a protein of 467-469 residues, comprising a PF00781.24, diacylglycerol kinase catalytic domain, (orthologues annotated as sphingoid long chain kinases) lies between the *hxnZ* and *hxnM* genes. This gene is duplication of a gene present elsewhere in these organisms and omnipresent in the *Eurotiomycetes*. In *Talaromyces stipitatus* a *pfdB* pseudogene is extant between *hxnZ* and *hxnM*, and additionally, an intron-less gene encoding 751 residue-multidomain protein, comprising an N terminal PF0104820.11 (phosphorylase superfamily N-terminal, most similar to nucleoside phophorylases) domain and a C-terminal PF05960.11 (bacterial protein of unknown function) domain is located between *hxnN* and *hxnM*, the nearest homologues of the inserted gene being present and unlinked to any *hxn* gene in *T. verruculosus*.

In *Kregervanrija fluxuum* (*Saccahromycotina*, *Pichiaceae*) a putative amidase gene is inserted in the cluster between *hxnM* and *hxnT* (S5B Fig). The encoded protein has only 35% identity with *hxnN* of *A. nidulans*, compared with the 51% identity shown by the genuine HxnN proteins of *Lipomyces starkei, Trigonopsis variabilis* and *Saitoella complicata*. Its nearest homologue is a putative amidase from *Ogatea parapolymorpha* (56% identity). It is tempting to speculate that this amidase has been recruited to the cluster to carry out a similar catalytical function to that afforded by HxnN.

### HGT events involving *hxn* genes

In most Penicillia and *Talaromyces* species the *hxnS* gene is absent. The un-clustered *hxnS* genes of *T. islandicus*, *T. piceae*, and *T. wortmanii* have as sister clade the *hxnS* of *Monascus* species, consistent with standard phylogeny. The phylogeny [11] (S5 and S12 Figs) together with *hxnS* sequence identity strongly suggests episodes of *hxnS* de-clustering for these three species. A different situation occurs in *P. citrinum*, *P. paxilli* and *P. steckii*. These sister species (Section *Citrina*, [43]) have reacquired an un-clustered *hxnS* gene by HGT from either a *Fusarium* or a *Colletotrichum* species (both are *Sordariomycetes*, S12 Fig) [11].

In all investigated dikarya, HxnM paralogues, presumably non-related to NA metabolism are extant. Based on comprehensive phylogeny of *hxnM* and its paralogues (S9 Fig) subjected to reconciliation with the species tree (using GeneRax), we confirmed HGTs amongst *Ascomycota* taxons and HGT from *Ascomycota* to *Basidiomycota*. *Dothideomycetes* acquired clustered *hxnM* from *Symbiotaphrina* (*Xylonomycetes*). Additionally, within the large clade of unclustered *hxnM* genes of the *Pezizomycotina* a group containing *P. brasillianum* was found as the donor of *hxnM* to *Aspergillus* section *Usti*. The common ancestor of the *Panellus stipticus* and *Mycena galopus* (*Basidiomycota*) acquired *hxnM* from the common ancestor of the Fusaria (*Ascomycota*). Since these two *Basidiomycetes* have only a single, Fusaria-derived *hxnM* gene, the *Basidiomycota hxnM* must necessarily be lost from these species.

S9 Fig is consistent with a vertical inheritance of *hxnM* homologues in the dikarya, excluding a recent HGT from bacteria. The phylogeny of HxnM is compatible with an originally un-clustered *hxnM* homologue being duplicated, one copy being recruited in an *hxn* cluster. Within the *Pezizomycotina*, there are two clades including un-clustered *hxnM* homologues, one (shown by light green highlighting in S9 Fig) is basal to all the *hxnM* clustered genes in both *Pezizomycotina* and *Saccharomycotina*. In the early diverging *Lipomyces* (*Lipomycetaceae*, *Saccharomycotina*) genus, *hxnM* and *hxnN* are clustered and divergently transcribed, and no other putative *hxn* genes are extant.

While the clustered *hxnM* genes appear monophyletic, originating from the same clade of un-clustered genes, clustering in the *Pezizomycotina* occurred independently from that within the *Saccharomycotina*, followed by several independent instances of separation of an *hxnN*-*hxnM* minicluster (such as detailed above for the *Aspergilli*) and presence of an *hxnM* un-clustered homologue, as it occurred in the *Leotiomycetes*.

The clade comprising the HxnM homologues of the *Saccharomycotina* seems monophyletic. However, it does not occur as expected as a sister clade of all the homologues of the *Pezizomycotina*, but within the different *Pezizomycotina* clades. The low aLRT value at the relevant node, however, does not support *Saccharomycotina* acquiring an *hxnM* gene by HGT from *Pezizomycotina*.

Reconciliation of the phylogeny of the nicotinate catabolism non-related PfdB found in *Eurotiomycetes* with the species tree (by using GeneRax) confirmed that the *pfdB* of *Talaromyces* which was acquired by HGT from an ancestral species of Penicillia was further transferred from a *Talaromyces* by HGT to an ancestor of *Aspergillus* section *Flavi* (Fig 6). Since all Penicillia and *Aspergillus* section *Flavi* share an identical cluster organisation with some species of *Talaromyces*, the HGT events probably involved two episodes of HGT of the whole *hxn* cluster. This outlines a scenario by which, after the appearance of *pfdB* by a single gene duplication of *pfdA* in the ancestral species of Penicillia, *pfdB* subsequently integrated into the cluster in this genus. An HGT of the whole cluster to an early diverging species of *Talaromyces* would have occurred followed by a further HGT from *Talaromyces* to the ancestor of *Aspergillus* section *Flavi*. This scenario implies that the putative acceptor ancestor *Aspergillus* must have lost previously the cluster present in other Aspergilli. This is strikingly confirmed by genomes of early diverging species of section *Flavi* (*A. leporis*, *A. aliaceus*, *A. albertensis* and *A. bertholletius*), which show both instances of *hxn* gene loss and presence of *hxn* pseudogenes (S5A Fig). The most extreme case being that of *A. coremiiformis*, where no *hxn* genes are present. In *A. bertholletius* a cluster of 7 *hxn* pseudogenes is extant, where the only intact gene is *hxnT* (S5A Fig), however this gene is not a fossil, but it derives from *Talaromyces* by HGT (S7 Fig). The earliest diverged species of section *Flavi* is supposed to be *A. avenaceus* [44, 45]. This is fully supported by the position of the cluster-independent *pfdA* and *pfdC* genes in the phylogenetic tree (S10 Fig). The cluster of this species, which includes *pfdB,* is similar to that of other *Flavi*, except that *hxnP* is missing and neither of the two *hxnM* paralogues is included in the cluster.

**Fig 6.**
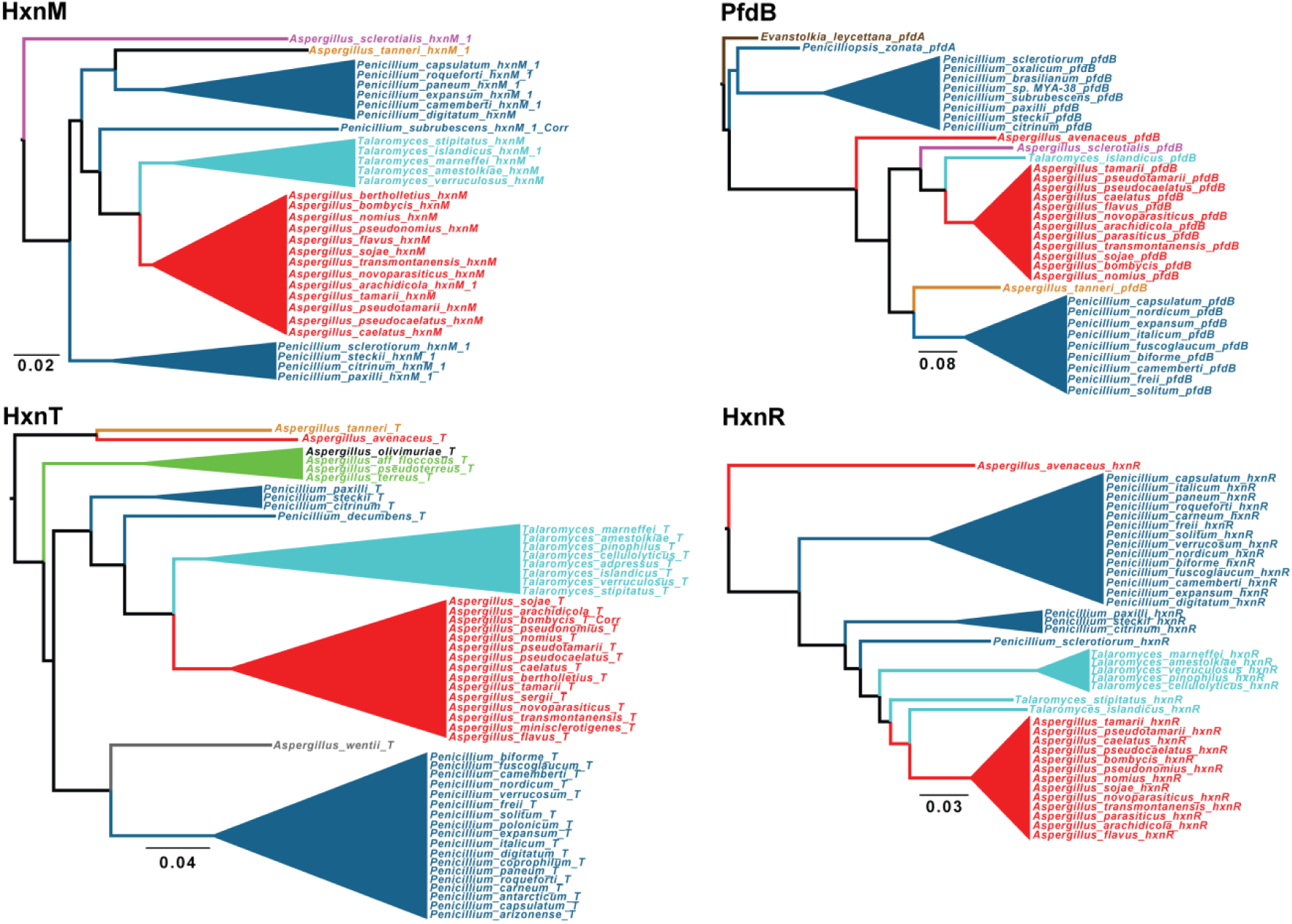
HGT from *Talaromyces* to *Aspergillus* section *Flavi* is supported by the phylogenies of four different proteins based on *Eurotiomycetes* data set (see S6, S7, S9 and S10 Figs. for the complete phylogenies). Cyan: *Talaromyces*; Blue: *Penicillium*; Red: *Aspergillus* section *Flavi*; Purple: *Aspergillus* section *Polypaecilum*; Light brown: *Aspergillus* section *Tannerorum*; Green: *Aspergillus* section *Terrei*; Gray: *Aspergillus* section *Cremei*; Black: *Aspergillus* section *Flavipedes*.

Reconciliation analysis of HxnT, HxnR and HxnM phylogeny restricted to *Eurotiomycetes* confirmed the HGT between *Talaromyces* and section *Flavi* only in the case of HxnR, despite the layout of the phylogenetic trees of HxnM and HxnT were basically the same as in case of PfdB and HxnR (Fig 6). In spite of contradictory results, the evidence strongly suggests the whole HG transfer of the cluster as detailed above.

Disturbingly, in the *hxnR*, *hxnV* and *hxnT* phylogenies, *A. avenaceus* appears as out-species of the *Talaromyces*/*Penicillium* clade which transferred the cluster to other *Flavi* (Fig 6 and S6-S8 figs.). There is obviously a complex series of HGTs which may be solved when more genomes of closely related species become available.

*A. sclerotialis* (subgenus *Polypaecilum*) has an *hxn* cluster of complex origin. Supported by phylogeny of HxnS, HxnR, HxnV and HxnT, the corresponding genes derived by HGT from a *Fusarium* (*Nectria*) species (S6-S8 and S12 Figs). Its cluster includes a *pfdB* gene, necessarily derived from a *Talaromyces/Penicillium* species. Its clustered *hxnM1* is sister to the *hxnM1* of *A. tanneri* (see below) (Fig 6, S9 Figs). The most parsimonious hypothesis is that the complex *hxn* cluster of this species originated in the confluence of two HGT events, one from a *Fusarium*/*Nectria* species and another from a *Talaromyces/Penicillium* species, together with an extensive rearrangement of the cluster, leading to a unique pattern of gene organisation.

*Aspergillus tanneri* belongs to section *Tannerorum*, a sister clade to section *Circumdati* [32, 43]. In this species two clusters and an isolated *hxnM-hxnN* pair are extant (S5 Fig). Five enzyme-encoding genes are present in two copies, *hxnV*, *hxnX* and *hxnM*, *hxnS* and *hxnW*. No *hxnZ, hxnY* and *hxnP* genes are extant in *A. tanneri*. The *hxnS* gene present in both clusters is a product of a recent duplication (S12 Fig). In the larger cluster (7 genes), *pfdB* originates from an HGT from *Talaromyces* or *Penicillium* (Fig 6), most likely independently from the HGT to section *Flavi*, *hxnV1, hxnM1* and *hxnT* genes, sister to those extant in *A. avenaceus,* are likely of the same origin, while *hxnR* has not originated from an HGT event (S6-S10 Figs). This is a composite cluster, where HGT events are coalesced with vertically inherited genes. The smaller cluster includes an *hxnV2* paralogue beside an *hxnS* paralogue that also originated from a recent duplication together with an *hxnW* pseudogene (S8 and S12 Figs).

### Concluding remarks

Experimental work has shown that three gene clusters in *A. nidulans* constitute a nicotinate (actually a nicotinate derivative) inducible regulon, under the control of a specific Zn-finger transcription factor, HxnR. Deletion of HxnR have shown that expression of some or all of the genes in this regulon are necessary for NA, 6-NA and the putative intermediate 2,5-dihydroxypyridine utilisation as nitrogen sources [11]. The specific metabolic function of each encoded protein will be reported separately. This regulon is extant only in the *Ascomycetes*. The variable organisation seen in different species includes instances of complete clustering of all 11 genes, which may suggest an evolutionary pressure towards the global integration of the *hxn* genes, together with instances of de-clustering such as the separation of clusters 1(VI) and 2(VI) in section *Nidulantes* of the Aspergilli. Several instances of HGT were detected (Fig 7), most notably the origin of the cluster of *Aspergillus* section *Flavi* from *Talaromyces*/Penicillia. The events of HGT, together with the recruitment of genes after duplication, including *hxnS* and *hxnM*, and additional genes such as *pfdB*, underlies both the dynamic nature and the reticulate character of metabolic cluster evolution.

**Fig 7.**
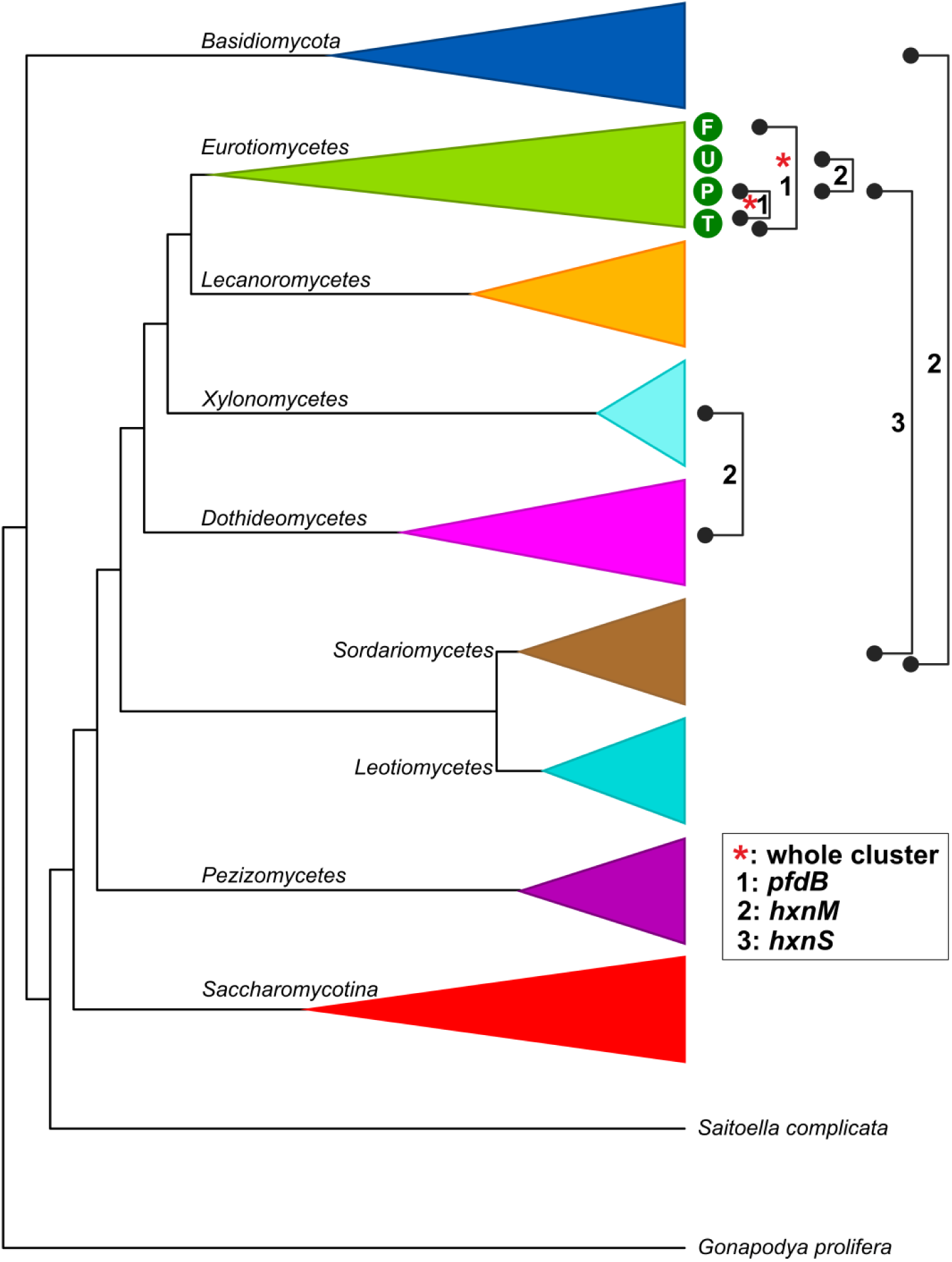
Summary of supported HGT events on collapsed species tree related to *pfdB*, *hxnM* and *hxnS* genes. F: *Aspergillus* section *Flavi*; U: *Aspergillus* section *Usti*; P: Penicillia; T: early diverging species of *Talaromyces*; 1: HGT of *pfdB* gene found between Penicillia and species of *Talaromyces* and between species of *Talaromyces* and *Aspergillus* section *Flavi*; 2: HGTs of *hxnM* gene between *Xylonomycetes* and *Dothideomycetes*, Fusaria (*Sordariomycetes*) and *Basidiomycota* and between a group of Penicillia containing *P. brasilianum* and *Aspergillus* section *Usti*; 3: HGT of *hxnS* gene between *Aspergillus* section *Usti* and Penicillia ; red asterisk: transfer of the whole *hxn* cluster composed of nine *hxn* genes and including the *pfdB* gene from Penicillia to species of *Talaromyces* and from *Talaromyces* to *Aspergillus* section *Flavi*. Solid lines mark confirmed HGTs.

## Materials and Methods

### Strains and growth conditions

The *A. nidulans* strains used in this work are listed in S1 Table. Standard genetic markers are described in http://www.fgsc.net/Aspergillus/gene_list/. Minimal media (MM) contained glucose as the carbon source; the nitrogen source varied according to the experimental condition [11]. The media were supplemented according to the requirements of each auxotrophic strain (www.fgsc.net). Nitrogen sources, inducers and repressors were used at the following concentrations: 10 mM acetamide, 10 mM nicotinic acid (1:100 dilution from 1 M nicotinic acid dissolved in 1 M sodium hydroxide) and 5 mM L-(+)di-ammonium-tartrate as sole N-sources; 1 mM 6-hydroxynicotinic acid sodium salt as inducer and 5 mM L-(+)di-ammonium-tartrate as repressor. Growth conditions are detailed in the figure legends of corresponding experiments.

### RNA manipulation

Total RNA was isolated using a NucleoSpin RNA Plant Kit (Macherey-Nagel) and RNase-Free DNase (Qiagen) according to the manufacturer’s instructions. cDNA synthesis was carried out with a mixture of oligo-dT and random primers using a RevertAid First Strand cDNA Synthesis Kit (Fermentas). Quantitative RT-PCR (RT-qPCR) were carried out in a CFX96 Real Time PCR System (BioRad) with SYBR Green/Fluorescein qPCR Master Mix (Fermentas) reaction mixture (94 °C 3 min followed by 40 cycles of 94 °C 15 s and 60 °C 1 min). Data processing was done by the standard curve method [24]. DNA sequencing was done by the Sanger sequencing service of LGC (http://www.lgcgroup.com). Primers used are listed in the S2 Table.

### Data mining

The coding sequences of fungal *hxn* genes (ATG-STOP) were mined by TBLASTN screening of DNA databases at the NCBI servers, mainly the Whole Genome Shotgun contigs (WGS) database, using the available on-line tools [46]. For a few species (*N. crassa*, *P. anserina*, *P. chrysogenum*, *A. oryzae*, *A. niger* ATCC 1015, *Leptosphaeria maculans* and some *Saccharomycotina*), the sequence contings of the published genome are located in the nr/nt database or the Refseq genome database. Additional *Eurotiales* genomes (outside *Aspergillaceae*) are publicly accessible at the website of the Centre for Structural and Functional Genomics (Concordia University Montreal, Canada; https://gb.fungalgenomics.ca/portal/). We also included some species from the 1000 Fungal Genomes Project (http://1000.fungalgenomes.org) exclusively available at the Mycocosm database (Joint Genome Institute, US Department of Energy) (https://mycocosm.jgi.doe.gov/mycocosm/home). For the two classes of *Pezizomycotina* for which few genome sequences are public (*Xylonomycetes*, *Pezizomycetes*), we have obtained permission to use the *hxn* complement in the genome sequences of five species lodged at JGI in our current work: *Symbiotaphrina kochii* (Project ID: 404190); *Trinosporium guianense* (Project ID: 1040180); *Gyromitra esculenta* (Project ID: 1051239); *Plectania melastoma* (Project ID: 1040543); and *Sarcoscypha coccinea* (Project ID: 1042915). TBLASTN query sequences for the 11 *hxn* genes were the full-length proteins deduced from the cDNA sequences we experimentally determined for each of the *A. nidulans hxn* genes (see Table 1 for GenBank Accession numbers). Where necessary, to confirm gene orthology amongst multiple homologous sequences, the TBLASTN hits and their surrounding sequences were further inspected for the conservation of occupied intron positions between species and for synteny with other *hxn* genes in the sequence contig identified (gene clustering). We did not use the results of automated annotation (“Models” or “mRNA” at nr/nt) nor did we use deduced protein databases for the eukaryotic (Hxn) proteins. We used a selection of autoannotated proteins for the prokaryote HxnM outgroup extracted from the nr/nt database, using the *Pseudomonas putida* cyclic imide hydrolase (GenBank AAY98498: [33]) as the BLASTP query. We manually predicted the intron-exon structure of each (*hxn*) gene, guided by comparative genomics and after (*in silico*) intron removal, deduced the encoded proteins subsequently used in phylogenetic analyses (see below). Alternative yeast nuclear codes were utilised where appropriate (*Pachysolen*: CUG=Ala, *Priceomyces*: CUG=Ser). For some species in under-represented taxa, we could use the Transcriptome Shotgun Assembly (TSA) database to obtain intronless sequences coding for full-length protein.

### Construction or Maximum likelihood trees

Criteria for identification of orthologues/paralogues is detailed for each tree. Alignments were done with MAFFT G-INS-i unless otherwise indicated, with default parameters [47, 48] (https://mafft.cbrc.jp/alignment/server/). Alignments were trimmed with BMGE with default parameters unless otherwise indicated (https://ngphylogeny.fr/workflows/wkmake/42f42d079b0a46e9, [49]. Maximum likelihood trees were constructed with PhyML 3.0. Automatic model selection by SMS (http://www.atgc-montpellier.fr/phyml [50, 51]) and drawn with FigTree v 1.4.4. Values at nodes of all trees are aLRTs (Approximate Likelihood ratio test, [52]). All trees are shown in a circular cartooned form. Trees are rooted in the specified out group. Reconciliation was done by GeneRax v1.2.3, a Maximum Likelihood based method [53] with default settings in 500 replicates. Only those transfers were considered, which were present in at least 70% of the replicates. Species tree for the reconciliation was drawn after [54, 55].

## Statements

### Data accessibility

The datasets supporting this article are included in the paper and detailed in the electronic supplementary material tables. Sequences determined by us are available on GenBank accession numbers: MT707473, MT707472, MN718567, MN718568, MN718569, MN718566, MN718565, KX585439, MT707474, MT707475.

### Authors’ contributions

ZH and CS conceived the project. EB, ZH and JA contributed to various aspects of the wet laboratory work; ZH and CS wrote the manuscript. MF discovered the additional two clusters in A. nidulans, manually curated gene models of hxnV, hxnP and hxnZ orthologs and constructed the schemes of hxn clusters for hundreds of species. CV contributed to in silico promoter analysis. CS and SK did the phylogenetic analysis. All authors analysed the results and gave final approval for publication.

## Acknowledgement

JGI sequences used in the construction of the phylogenetic tree were from the US Department of Energy Joint Genome Institute (http://www.jgi.doe.gov/) in collaboration with the user community. We thank I. V. Grigoriev for permitting the use of genome sequences included in the 1000 Fungal Genomes project and we thank J. Spatafora, J. Magnuson and R. Gazis for allowing access to the genomes of some individual species prior to publication (Project IDs: 404190, 1040180, 1051239, 1040543 and 1042915). We thank Prof. Joan M. Kelly for allowing us to cite her early genetics work. We thank László G. Nagy for critical comments on the manuscript.

## Supporting information

**S1 Fig. Exon-intron organisation of the *hxnV* coding region based on cDNA sequencing and deduced HxnV protein sequence.**

(A) Exon-intron organisation of the coding region deduced through comparison of the sequenced cDNA with the genomic DNA sequence. Introns (lower case letters) are highlighted in blue (splice site consensus sequences: 5’-donor, lariat branch point sequence, and 3’-acceptor) and grey (other intronic sequences). cDNA (GenBank: MN718569) confirms the gene model. (B) Peptidic HxnV sequence deduced from the cDNA sequence.

**S1 Table. *A. nidulans* strains used in this work.** (All strains are *veA1* mutant)

**S2 Fig. Analysis of the *hxn7* mutation by Southern blot.**

(A) Location of the second *hxn* cluster on chromosome VI (cluster 2/VI, comprising the *hxnX*, *hxnW* and *hxnV genes* shown as blue, purple and yellow arrows, respectively) within a 50 kb genomic sequence (part of TPA Accession number BN001306). The figure shows the cleavage sites of DraI (D),NdeI (Nd), SalI (S), XbaI (X), NcoI (Nc), PvuII (P) and BamHI (B) restriction endonucleases that were used in Southern blot analysis (see Panel C). The gene probe used in Southern blots (labelled as a yellow box above the *hxnV* gene) was a 486 bp fragment of *hxnV* (probe), obtained by using “hxnV AS frw” (cagcgtcaagtctcatatctatactg) and “hxnV AS rev” (cagagcacgggtacaaagaaggtg) as PCR primers. Yellow arrows above the 50 kb genomic region show, for each enzyme used, the endonuclease-cleaved fragments that hybridize with the probe. (B) Predicted fragments obtained by restriction of the 50 kb genomic region described above. (C) Southern blots result with the HZS.145 control (*hxn^+^*) and HZS.697 *hxn7* mutant (*hxn7*) strains. Southern hybridization was carried out with the DIG-DNA labelling- and detection kit (Roche) on restriction endonuclease (listed in Panels A and B) digested total DNA of the *hxn^+^* and *hxn7* strains. M: DIG-labelled DNA Molecular Weight Marker (Fermentas). The DraI digest excludes that *hxn7* could be a sizeable deletion. An inversion internal to the ≈15 kb DraI fragment is excluded by several digests such as BamHI and NdeI. A translocation or more likely an insertion is compatible with the results shown.

**S2 Table. Primers used in this work.**

**S3 Fig. Correction of the gene model for the *A. nidulans hxnP* gene, locus AN11189.**

(A) The manually predicted exon-intron organisation of the coding region: introns (lower case letters) highlighted in grey; donor-, lariat- and acceptor sequences highlighted in blue. cDNA (accession KX585439) confirms this gene model. (B) Peptidic sequence of HxnP. The 4th intron splits a Met codon highlighted in purple in both panels, and this feature is conserved throughout the *Aspergillaceae* as well as in *Talaromyces*. Both the *Aspergillus* database (http://www.aspergillusgenome.org/cgi-bin/locus.pl?locus=AN11189&organism=A_nidulans_FGSC_A4) and the NCBI nr/nt database (gb|AACD01000170.1|:21152-23003 and gb|BN001306.1|:193501-195352; locus identifiers AN9176.2 and ANIA_11189, respectively) feature an erroneous gene model resulting in an incorrect HxnP protein sequence (accession numbers XP_682445 and EAA61467; TPA accession number CBF82384).

**S4 Fig. Exon-intron structure of the *hxnZ* coding region based on cDNA sequencing, and deduced HxnZ protein sequence.**

(A) Exon-intron organisation of the coding region deduced through the comparison of the sequenced cDNA with the genomic DNA sequence. Introns (lower case letters) are highlighted in grey; the donor-, lariat- and acceptor sequences are highlighted in blue. cDNA (GenBank: MT707474) confirms this gene model. (B) Peptidic sequence of HxnZ deduced from the cDNA sequence.

**S5 Fig. Distribution of hxn genes in gene clusters in selected *Eurotiomycetes* (Panel A) and in other classes of *Pezizomycotina* as well as in *Saccharomycotina* (Panel B).**

Colour coded arrows indicate specific hxn genes and relative gene orientation, as detailed: *hxnN* (dark green), *hxnM* (pink), *hxnS* (grey), *hxnT* (khaki), *hxnR* (red), *hxnP* (light green), *hxnY* (mid blue), *hxnZ* (orange), *hxnX* (ice blue), *hxnW* (purple) and *hxnV* (yellow). A single vertical line symbolises physical separation of *hxn* genes on different contigs, while a double vertical line symbolises location of genes on different chromosomes (*A. nidulans*). Grey bar: classes of *Pezizomycotina* subphylum. Purple bar: class of *Saccharomycotina* subphylum.

**S6 Fig. Phylogeny of the HxnR transcription factor.**

All putative orthologues have the same protein domain organisation as the *A. nidulans* HxnR protein [11]. HxnR orthologues are > 30 % identical to the *A. nidulans* regulatory protein outside the N-terminal DNA-binding, zinc finger domain. The “HxnR-like” proteins are Cys2His2 proteins that appear exclusively in the early divergent class of the *Pezizomycetes,* and which also are > 30 % identical to *A. nidulans* HxnR (beyond the zinc finger domain). Both *Pezizomycetes* genes have three centrally positioned introns, the first two of which positions are conserved always flank an exon with 132-138 nt length. In contrast to the orthologous *hxnR* gene, the “HxnR-like” gene is never clustered with enzyme or transporter encoding *hxn* genes and occurs in all sequenced *Pezizomycetes* species, i.e., including those that do not have *hxn* genes (except for *hxnM* paralogues). Colour code: Purple: *Pezizomycetes*, including “HxnR-like proteins” serving as an out group; Magenta: *Leotiomycetes*; Brown: *Sordariomycetes*; Orange: *Dothideomycetes*; Green: *Aspergillus;* Olive Green: *Aspergillus Section Flavi;* Cyan: *Eurotiomycetes* except *Aspergillus* and *Penicillium;* Darker Cyan: *Penicillium;* Red: *Saccharomycotina*. Species names in red: those which map outside its cognate phylogenetic clade, suggesting HGT events.

**S7 Fig. Phylogeny of the HxnT putative FMN oxidoreductase.**

Orthologous HxnT proteins are at least 40-45 % identical to the *A. nidulans* protein and can be distinguish from other homologue sequences in the genome by the synteny of *hxn* genes and the conservation of intron positions. The three-exon model of *hxnT* is broadly conserved. In some taxa, like *Monascus* [11], *hxnT* is duplicated. In such cases, we have labelled the protein from the cluster-associated *hxnT* gene, “1”. The tree is rooted with the *Saccharomycotina* clade, shown in blue, which also contains the HxnT- like protein from the early divergent yeast, *Saitoella complicata* (*Taphrinomycotina*). This species lacks the *hxnV* and *hxnX* genes, ubiquitous in all other *Ascomycota* with a minimal *hxn* complement. Colour code: Brown: *Sordariomycetes*; Purple: *Leotiomycetes*; Orange: *Dothideomycetes*. Other colours: Eurotiomycetes; Cyan: non-*Aspergillus*, non-*Penicillium;* Darker Cyan: *Penicillium;* Magenta: *Aspergillus* sections *Nidulantes/Versicolores;* Red: section *Flavi;* Green: section *Terrei. A. tanneri* (section *Circumdati*) and *A. wentii* (Section *Cremei*) are indicated with black lines. Grey: section *Nigri*. Species names marked in red indicate an anomalous phylogenetic position, suggesting HGT events. *A. bertholletius* is in blue, to distinguish it from the other members of section *Flavi*: *hxnT* is in this organism the only gene intact in an *hxn* cluster where all other are pseudo genes.

**S8 Fig. Phylogeny of the HxnV putative monooxygenase.**

Orthologous HxnV proteins are 40-45 % identical to the *A. nidulans* protein and can be distinguish from other homologue sequences in the genome by the synteny of *hxn* genes and the conservation of intron positions. Certain taxa of *Sordariomycetes* have duplicated *hxnV* genes: in the smaller clade, the genes that encode the paralogues labelled “1” are associated with other *hxn* genes. In the larger clade, the copy of the *hxnV* gene for the paralogues labelled “2” are unlinked to resident *hxn* clusters. This larger clade also includes several species, token species like *Neurospora crassa*, *Magnaporthe oryzae* and *Trichoderma reesei*, that lack the *hxn* system and harbour only a lone copy of the *hxnV* gene, suggesting a loss of the nicotinate assimilation pathway and probably a novel function for *hxnV* in these *Sordariomycetes*. The tree is rooted in the monophyletic HxnV clade of the *Saccharomycotina*. Colour code: Blue: *Saccharomycotina*; Purple: *Pezizomycetes*; Magenta: *Leotiomycetes*; Brown: *Sordariomycetes*; Black lines *Xylonomycetes*; Orange: *Dothideomycetes*; Cyan: non-*Aspergillus*, non-*Penicillium Eurotiomycetes*; Darker Cyan: *Penicillium;* Green: *Aspergillus*, except Olive green: section *Flavi;* Red: sections *Nidulantes/Versicolores*. Species names in red, those mapping outside their cognate phylogenetic clade suggesting HGT events. Species names in blue: those showing a duplication of the *hxnV* gene.

**S9 Fig. Maximum Likelihood Phylogeny of the HxnM putative imino-hydrolase protein.**

All putative HxnM orthologues are >50% identical to the *A. nidulans* HxnM protein and >50% identical to the biochemically characterised cyclic imide hydrolase from *Pseudomonas putida* (strain YZ-26) (GenBank AAY98498). Methods used in the construction of the tree are detailed in the Materials and Methods section. Numbers in nodes are aLRTs (approximate Likelihood Ratio Tests). Names of species in black: Putative HxnM proteins encoded by genes (*hxnM*) not clustered with any other *hxn* gene. Names of species in blue: *hxnM* genes clustered only with *hxnN* genes (e.g., like in *A. nidulans*, see Fig 1 or S5 Fig, or in *Lipomyces*). Names of species in red: *hxnM* included in a large cluster, encoding ≥ 4 genes. Highlighted in yellow: a monophyletic clade and putatively iso-functional clade comprising HxnMs from both clustered and un-clustered *hxnM* genes. Highlighted in light green: a clade comprising only proteins from un-clustered HxnM encoding genes, which appears as the outgroup to all clades with HxnMs from clustered genes detailed above. Colour code: Grey: prokaryotic outgroup (comprising both Bacterial and Archeal proteins); Blue: *Basidiomycota* (phylum); Red: *Saccharomycotina* (subphylum); Other colours: Pezizomycotina taxa; namely: Green: Eurotiomycetes; Olive green: *Aspergillus* section *Flavi* to highlight its anomalous position (HGT); Brown: *Sordariomycetes*; Magenta: *Dothideomycetes*; Orange: *Lecanoromycetes*; Cyan: *Leotiomycetes*; Purple: *Pezizomycetes*.

**S10 Fig. Phylogeny of the PfdA, B, and C paralogues in the *Eurotiales*.**

The *pfdA* gene is ubiquitous in *Pezizomycotina* and encodes a well conserved protein of ∼ 500 amino acids with a canonical peroxisome targeting sequence PTS-1 at its C-terminus. Generally, *Eurotiales* PfdAs are >72 % identical to the *A. flavus* protein. The *pfdB* and *pfdC* paralogues are restricted to specific taxa of the *Aspergillaceae* or *Trichocomaceae* families of the *Eurotiales* order; some species have two-while others have all three genes. The *pfdB* gene is regularly but not always, associated with the *hxn* gene cluster. The PfdB and C proteins are respectively, >60 % and >50 % identical to *A. flavus* PfdA. Typically, both paralogues are shorter than ubiquitous PfdA; multiple sequence alignments of the three paralogues (from one species) show a distinct gap in the middle of the alignment. All PfdAs and PfdBs feature a canonical PTS-1 [56] but some PdfCs appear to have lost the canonical signal sequence for peroxisome entry. All *Eurotiales* Pfd paralogue proteins included show the same domain organisation and their encoding genes have a conserved exon/intron structure with five exons (see Sup. Figure S12). In *Aspergillus*, the four introns in *pfdA* are confirmed by non-overlapping EST clones accessions DR703303 (introns 1 and 2 near the ATG) and CO136618 (introns 3 and 4 near the STOP). The exon 2 is always 59 nt long and exon 4 is always 74 nt long in nearly all Pfd paralogues. The size of the large central exon 3 distinguishes *pfdA* from its *B* and *C* paralogues. Tree construction was as detailed in Material and Methods, except that a Blosum 30 matrix was used for the BMGE alignment trimming. The species involved in the *hxn* cluster transfer (*Penicillium*, *Talaromyces*, *Aspergillus* section *Flavi*) are in red. Species names in magenta are *Aspergillu*s of sections other than *Flavi* which also include a *pfdB* gene in their *hxn* gene clusters. Colour code: Green: Outgroups *Pezizomycetes* and *Lecanoromycetes* (all PfdA; these taxa do not feature the *pfdBC* paralogues); Cyan: *Eurotiales* other than *Aspergillus* and *Penicillium*; Darker Cyan: *Penicillium;* Other colours: sections of the genus *Aspergillus.* Magenta: sections *Nidulantes*/*Versicolores*; Red: section *Flavi*; Orange: sections *Fumigati*/*Clavati*; Brown: sections *Terrei*/*Candidi*; Yellow: section *Aspergillus*; Grey: section *Nigri*; Purple: section *Circumdati*.

**S11 Fig. Intron positions in *pfdA*, *pfdB*, and *pfdC* paralogues in *Aspergillus nomius*.**

According to the data of *A. nomius* whole genome sequence project (JNOM00000000, https://www.ncbi.nlm.nih.gov/nuccore/JNOM00000000), all three *pfd* paralogues have four introns in conserved positions, two near the 5’end of the CDS and two near the 3’. Introns (lower case letters) are highlighted in blue (splice site consensus sequences: 5’- donor, lariat branch point sequence, and 3’-acceptor) and grey (other intronic sequences). The sizes of the small exons 2 and 4 are absolutely conserved throughout the orthologues. The size of the large central exon 3 distinguishes the three paralogues. In the ubiquitous *pfdA* gene exon 3 is 30 codons longer than the exon 3 of *pfdB* and 20 codons longer than exon 3 of *pfdC*. The canonical PTS-1 [56] has been lost in some but not all PfdC paralogues that supports a scenario of gene duplication of *pfdA* with simultaneous or subsequent cluster integration (mean similarity between *A* and *B* paralogues 65% compared with 88% of *A* orthologues among themselves). The third paralogue *pfdC* (mean identity between *A* and *C* paralogues 57%) present in section *Flavi,* and in a number of *Talaromyces* and *Penicilium* species and in a few species of other clades (supplementary fig. S11), which is consistent with a basal duplication followed by several episodes of loss completely unrelated to the evolution of the *hxn* cluster.

**S12 Fig. Phylogeny of HxnS in the *Pezizomycotina*.**

The tree is shown in a circular, cartoon form. Tree was constructed as indicated in Materials and Methods, except that MAFFT E-INS-i was used for the alignment. The HxnS sequences included are from Ámon et al. [11] to which orthologue proteins from some more recently published *Aspergillus*, *Penicillium* and *Talaromyces* species were added: *A*. *sclerotiales*, *A*. *tanneri*, *P*. *steckii*, *T*. *wortmannii*, and *T*. *picheae*. Colour code: Blue: Outgroups, including non-fungal xanthine hydroxylases and selected HxA orthologues; Purple: *Pezizomycetes*; Magenta: *Leotiomycetes*; Brown: *Sordariomycetes*; Black (lines): *Xylonomycetes*; Orange: *Dothideomycetes*; Cyan: non-*Aspergillus*, non-*Penicillium Eurotiales* (such as *Talaromyces*); Darker Cyan: *Penicillium* genus; Red: Sections *Nidulantes/Versicolores* (also called subgenus *Nidulantes* by Houbraken and Samson, 2011); Grey: Section *Nigri;* Green: all other *Aspergilli*. The three *Penicillium* species and *Aspergillus sclerotialis*, which cluster together within the *Sordariomycetes* are shown in red. The duplicated HxnS sequences of *A. tanneri* are shown in blue. The three *Talaromyces* species which conserve HxnS are shown in magenta.

