## Supplementary material for "Genome organisation and evolution of a eukaryotic nicotinate co-inducible pathway": S1 Fig, S2 Fig, S3 Fig, S4 Fig, S5 Fig, S6 Fig, S7 Fig, S8 Fig, S9 Fig, S10 Fig, S11 Fig, S12 Fig, S1 Table, S2 Table

**A**

>*hxnV*

ATGGCTCGCAGCGCAGATCACGACCAAGGCTTCGCAAACATCGATACCGATGCCGCACCAGCCGGCGAGACGACT
GTCGTAATTGTCGGCGCAGGACCTTCGGGGCTGATGCTTGCgtgggtttacgcccgcccgttccttcaagtgcgg
atgtgttgacaattgccagAGTGAACCTCGTCCGCTTGGGTACCCCCATCGTCCTGCTCGACGATCGGCCCGACA
AGACGTCAACCGGAAAGGCAGATGGAATCCAGCCCAAGACGATTGAGACGCTGAAGCAGCTCAGACTTGCAGATA
AACTATTGCGGGACGGGGCGCGTATCTACGATATTTCATTCTGGgtaggcccactccttctacggccgtcgttgg
caaattagacatgcagggatatgcgatatattatatgcgctgatagctgatacggtgcgagtgcagGATTCAACA
GAATCTCACCCTTTGAGGCGGAAGGGTCGCCAGACTCATTACCCGGATCACCTGGTCGGTGCATCGGACCCGTAT
ATTCTCCTCGTGCACCAAGGGATGTTGGAGGACGTGCTGATCGATGACCTTGCGGAGCGAGGCGTCACTGTTACG
CGGAATAGTTCATTTCTGTCTTGTTCGAGAAATCCATCTAAAAAATTAGATGTCGTCTACGAAGATCAGAGCACG
GGTACAAAGAAGGTGATCCAGACAGAATACCTCGTTGGTTGTGACGGCGCGCGATCGAGCGTGCGGGAGTTCATC
CCTGATGCGCAGCTGGAGGGTGAGATGACAAATGCCAGCTGGGGAGTTCTTGACGgtttgttggcattcattccc
cttaacctgccgtatgcgaccattgctgaagcagttactagGCGTCATTGAGACCGACTTTCCCGATTTATGGAG
CAAAGTTGCCGTGCGCACCCATACCGTCGGCTCCCTCTTGTGGATTCCGCGAGAACGAGGCATGACGAGGTTGTA
CGTTGAGTTGAGCGCGACAGCCGGAGAACGCATTGATAAGGCTAAGGCAACACCGCAATATGTCATGGAGCGCGC
AAAGGAGGCAATGAAGCCGTTTAGTCTGGAGTGGAAATCGATCGgtgcgcttacgacctcactttaattaggttg
tcagtatagatatgagacttgacgctgacaaaaactgtaaagAATGGTTCGGAAACTACGTCGTTGGGCAGCGCG
TGGCGAGACATTTCTCTGATCCTGATTACCAGATCTTCATTGCTGGCGATgcacgtccccttccgtctgacctct
ttctcagtttgtctaacaggttctttagGCCGGTCACTGCCACTCTGCGCTCGCCGCCCAAGGTGCAAACACCAG
TATGCACGACTCTTTCAACCTGGCATGGAAGCTGAACCTAGTGGCGCGTGGCCTAGCCTCTCCGTCTCTGCTGGA
GACTTACGAAACTGAGCGGCGCAAGATCGCAAACGACCTCATTGCCTTTGACGCCGAGCACTGTGCTGCATTTGA
GGCAGGCGAAGCCGCCCTTGCCAGGAACTTTGATGAGAATATCCGGTTCATCTCTGGGGTCGGAGCGGAGTATGA
CGCTAGCATTTTGACGCAAACCAAAGTATCAGACGCTGGGAAGGGATCCAGAAGATTGAAGCCAGGGGCGCTCTT
AATCCCAGCCAAAGCGACGCGGTACATCGATGCGAATCCGGTCGATATCCAGCTTGATGTCCCACTTTTGGGTCA
ATTCAGACTGTATTTCCTCATTCCGAATGTTAGCGCAGCGAAGGAGAAAGGATTCCTTGAGGTAGTCTGCCAGAT
TCTCAGCAGCCCGACGTCCATTCTAGCAATTTCTGCCGAGAAAGCAAAAGAATCCTACACCTCTCGCTCGCGAGG
CTGGTCTGCAACCGACGCTTACCAGGTTCCAGAACGGTACACTACTGTTTCTGAGATAATAACGCTCTCTCTCAT
TTCGGGCTCGAAGAGGGAGGTCTTCGAGATAGCGGATCTCCCGCTTGCGTTGCAGAAGAGCCGGTGGACTGTGTA
TCTGGATGATGTGGAAGGGTGTATTGAGAAGTGGGTTGGGGAGCTGCCAGAAACACAGGCTGGGATTGTGCTAGT
GAGGCCAGACGGGTATGTTGCGGGCCTTAGGGTTTGGGATCTCGGTGAGGGTGAGGTGGCAAGACGGTGGGTGGA
GGAGTATTTTGGATTCTTCTTGTGA

**B**

>HxnV

MARSADHDQGFANIDTDAAPAGETTVVIVGAGPSGLMLAVNLVRLGTPIVLLDDRPDKTSTGKADGIQPKTIETL
KQLRLADKLLRDGARIYDISFWDSTESHPLRRKGRQTHYPDHLVGASDPYILLVHQGMLEDVLIDDLAERGVTVT
RNSSFLSCSRNPSKKLDVVYEDQSTGTKKVIQTEYLVGCDGARSSVREFIPDAQLEGEMTNASWGVLDGVIETDF
PDLWSKVAVRTHTVGSLLWIPRERGMTRLYVELSATAGERIDKAKATPQYVMERAKEAMKPFSLEWKSIEWFGNY
VVGQRVARHFSDPDYQIFIAGDAGHCHSALAAQGANTSMHDSFNLAWKLNLVARGLASPSLLETYETERRKIAND
LIAFDAEHCAAFEAGEAALARNFDENIRFISGVGAEYDASILTQTKVSDAGKGSRRLKPGALLIPAKATRYIDAN
PVDIQLDVPLLGQFRLYFLIPNVSAAKEKGFLEVVCQILSSPTSILAISAEKAKESYTSRSRGWSATDAYQVPER
YTTVSEIITLSLISGSKREVFEIADLPLALQKSRWTVYLDDVEGCIEKWVGELPETQAGIVLVRPDGYVAGLRVW
DLGEGEVARRWVEEYFGFFL

**Supplementary Figure S1: Exon-intron organisation of the *hxnV* coding region based on cDNA sequencing and deduced HxnV protein sequence.** Panel A. Exon-intron organisation of the coding region deduced through comparison of the sequenced cDNA with the genomic DNA sequence. Introns (lower case letters) are highlighted in blue (splice site consensus sequences: 5’-donor, lariat branch point sequence, and 3’-acceptor) and grey (other intronic sequences). cDNA (GenBank: MN718569) confirms the gene model. Panel B. Peptidic HxnV sequence deduced from the cDNA sequence.

A

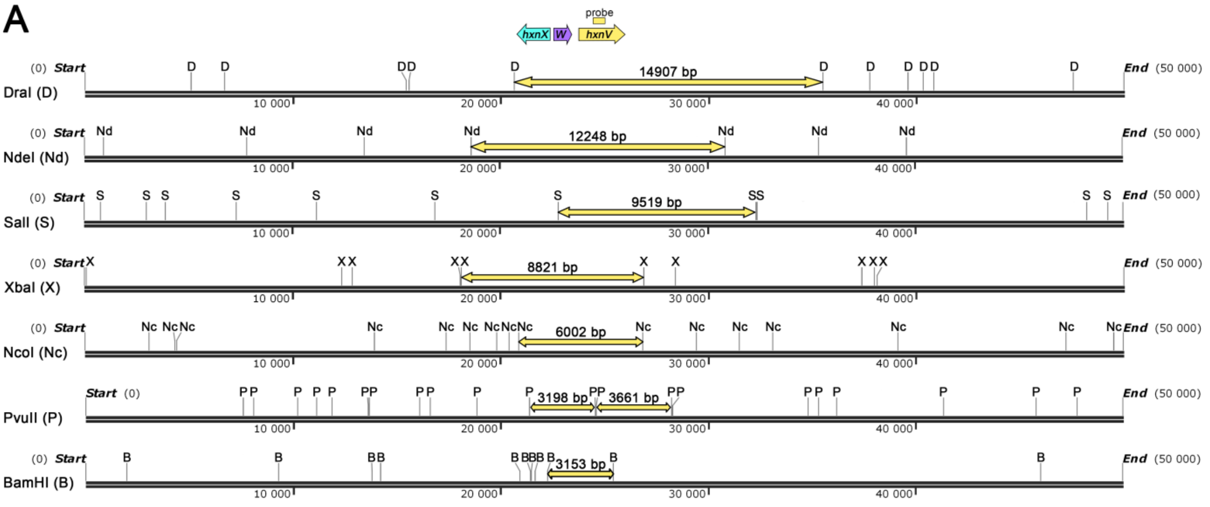


B

| Restiction enzyme | Cleavage site on chromosome VI, 50 kb genomic region comprising the second *hxn* cluster (cluster 2/VI) (bp) |
|---|---|
| DraI | 5119, 6732, 15483, 15585, 20652, 35559, 37820, 39622, 40376, 40863, 47623 |
| NdeI | 925, 7821, 13510, 18630, 30878, 35388, 39658 |
| SalI | 791, 2973, 3906, 7303, 11186, 16898, 22852, 32371, 32432, 48310, 49287 |
| XbaI | 88, 12375, 12892*, 18110, 18129, 26950*, 28476, 37453, 38008, 38183 |
| NcoI | 3094, 4312, 4441, 13950, 17389, 18556, 19858, 20434, 20895, 26897, 29444, 31540, 33164, 39200, 47302, 49567 |
| PvuII | 7581, 8109, 10203, 11156, 11859, 13649, 13674, 16094, 16617, 18886, 21395, 24593, 24619, 28280, 28294, 34842, 35380, 36228, 41390, 45856, 47822 |
| BamHI | 1969, 9318, 13830, 14248, 20967, 21437, 21491, 21665, 22297, 25450, 46094 |

C

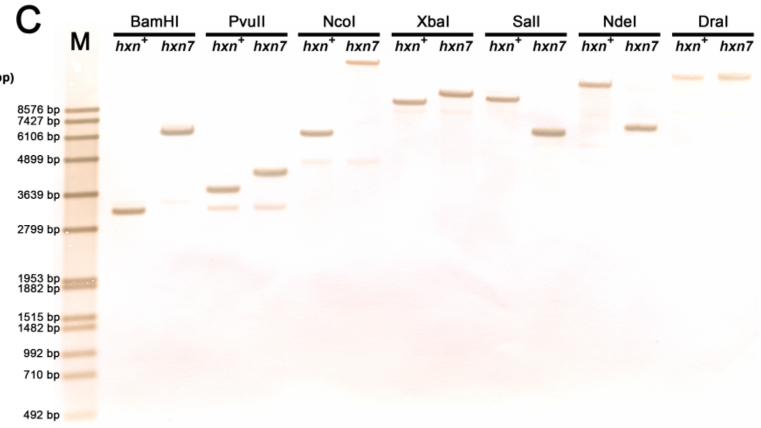


**Supplementary Figure S2: Analysis of the *hxn7* mutation by Southern blot.** Panel A. Location of the second *hxn* cluster on chromosome VI (cluster 2/VI, comprising the *hxnX*, *hxnW* and *hxnV genes* shown as blue, purple and yellow arrows, respectively) within a 50 kb genomic sequence (part of TPA Accession number BN001306). The figure shows the cleavage sites of DraI (D),NdeI (Nd), SalI (S), XbaI (X), NcoI (Nc), PvuII (P) and BamHI (B) restriction endonucleases that were used in Southern blot analysis (see Panel C). The gene probe used in Souther blots (labeled as a yellow box above the *hxnV* gene) was a 486 bp fragment of *hxnV* (probe), obtained by using “hxnV AS frw” (cagcgtcaagtctcatatctatactg) and “hxnV AS rev” (cagagcacgggtacaaagaaggtg) as PCR primers. Yellow arrows above the 50 kb genomic region show, for each enzyme used, the endonuclease-cleaved fragments that hybridize with the probe. Panel B. Predicted fragments obtained by restriction of the 50 kb genomic region described above. Panel C. Southern blots result with the HZS.145 control (*hxn*$^+$) and HZS.697 *hxn7* mutant (*hxn7*) strains. Southern hybridization was carried out with the DIG-DNA labeling- and detection kit (Roche) on restriction endonuclease (listed in Panel A and B) digested total DNA of the *hxn*$^+$ and *hxn7* strains. M: DIG-labelled DNA Molecular Weight Marker (Fermentas).

The DraI digest excludes that *hxn7* could be a sizeable deletion. An inversion internal to the ~15 kb DraI fragment is excluded by several digests such as BamHI and NdeI. A translocation or more likely an insertion is compatible with the results shown.

**A**

>*hxnP*

ATGGGCGCCACTGCCACCGATATTGAGAAGGTTCCATCGGCCGGGACACCGGACGAGCCCAAGGCCGGCGAGACT
AATGTCTATGTTGACACCGAGGCGGAGAAGAGTTTCGgtatgttgatcgagcggctcatgatcaatgagtcggaa
gggacgcgggaattgctgacccgtaccagTCCGCAAGGTGGACTTCTTTGTCCTGCCTATGCTATGTTTGgtata
ttgccctgatccctgtggtagacaggcggcgggtataggctaaccgcggcagATGTACTTCTTTGACTGCATGGA
TCGAgtatgttcagcctccctctactcgcttgaataaaaaaaacaacggcacactaatttagactatagAGCAA
CCTCGCCAACGCCAAAACGGACGGCCTGGAAGAGGACATCAATCTCAAGGGCAACGAATACTCGCTCCTCATCCT
GCTCTTCTATATCCCCTTTGGCCTGTTTGATCTGCCATGGAACCTGCTGATCAAGCGGTACTCGGCACGGATTAT
GCTCTCGCTCAgtaggtctttcatccgcagtgggaaaccctaatggctgataggacggtacgcagTGACCGTCGT
CTGGGGTATCTGCGCGTTGTGTCAATGCGCAGCAAATAACTTCGGCGGATTACTCGCTATTCGCATTATCCTAGG
gtatgtttcgcgcgcagagatcctgccaggatgtgactgacaaaaccgccggcaagAGTCTTTGAAGCAGGCTTC
TTCGCCGGCTCGACCTTCTACTTCACGCTCTTCTACACCCGCAATGAGATGGGGTTTCGGCTCGCGGTCCTGCAG
TCCTTCGCCGTGCTGGCGTCGGCATTCAGCGGCTTGATCTCGTTCGGCCTATTCCAGATAAACCATTCCGCCGTG
AAGGGATGGCAGTGGCTCTTTATTGTCGAGGGGGCCATGACGCTGATTATTGGAGTGATTGGGTTCTGGTGGTTA
CCGGACACAGCCCAGAGCGCGTGGTTTCTCACGCAGCGAGAACGAGACGCGGCTTCTGCCAGGCTCCTGAGGGAT
ACGTCTGCAGAGATCGAGACAAAACTGGAGCTGAAGGCTGCATTTCAAACATGGAGCGACTGGAAGTTTCCCATC
TGGGCTGTTATCACCTTTTCCTACCCGGTTGCGTATGCGACGGCGATGAACTTCTTTCCCATTgtatgtccccga
aagctacattaccctttttttttttttttacattattatcaggttatttttactgacgacgcagATCGTGGCTCG
CCTCGGCTACTCCGTCGTCAAGACGAACCTCTGGACTGTAGCGCCTAATCTCGTTGGCGCGGTGGTACTGCTCGT
CGTCGCTAAGTCATCTGATATCTTCCGCGAGCGGTCCTTGCATATCATCTTTAGCCTGACGGTTTCACTGGTGGG
AATGTTGATCCTGGCTAGTATTGATGTCTCGCATAACAAGGGCGTCTCGTACTTCGCCTGCTTCTTACTCGCATC
TGGTGCATACATTCCGACGTGTCTCGTGCACGCCTGGCACAATAACAATAACACGAACGAGAACTCCCGCGCTGC
TAACACTGGCTTCTTTGTCGGGCTGGGCAATATCGCTGGTGTTCTGAGTGCCGCTACCTTCAGGACAGAGTATGC
GCCCAAGTATGTGCCCACGCTGGTTGCGACGTGTGCGTGTAACGGGGTCTGCATACTCGCTACCGCCTTTATGGG
CACTTGGATGAGGCTGGAGAACCGGCGCAAAGACAAGGAGCAGGGTGCTCGAATTGTTGCCGGGCAGGTCGAGAC
GCGGATGCTGGCAGACGGCGAGAAGAGTCCAGAGTGGCGGTATTTTCTGTAG

**B**

>HxnP

MGATATDIEKVPSAGTPDEPKAGETNVYVDTEAEKSFVRKVDFFVLPMLCLMYFFDCMDRSNLANAKTDGLEEDI
NLKGNEYSLLILLFYIPFGLFDLPWNLLIKRYSARIMLSLMTVVWGICALCQCAANNFGGLLAIRIILGVFEAGF
FAGSTFYFTLFYTRNEMGFRLAVLQSFAVLASAFSGLISFGLFQINHSAVKGWQWLFIVEGAMTLIIGVIGFWWL
PDTAQSAWFLTQRERDAASARLLRDTSAEIETKLELKAAFQTWSDWKFPIWAVITFSYPVAYATAMNFFPIIVAR
LGYSVVKTNLWTVAPNLVGAVVLLVVAKSSDIFRERSLHIIFSLTVSLVGMLILASIDVSHNKGVSYFACFLLAS
GAYIPTCLVHAWHNNNNTNENSRAANTGFFVGLGNIAGVLSAATFRTEYAPKYVPTLVATCACNGVCILATAFMG
TWMRLENRRKDKEQGARIVAGQVETRMLADGEKSPEWRYFL

**Supplementary Figure S3. Correction of the gene model for the *A. nidulans hxnP* gene, locus AN11189.** Panel A. The manually predicted exon-intron organisation of the coding region: introns (lower case letters) highlighted in grey; donor-, lariat- and acceptor sequences highlighted in blue. cDNA (accession KX585439) confirms this gene model. Panel B. Peptidic sequence of HxnP. The 4th intron splits a Met codon highlighted in purple in both panels, and this feature is conserved throughout the *Aspergillacea* as well as in *Talaromyces*. Both the *Aspergillus* database (http://www.aspergillusgenome.org/cgi-bin/locus.pl?locus=AN11189&organism=A_nidulans_FGSC_A4) and the NCBI nr/nt database (gb|AACD01000170.1|:21152-23003 and gb|BN001306.1|:193501-195352; locus identifiers AN9176.2 and ANIA_11189, respectively) feature an erroneous gene model resulting in an incorrect HxnP protein sequence (accession numbers XP_682445 and EAA61467; TPA accession number CBF82384).

**A**

>*hxnZ*

ATGGATATATCATACCCTGTCATCAATGCTGGAGGTCTCAAGAATATCGCCAGCCAGATCATCATGGAAATCGAA
CTCGACAAGCGTGAGAATCGTCCGACGGACAATGTCCCGCCGGATGACATTGGGAAAATCGAGGTCGTCGACGAT
GCCGAGATGGAGCAGTTCTACGGCTCGTCTACCACTGATGCCTACCGTCTCAAGTCAGAGCTCGTTTCGCAATGC
ATGGCAGACATTGGGATGGGGCGgtgcgttattaaatcctcgcccatcagaatacgattctaaccaacccagATT
CCAGTGGAAACTGTTCACCGTCGCCGGCTTTGGCTGGATCGTCGATAATTTCTGTTCACAGGGGATATCGGCTGT
CCAGCCACCAATCCAGCAGGAGTTCAGCGGCATCAAGCAGGTGAGCTACAGCTCGGTGGCGTATTATGTGGGAAT
GATCATCGGTGCCTCATTCTGGGGCATTTCCTCCGACTTGATCGGCCGCAAGCCTGCCTTCAACTCGACGTTAGC
AATCGCGGGCATTTTCCTATGTGCCGCCGCGGGGACGTCGAACTTCATCGCCTTTAGTGCTCTCTGGGCAGTAAT
AGGAACGGCTGCAGGTGGTAATGTGGTTTGCGACTCGATGATTCTgtgagtgggttgcaaatacttgacagagga
cgcaaggcactaatcgtgcagCCTCGAATTCATCCCCGGGAGCCACCAGTACCTCCTGACCGCGCTCAGTGGATG
GTGGAATCTCGGACAGCTTGTAGTGTCACTGCTTGCATGGGTATTCCTCGCCAACTTCAGCTGCCCAACGGATGC
GACGCCAGACACCTGTTCGCGTGCCGACAATATGGGCTGGCGATATACCCTCATCACCCTCGGCGGGCTTTCCCT
TGCCTTCACCTTCGTCCGCATCTTCGTCTTCAAGATGCCTGAAACGCCCCGGTACCTGCTCTCGCAAGGAAATGA
CCAGGCTGCGGTAGACGCAGTGAACTATGTCGCCCGTCAGAACGGAAAGCCAGAGCCGCTAACGCTGTCGATGTT
GCAGGCTATCGACGTTCGGCTGGGTTTCACCCCAAACGCAGAGGAAAGATTGTCGACGAAGGATATCCTCAAAGA
GAACATGCAGGAGTTTCGAGGAGAACATTACCAAGCTCTCTTCGCGACACGAAAGCTTTCCCAGCATACGGCATT
GATCTGGGCTGTCTGGTTGATCATTGgtatccccctaactgttccctccagaccgagcacaatatcttcgaacaa
cacccgtagactaacaatatgttagGCATCGCATACCCGCTGTATTTCAACTTTCTCCCTTCTTATCTTGCCACC
AGATTCACGCAAGATTCCTCGCTCGACCTGACCTACCGCAACTACTGCATCCAGTCAGCCGTCGGGGTCGTTGGT
CCGTTATCTGCAGCGGTCCTCGTGAATACCTTCCTCGGCCGGCGCTGGATGATGGGCATTTCCTCGATCGTAACT
GGTGTCTTTCTCTTCGCGTACGTCGGCGTCAAGACTCCAATGTCGAGTCTCGCGTTTTCGTGTGTCACAGGGTTG
TTAGCGAATTTTGgtacgccagccgccactcctcacaccactggcctcctttccttcagctaatcagttgtcagA
GTACGCCATAATGTATGCCTTCACGCCGGAGTCCTTTCCTGCTCCTCATCGCGGCACAGCATCAGGGACAGCGGC
ATCGCTACTTCGATTCGGCGGGCTGGTCGCGAGCTTGATTGCGTCGGAGACGGGGTTCACGACAGCACCTATCTA
TGCTAGTGCGGCGCTTTGGGTCGGCGTCGGGGTTCTATGCTTTGGATTGCCGTTTGAGACTCATGGGCATGCTGC
TATCTAG

**B**

>HxnZ

MDISYPVINAGGLKNIASQIIMEIELDKRENRPTDNVPPDDIGKIEVVDDAEMEQFYGSSTTDAYRLKSELVSQC
MADIGMGRFQWKLFTVAGFGWIVDNFCSQGISAVQPPIQQEFSGIKQVSYSSVAYYVGMIIGASFWGISSDLIGR
KPAFNSTLAIAGIFLCAAAGTSNFIAFSALWAVIGTAAGGNVVCDSMILLEFIPGSHQYLLTALSGWWNLGQLVV
SLLAWVFLANFSCPTDATPDTCSRADNMGWRYTLITLGGLSLAFTFVRIFVFKMPETPRYLLSQGNDQAAVDAVN
YVARQNGKPEPLTLSMLQAIDVRLGFTPNAEERLSTKDILKENMQEFRGEHYQALFATRKLSQHTALIWAVWLII
GIAYPLYFNFLPSYLATRFTQDSSLDLTYRNYCIQSAVGVVGPLSAAVLVNTFLGRRWMMGISSIVTGVFLFAYV
GVKTPMSSLAFSCVTGLLANFEYAIMYAFTPESFPAPHRGTASGTAASLLRFGGLVASLIASETGFTTAPIYASA
ALWVGVGVLCFGLPFETHGHAAI

**Supplementary Figure S4. Exon-intron structure of the *hxnZ* coding region based on cDNA sequencing, and deduced HxnZ protein sequence.** Panel A. Exon-intron organisation of the coding region deduced through the comparison of the sequenced cDNA with the genomic DNA sequence. Introns (lower case letters) are highlighted in grey; the donor-, lariat- and acceptor sequences are highlighted in blue. cDNA (GenBank: MT707474) confirms this gene model. Panel B. Peptidic sequence of HxnZ deduced from the cDNA sequence.

A

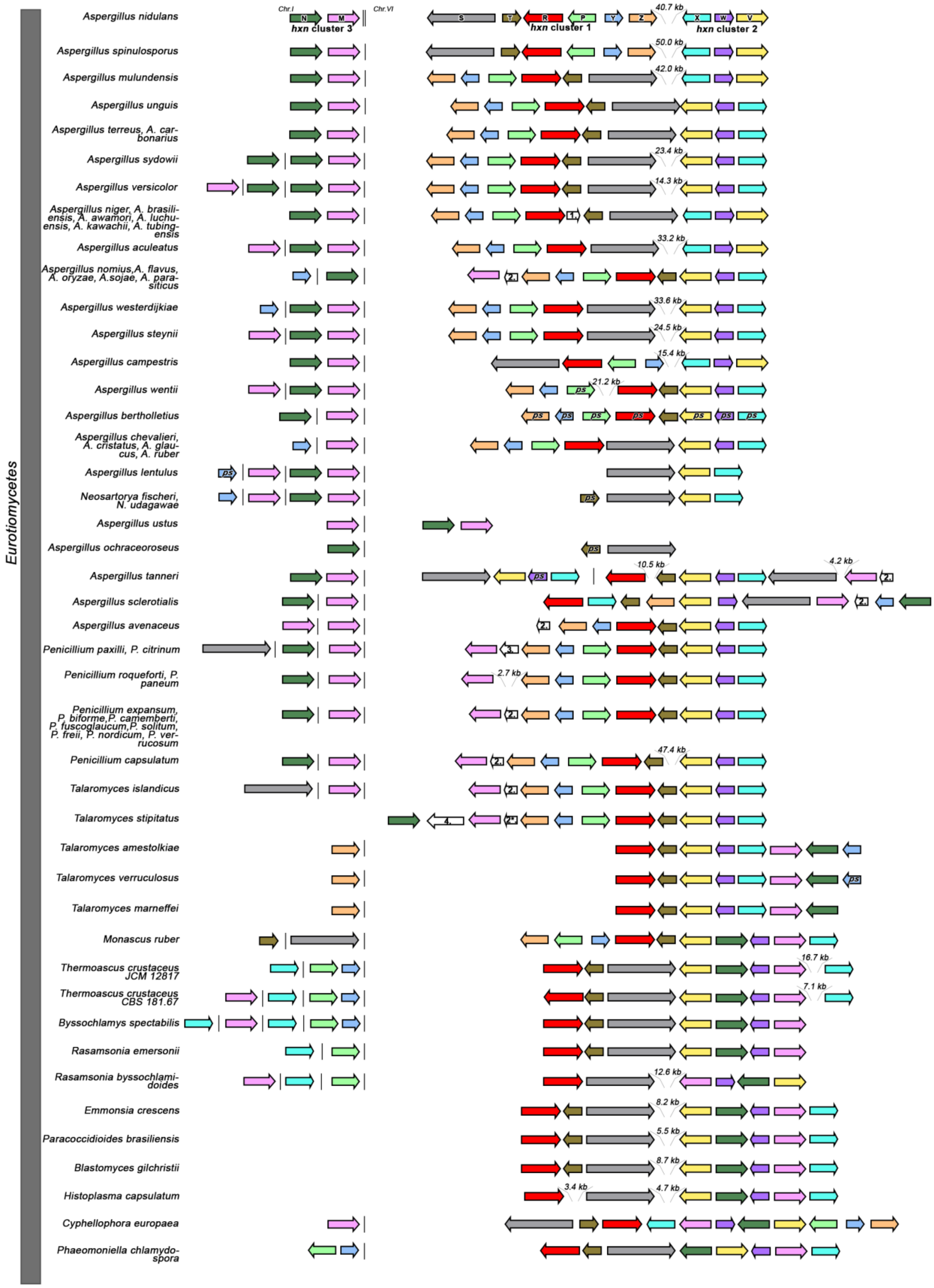


**1:** ***nfsA***
**2:** ***pfdB***
**2*:** ***pfdB* pseudogene**
**3: putative sphingosine kinase type 2 protein coding gene**
**4: putative phosphorylase protein coding gene**
***ps*: pseudogene**

B

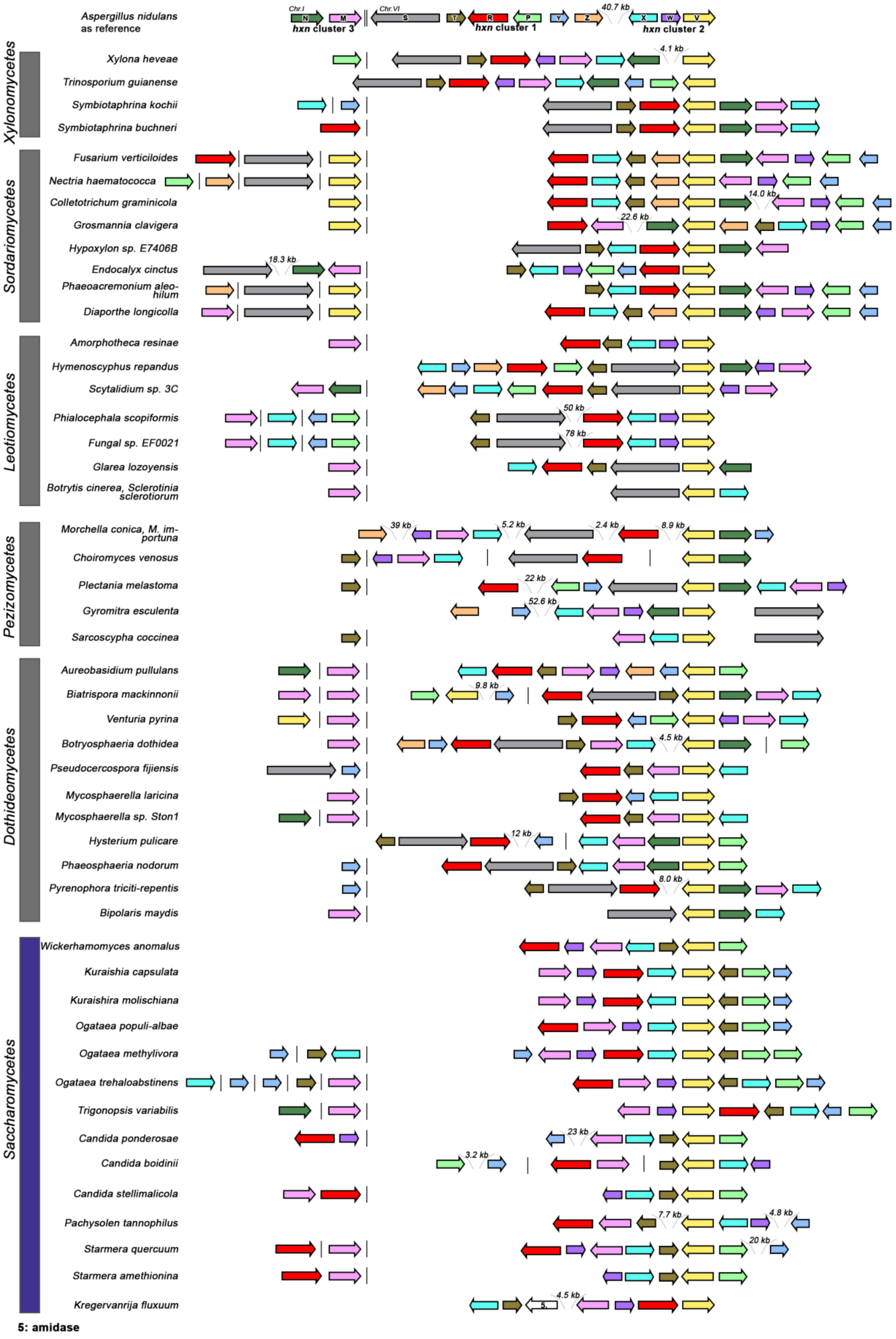
Aspergillus nidulans
as reference
Chr.I
Chr.VI
N
M
S
T
R
P
Y
Z
X
W
V
hxn cluster 3
hxn cluster 1
hxn cluster 2
40.7 kb
Xylonomycetes
Xylona heveae
4.1 kb
Trinosporium guianense
Symbiotaphrina kochii
Symbiotaphrina buchneri
Sordariomycetes
Fusarium verticiloides
Nectria haematococca
Colletotrichum graminicola
14.0 kb
Grosmannia clavigera
22.6 kb
Hypoxylon sp. E7406B
Endocalyx cinctus
18.3 kb
Phaeoacremonium aleo-
hilum
Diaporthe longicolla
Leotiomycetes
Amorphotheca resinae
Hymenoscyphus repandus
Scytalidium sp. 3C
Phialocephala scopiformis
50 kb
Fungal sp. EF0021
78 kb
Glarea lozoyensis
Botrytis cinerea, Sclerotinia
sclerotiorum
Pezizomycetes
Morchella conica, M. im-
portuna
39 kb
5.2 kb
2.4 kb
8.9 kb
Choiromyces venosus
Plectania melastoma
22 kb
Gyromitra esculenta
52.6 kb
Sarcoscypha coccinea
Dothideomycetes
Aureobasidium pullulans
Biatrispora mackinnonii
9.8 kb
Venturia pyrina
Botryosphaeria dothidea
4.5 kb
Pseudocercospora fijiensis
Mycosphaerella laricina
Mycosphaerella sp. Ston1
Hysterium pulicare
12 kb
Phaeosphaeria nodorum
Pyrenophora triciti-repentis
8.0 kb
Bipolaris maydis
Saccharomycetes
Wickerhamomyces anomalus
Kuraishia capsulata
Kuraishira molischiana
Ogataea populi-albae
Ogataea methylivora
Ogataea trehaloabstinens
Trigonopsis variabilis
Candida ponderosae
23 kb
Candida boidinii
3.2 kb
Candida stellimalicola
Pachysolen tannophilus
7.7 kb
4.8 kb
Starmera quercuum
20 kb
Starmera amethionina
Kregervanrija fluxuum
4.5 kb
5
5: amidase

**Supplementary Figure S5. Distribution of hxn genes in gene clusters in selected *Eurotiomycetes* (Panel A) and in other classes of *Pezizomycotina* as well as in *Saccharomycotina* (Panel B).** Colour coded arrows indicate specific hxn genes and relative gene orentiation, as detailed: *hxnN* (dark green), *hxnM* (pink), *hxnS* (grey), *hxnT* (khaki), *hxnR* (red), *hxnP* (light green), *hxnY* (mid blue), *hxnZ* (orange), *hxnX* (ice blue), *hxnW* (purple) and *hxnV* (yellow). A single vertical line symbolises physical separation of *hxn* genes on different contigs, while a double vertical line symbolises location of genes on different chromosomes (*A. nidulans*). Grey bar: classes of *Pezizomycotina* subphylum. Purple bar: class of *Saccharomycotina* subphylum.

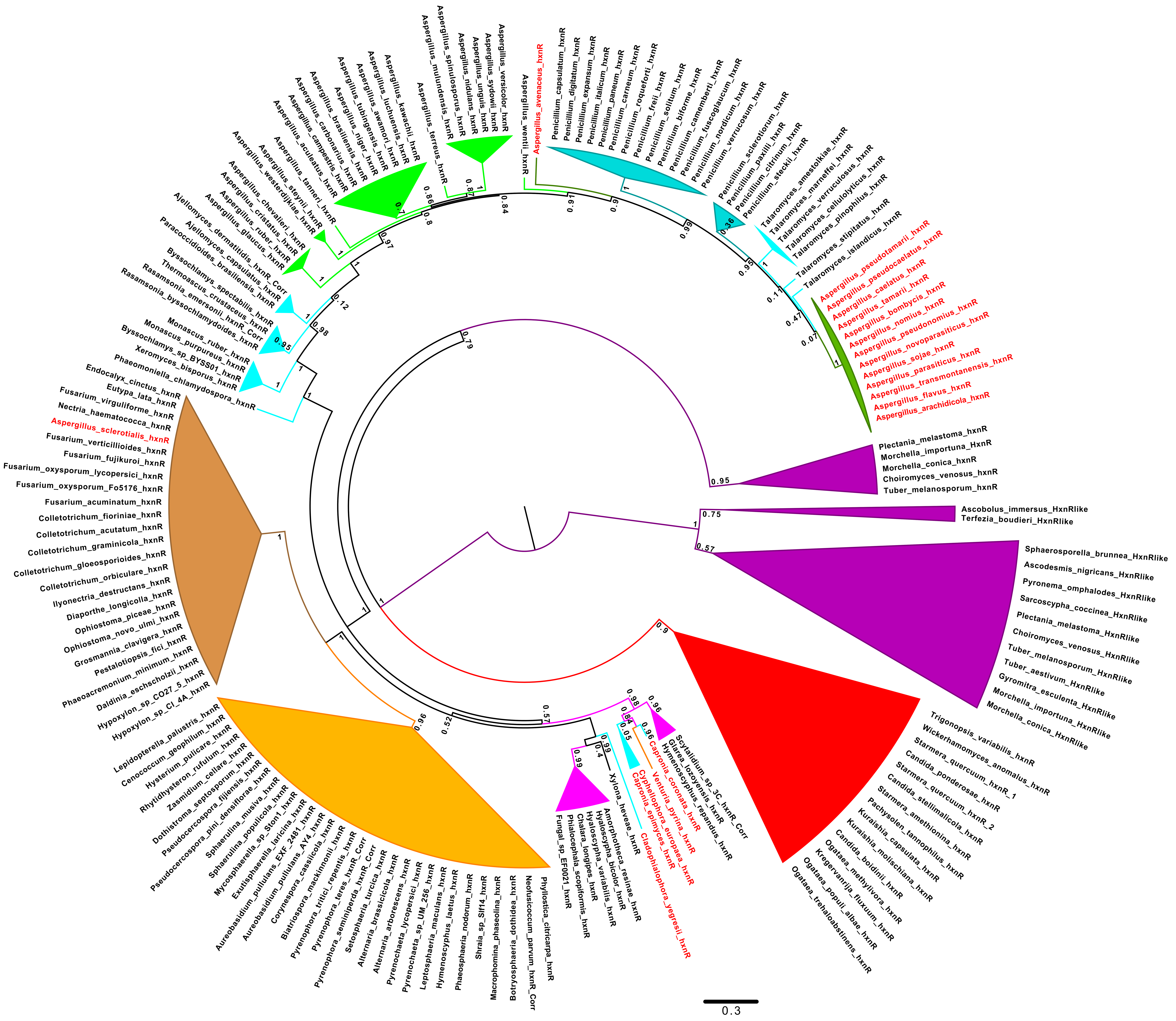
Aspergillus_versicolor_hxnR
Aspergillus_sydowii_hxnR
Aspergillus_unguis_hxnR
Aspergillus_nidulans_hxnR
Aspergillus_mulundensis_hxnR
Aspergillus_spinulosporus_hxnR
Aspergillus_terreus_hxnR
Aspergillus_kawachii_hxnR
Aspergillus_luchuensis_hxnR
Aspergillus_awamori_hxnR
Aspergillus_tubingensis_hxnR
Aspergillus_niger_hxnR
Aspergillus_brasiliensis_hxnR
Aspergillus_carbonarius_hxnR
Aspergillus_campestris_hxnR
Aspergillus_aculeatus_hxnR
Aspergillus_tanneri_hxnR
Aspergillus_steynii_hxnR
Aspergillus_westerdijkiae_hxnR
Aspergillus_chevalieri_hxnR
Aspergillus_cristatus_hxnR
Aspergillus_ruber_hxnR
Aspergillus_glaucus_hxnR
Ajellomyces_dermatitidis_hxnR_Corr
Ajellomyces_capsulatus_hxnR
Paracoccidioides_brasiliensis_hxnR
Byssochlamys_spectabilis_hxnR
Thermoascus_crustaceus_hxnR
Rasamsonia_emersonii_hxnR_Corr
Rasamsonia_byssochlamydoides_hxnR
Monascus_ruber_hxnR
Monascus_purpureus_hxnR
Byssochlamys_sp_BYSS01_hxnR
Xeromyces_bisporus_hxnR
Phaeomoniella_chlamydospora_hxnR
Endocalyx_cinctus_hxnR
Eutypa_lata_hxnR
Fusarium_virguliforme_hxnR
Nectria_haematococca_hxnR
Aspergillus_sclerotialis_hxnR
Fusarium_verticillioides_hxnR
Fusarium_fujikuroi_hxnR
Fusarium_oxysporum_lycopersici_hxnR
Fusarium_oxysporum_Fo5176_hxnR
Fusarium_acuminatum_hxnR
Colletotrichum_fioriniae_hxnR
Colletotrichum_acutatum_hxnR
Colletotrichum_graminicola_hxnR
Colletotrichum_gloeosporioides_hxnR
Colletotrichum_orbiculare_hxnR
Ilyonectria_destructans_hxnR
Diaporthe_longicolla_hxnR
Ophiostoma_piceae_hxnR
Ophiostoma_novo_ulmi_hxnR
Grosmannia_clavigera_hxnR
Pestalotiopsis_fici_hxnR
Phaeoacremonium_minimum_hxnR
Daldinia_eschscholzii_hxnR
Hypoxylon_sp_CO27_5_hxnR
Hypoxylon_sp_CI_4A_hxnR
Lepidopterella_palustris_hxnR
Cenococcum_geophilum_hxnR
Hysterium_pulicare_hxnR
Rhytidhysteron_rufulum_hxnR
Zasmidium_cellare_hxnR
Dothistroma_septosporum_hxnR
Pseudocercospora_fijiensis_hxnR
Pseudocercospora_pini_densiflorae_hxnR
Sphaerulina_musiva_hxnR
Sphaerulina_populicola_hxnR
Mycosphaerella_sp_Ston1_hxnR
Exutisphaerella_laricina_hxnR
Aureobasidium_pullulans_EXF_2481_hxnR
Aureobasidium_pullulans_AY4_hxnR
Corynespora_cassiicola_hxnR
Biatriospora_mackinnonii_hxnR
Pyrenophora_tritici_repentis_hxnR_Corr
Pyrenophora_teres_hxnR_Corr
Pyrenophora_seminiperda_hxnR
Setosphaeria_turcica_hxnR
Alternaria_brassicicola_hxnR
Alternaria_arborescens_hxnR
Pyrenochaeta_lycopersici_hxnR
Pyrenochaeta_sp_UM_256_hxnR
Leptosphaeria_maculans_hxnR
Hymenoscyphus_laetus_hxnR
Phaeosphaeria_nodorum_hxnR
Shraia_sp_Slf14_hxnR
Macrophomina_phaseolina_hxnR
Botryosphaeria_dothidea_hxnR
Neofusicoccum_parvum_hxnR_Corr
Phyllosticta_citricarpa_hxnR
Fungal_sp_EF0021_hxnR
Phialocephala_scopiformis_hxnR
Chalara_longipes_hxnR
Hyaloscypha_variabilis_hxnR
Hyaloscypha_bicolor_hxnR
Amorphotheca_resinae_hxnR
Xylona_heveae_hxnR
Cladophialophora_yegresii_hxnR
Capronia_epimyces_hxnR
Cyphellophora_europaea_hxnR
Venturia_pyrina_hxnR
Capronia_coronata_hxnR
Hymenoscyphus_repandus_hxnR
Glarea_lozoyensis_hxnR
Scytalidium_sp_3C_hxnR_Corr
Ogataea_trehaloabstinens_hxnR
Ogataea_populi_albae_hxnR
Kregervanrija_fluxuum_hxnR
Ogataea_methylivora_hxnR
Candida_boidinii_hxnR
Kuraishia_molischiana_hxnR
Kuraishia_capsulata_hxnR
Pachysolen_tannophilus_hxnR
Starmera_amethionina_hxnR
Candida_stellimalicola_hxnR
Starmera_quercuum_hxnR_2
Candida_ponderosae_hxnR
Starmera_quercuum_hxnR_1
Wickerhamomyces_anomalus_hxnR
Trigonopsis_variabilis_hxnR
Morchella_conica_HxnRlike
Morchella_importuna_HxnRlike
Gyromitra_esculenta_HxnRlike
Tuber_aestivum_HxnRlike
Tuber_melanosporum_HxnRlike
Choiromyces_venosus_HxnRlike
Plectania_melastoma_HxnRlike
Sarcoscypha_coccinea_HxnRlike
Pyronema_omphalodes_HxnRlike
Ascodesmis_nigricans_HxnRlike
Sphaerosporella_brunnea_HxnRlike
Ascobolus_immersus_HxnRlike
Terfezia_boudieri_HxnRlike
Plectania_melastoma_hxnR
Morchella_importuna_HxnR
Morchella_conica_hxnR
Choiromyces_venosus_hxnR
Tuber_melanosporum_hxnR
Aspergillus_arachidicola_hxnR
Aspergillus_flavus_hxnR
Aspergillus_transmontanensis_hxnR
Aspergillus_parasiticus_hxnR
Aspergillus_sojae_hxnR
Aspergillus_novoparasiticus_hxnR
Aspergillus_pseudonomius_hxnR
Aspergillus_nomius_hxnR
Aspergillus_bombycis_hxnR
Aspergillus_tamarii_hxnR
Aspergillus_caelatus_hxnR
Aspergillus_pseudocaelatus_hxnR
Aspergillus_pseudotamarii_hxnR
Talaromyces_islandicus_hxnR
Talaromyces_stipitatus_hxnR
Talaromyces_pinophilus_hxnR
Talaromyces_cellulolyticus_hxnR
Talaromyces_verruculosus_hxnR
Talaromyces_marneffei_hxnR
Talaromyces_amestolkiae_hxnR
Penicillium_steckii_hxnR
Penicillium_citrinum_hxnR
Penicillium_paxilli_hxnR
Penicillium_sclerotiorum_hxnR
Penicillium_verrucosum_hxnR
Penicillium_nordicum_hxnR
Penicillium_fuscoglaucum_hxnR
Penicillium_camemberti_hxnR
Penicillium_biforme_hxnR
Penicillium_solitum_hxnR
Penicillium_freii_hxnR
Penicillium_roqueforti_hxnR
Penicillium_carneum_hxnR
Penicillium_paneum_hxnR
Penicillium_italicum_hxnR
Penicillium_expansum_hxnR
Penicillium_digitatum_hxnR
Penicillium_capsulatum_hxnR
Aspergillus_avenaceus_hxnR
Aspergillus_wentii_hxnR
0.3

**Supplementary Figure S6. Phylogeny of the HxnR transcription factor.** All putative orthologues have the same protein domain organisation as the *A. nidulans* HxnR protein (Amon et al., 2017). HxnR orthologues are > 30 % identical to the *A. nidulans* regulatory protein outside the N-terminal DNA-binding, zinc finger domain. The “HxnR-like” proteins are Cys2His2 proteins that appear exclusively in the early divergent class of the *Pezizomycetes*, and which also are > 30 % identical to *A. nidulans* HxnR (beyond the zinc finger domain). Both *Pezizomycetes* genes have three centrally positioned introns, the first two of which positions are conserved always flank an exon with 132-138 nt length. In contrast to the orthologous *hxnR* gene, the “HxnR-like” gene is never clustered with enzyme or transporter encoding *hxn* genes and occurs in all sequenced *Pezizomycetes* species, i.e., including those that do not have *hxn* genes (except for *hxnM* paralogues). Colour code: Purple: *Pezizomycetes*, including "HxnR-like proteins" serving as an out group; Magenta: *Leotiomycetes*; Brown: *Sordariomycetes*; Orange: *Dothideomycetes*; Green: *Aspergillus;* Olive Green: *Aspergillus Section Flavi;* Cyan: *Eurotiomycetes* except *Aspergillus* and *Penicillium;* Darker Cyan: *Penicillium;* Red: *Saccharomycotina*. Species names in red: those which map outside its cognate phylogenetic clade, suggesting HGT events.

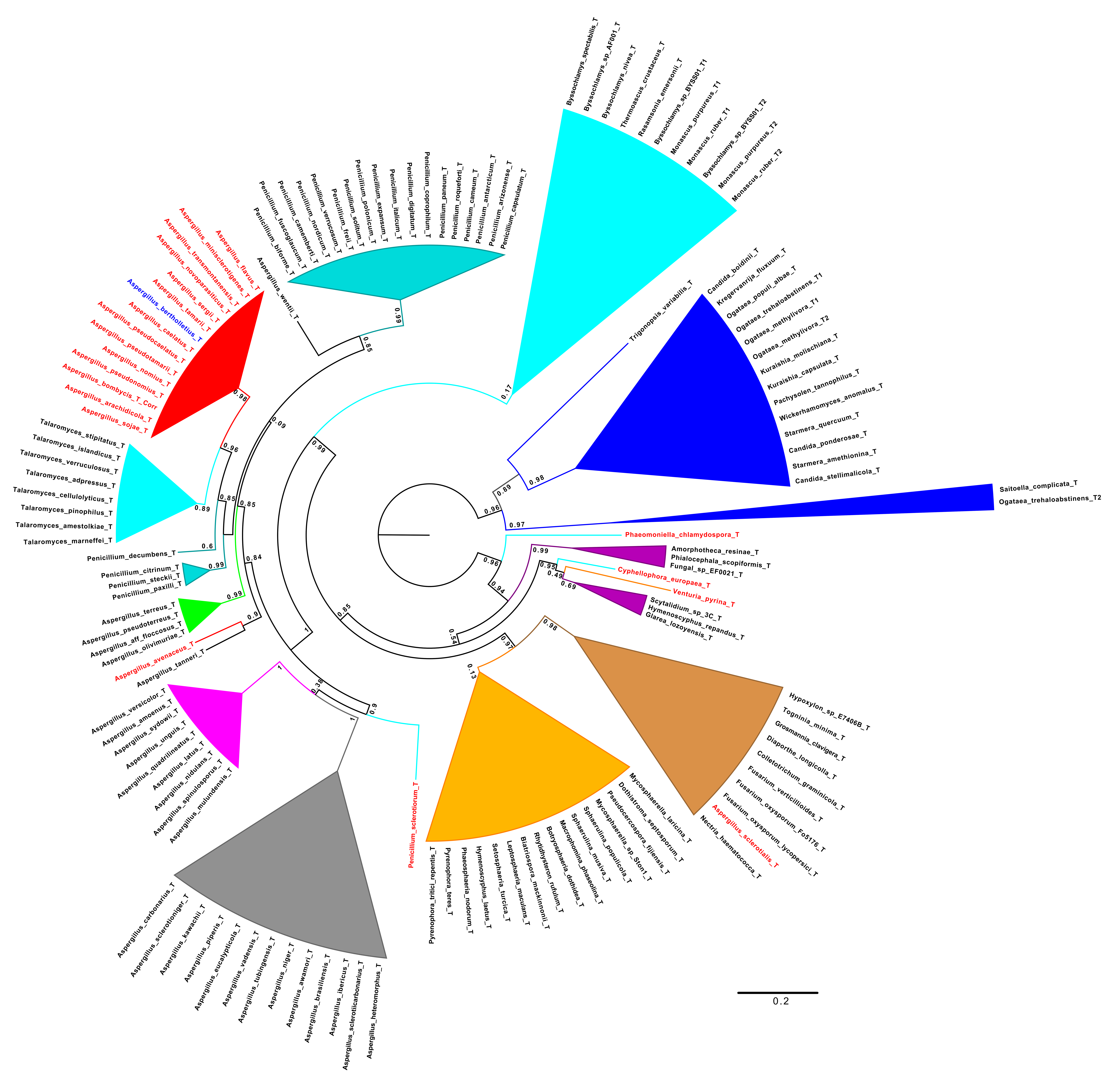

Byssochlamys_spectabilis_T
Byssochlamys_sp_AF001_T
Byssochlamys_nivea_T
Thermoascus_crustaceus_T
Rasamsonia_emersonii_T
Byssochlamys_sp_BYSS01_T1
Monascus_purpureus_T1
Monascus_ruber_T1
Byssochlamys_sp_BYSS01_T2
Monascus_purpureus_T2
Monascus_ruber_T2
Penicillium_biforme_T
Penicillium_fuscoglaucum_T
Penicillium_camemberti_T
Penicillium_nordicum_T
Penicillium_verrucosum_T
Penicillium_freii_T
Penicillium_solitum_T
Penicillium_polonicum_T
Penicillium_expansum_T
Penicillium_italicum_T
Penicillium_digitatum_T
Penicillium_coprophilum_T
Penicillium_paneum_T
Penicillium_roqueforti_T
Penicillium_carneum_T
Penicillium_antarcticum_T
Penicillium_arizonense_T
Penicillium_capsulatum_T
Aspergillus_wentii_T
Aspergillus_flavus_T
Aspergillus_minisclerotigenes_T
Aspergillus_transmontanensis_T
Aspergillus_novoparasiticus_T
Aspergillus_sergii_T
Aspergillus_tamarii_T
Aspergillus_bertholletius_T
Aspergillus_caelatus_T
Aspergillus_pseudocaelatus_T
Aspergillus_pseudotamarii_T
Aspergillus_nomius_T
Aspergillus_pseudonomius_T
Aspergillus_bombycis_T_Corr
Aspergillus_arachidicola_T
Aspergillus_sojae_T
Talaromyces_stipitatus_T
Talaromyces_islandicus_T
Talaromyces_verruculosus_T
Talaromyces_adpressus_T
Talaromyces_cellulolyticus_T
Talaromyces_pinophilus_T
Talaromyces_amestolkiae_T
Talaromyces_marneffei_T
Penicillium_decumbens_T
Penicillium_citrinum_T
Penicillium_steckii_T
Penicillium_paxilli_T
Aspergillus_terreus_T
Aspergillus_pseudoterreus_T
Aspergillus_aff_floccosus_T
Aspergillus_olivimuriae_T
Aspergillus_avenaceus_T
Aspergillus_tanneri_T
Aspergillus_versicolor_T
Aspergillus_amoenus_T
Aspergillus_sydowii_T
Aspergillus_unguis_T
Aspergillus_quadrilineatus_T
Aspergillus_latus_T
Aspergillus_nidulans_T
Aspergillus_spinulosporus_T
Aspergillus_mulundensis_T
Aspergillus_carbonarius_T
Aspergillus_sclerotioniger_T
Aspergillus_kawachii_T
Aspergillus_piperis_T
Aspergillus_eucalypticola_T
Aspergillus_vadensis_T
Aspergillus_tubingensis_T
Aspergillus_niger_T
Aspergillus_awamori_T
Aspergillus_brasiliensis_T
Aspergillus_ibericus_T
Aspergillus_sclerotiicarbonarius_T
Aspergillus_heteromorphus_T
Penicillium_sclerotiorum_T
Pyrenophora_tritici_repentis_T
Pyrenophora_teres_T
Phaeosphaeria_nodorum_T
Hymenoscyphus_laetus_T
Setosphaeria_turcica_T
Leptosphaeria_maculans_T
Biatriospora_mackinnonii_T
Rhytidhysteron_rufulum_T
Botryosphaeria_dothidea_T
Macrophomina_phaseolina_T
Sphaerulina_musiva_T
Sphaerulina_populicola_T
Mycosphaerella_sp_Ston1_T
Pseudocercospora_fijiensis_T
Dothistroma_septosporum_T
Mycosphaerella_laricina_T
Hypoxylon_sp_E7406B_T
Togninia_minima_T
Grosmannia_clavigera_T
Diaporthe_longicolla_T
Colletotrichum_graminicola_T
Fusarium_verticillioides_T
Fusarium_oxysporum_Fo5176_T
Fusarium_oxysporum_lycopersici_T
Aspergillus_sclerotialis_T
Nectria_haematococca_T
Trigonopsis_variabilis_T
Candida_boidinii_T
Kregervanrija_fluxuum_T
Ogataea_populi_albae_T
Ogataea_trehaloabstinens_T1
Ogataea_methylivora_T1
Ogataea_methylivora_T2
Kuraishia_molischiana_T
Kuraishia_capsulata_T
Pachysolen_tannophilus_T
Wickerhamomyces_anomalus_T
Starmera_quercuum_T
Candida_ponderosae_T
Starmera_amethionina_T
Candida_stellimalicola_T
Saitoella_complicata_T
Ogataea_trehaloabstinens_T2
Phaeomoniella_chlamydospora_T
Amorphotheca_resinae_T
Phialocephala_scopiformis_T
Fungal_sp_EF0021_T
Cyphellophora_europaea_T
Venturia_pyrina_T
Scytalidium_sp_3C_T
Hymenoscyphus_repandus_T
Glarea_lozoyensis_T
0.99
0.85
0.09
0.98
0.96
0.85
0.85
0.89
0.6
0.99
0.84
0.99
0.9
1
0.38
1
0.9
1
0.99
0.17
0.98
0.89
0.96
0.97
0.99
0.96
0.95
0.49
0.69
0.94
0.85
0.54
0.97
0.98
0.13
0.2

**Supplementary Figure S7. Phylogeny of the HxnT putative FMN oxidoreductase.** Orthologous HxnT proteins are at least 40-45 % identical to the *A. nidulans* protein and can be distinguish from other homologue sequences in the genome by the synteny of *hxn* genes and the conservation of intron positions. The three-exon model of *hxnT* is broadly conserved. In some taxa, like *Monascus* (Amon et al., 2017), *hxnT* is duplicated. In such cases, we have labeled the protein from the cluster-associated *hxnT* gene, "1". The tree is rooted with the *Saccharomycotina* clade, shown in blue, which also contains the HxnT-like protein from the early divergent yeast, *Saitoella complicata* (*Taphrinomycotina*). This species lacks the *hxnV* and *hxnX* genes, ubiquitous in all other *Ascomycota* with a minimal *hxn* complement. Colour code: Brown: *Sordariomycetes*; Purple: *Leotiomycetes*; Orange: *Dothideomycetes*. Other colours: Eurotiomycetes; Cyan: non-*Aspergillus*, non-*Penicillium;* Darker Cyan: *Penicillium;* Magenta: *Aspergillus* sections *Nidulantes/Versicolores;* Red: section *Flavi;* Green: section *Terrei*. *A. tanneri* (section *Circumdati*) and *A. wentii* (Section *Cremei*) are indicated with black lines. Grey: section *Nigri*. Species names marked in red indicate an anomalous phylogenetic position, suggesting HGT events. *A. bertholletius* is in blue, to distinguish it from the other members of section *Flavi*: *hxnT* is in this organism the only gene intact in an *hxn* cluster where all other are pseudo genes.

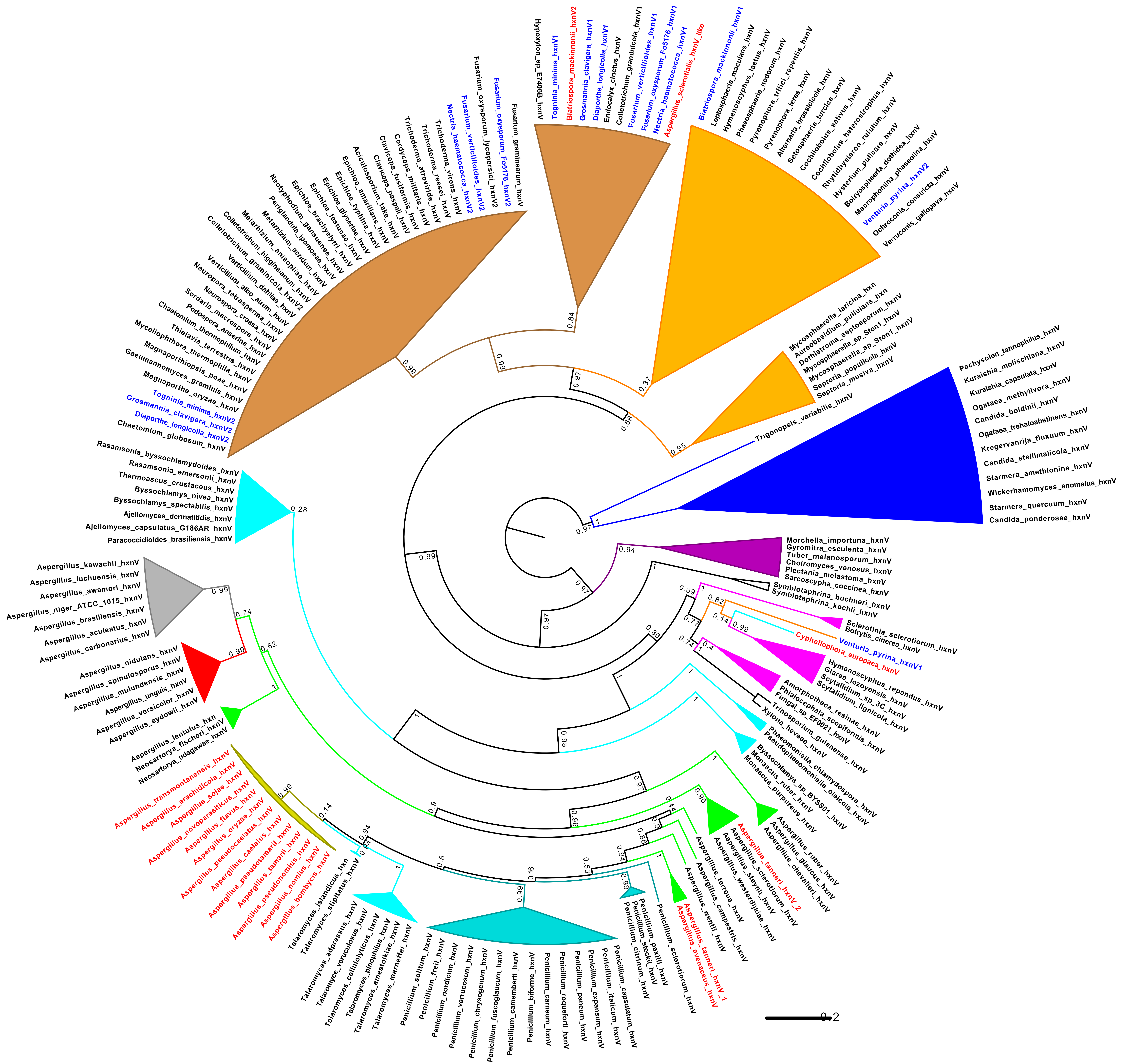
Fusarium_graminearum_hxnV
Fusarium_oxysporum_Fo5176_hxnV2
Fusarium_oxysporum_lycopersici_hxnV2
Fusarium_verticillioides_hxnV2
Nectria_haematococca_hxnV2
Trichoderma_virens_hxnV
Trichoderma_reesei_hxnV
Trichoderma_atroviride_hxnV
Cordyceps_militaris_hxnV
Claviceps_fusiformis_hxnV
Claviceps_paspali_hxnV
Aciculosporium_take_hxnV
Epichloe_amarillans_hxnV
Epichloe_typhina_hxnV
Epichloe_glyceriae_hxnV
Epichloe_festucae_hxnV
Epichloe_brachyelytri_hxnV
Neotyphodium_gansuense_hxnV
Periglandula_ipomoeae_hxnV
Metarhizium_acridum_hxnV
Metarhizium_anisopliae_hxnV
Colletotrichum_higginsianum_hxnV
Colletotrichum_graminicola_hxnV2
Verticillium_dahliae_hxnV
Verticillium_albo_atrum_hxnV
Neurospora_tetrasperma_hxnV
Neurospora_crassa_hxnV
Sordaria_macrospora_hxnV
Podospora_anserina_hxnV
Chaetomium_thermophilum_hxnV
Thielavia_terrestris_hxnV
Myceliophthora_thermophila_hxnV
Magnaporthiopsis_poae_hxnV
Gaeumannomyces_graminis_hxnV
Magnaporthe_oryzae_hxnV
Tognia_minima_hxnV2
Grosmannia_clavigera_hxnV2
Diaporthe_longicolla_hxnV2
Chaetomium_globosum_hxnV
Hypoxylon_sp_E7406B_hxnV
Tognia_minima_hxnV1
Biatriospora_mackinnonii_hxnV2
Grosmannia_clavigera_hxnV1
Diaporthe_longicolla_hxnV1
Endocalyx_cinctus_hxnV
Colletotrichum_graminicola_hxnV1
Fusarium_verticillioides_hxnV1
Fusarium_oxysporum_Fo5176_hxnV1
Nectria_haematococca_hxnV1
Aspergillus_sclerotialis_hxnV_like
Biatriospora_mackinnonii_hxnV1
Leptosphaeria_maculans_hxnV
Hymenoscyphus_laetus_hxnV
Phaeosphaeria_nodorum_hxnV
Pyrenophora_tritici_repentis_hxnV
Pyrenophora_teres_hxnV
Alternaria_brassicicola_hxnV
Setosphaeria_turcica_hxnV
Cochliobolus_sativus_hxnV
Cochliobolus_heterostrophus_hxnV
Rhytidhysteron_rufulum_hxnV
Hysterium_pulicare_hxnV
Botryosphaeria_dothidea_hxnV
Macrophomina_phaseolina_hxnV
Venturia_pyrina_hxnV2
Ochroconis_constricta_hxnV
Verruconis_gallopava_hxnV
Mycosphaerella_laricina_hxn
Aureobasidium_pullulans_hxn
Dothistroma_septosporum_hxnV
Mycosphaerella_sp_Ston1_hxnV
Mycosphaerella_sp_Ston1_hxnV
Septoria_populicola_hxnV
Septoria_musiva_hxnV
Trigonopsis_variabilis_hxnV
Pachysolen_tannophilus_hxnV
Kuraishia_molischiana_hxnV
Kuraishia_capsulata_hxnV
Ogataea_methylivora_hxnV
Candida_boidinii_hxnV
Ogataea_trehaloabstinens_hxnV
Kregervanrija_fluxuum_hxnV
Candida_stellimalicola_hxnV
Starmera_amethionina_hxnV
Wickerhamomyces_anomalus_hxnV
Starmera_quercuum_hxnV
Candida_ponderosae_hxnV
Morchella_importuna_hxnV
Gyromitra_esculenta_hxnV
Tuber_melanosporum_hxnV
Choiromyces_venosus_hxnV
Plectania_melastoma_hxnV
Sarcoscypha_coccinea_hxnV
Symbiotaphrina_buchneri_hxnV
Symbiotaphrina_kochii_hxnV
Sclerotinia_sclerotiorum_hxnV
Botrytis_cinerea_hxnV
Cyphellophora_europaea_hxnV
Venturia_pyrina_hxnV1
Hymenoscyphus_repandus_hxnV
Glarea_lozoyensis_hxnV
Scytalidium_sp_3C_hxnV
Scytalidium_lignicola_hxnV
Amorphotheca_resinae_hxnV
Phialocephala_scopiformis_hxnV
Fungal_sp_EF0021_hxnV
Trinosporium_guianense_hxnV
Xylona_heveae_hxnV
Phaeomoniella_chlamydospora_hxnV
Pseudophaeomoniella_oleicola_hxnV
Byssochlamys_sp_BYSS01_hxnV
Monascus_ruber_hxnV
Monascus_purpureus_hxnV
Aspergillus_ruber_hxnV
Aspergillus_glaucus_hxnV
Aspergillus_chevalieri_hxnV
Aspergillus_tamari_hxnV_2
Aspergillus_sclerotiorum_hxnV
Aspergillus_steynii_hxnV
Aspergillus_westerdijkiae_hxnV
Aspergillus_terreus_hxnV
Aspergillus_campestris_hxnV
Aspergillus_wentii_hxnV
Aspergillus_tamari_hxnV_1
Aspergillus_avenaceus_hxnV
Penicillium_sclerotiorum_hxnV
Penicillium_paxilli_hxnV
Penicillium_steckii_hxnV
Penicillium_citrinum_hxnV
Penicillium_capsulatum_hxnV
Penicillium_italicum_hxnV
Penicillium_expansum_hxnV
Penicillium_paneum_hxnV
Penicillium_roqueforti_hxnV
Penicillium_carneum_hxnV
Penicillium_biforme_hxnV
Penicillium_camemberti_hxnV
Penicillium_fuscoglaucum_hxnV
Penicillium_chrysogenum_hxnV
Penicillium_verrucosum_hxnV
Penicillium_nordicum_hxnV
Penicillium_freii_hxnV
Penicillium_solitum_hxnV
Talaromyces_marneffei_hxnV
Talaromyces_amestolkiae_hxnV
Talaromyces_pinophilus_hxnV
Talaromyces_cellulolyticus_hxnV
Talaromyce_veruculosus_hxnV
Talaromyces_adpressus_hxnV
Talaromyces_stipitatus_hxnV
Talaromyces_islandicus_hxn
Aspergillus_bombycis_hxnV
Aspergillus_nomius_hxnV
Aspergillus_pseudonomius_hxnV
Aspergillus_tamarii_hxnV
Aspergillus_pseudotamarii_hxnV
Aspergillus_caelatus_hxnV
Aspergillus_pseudocaelatus_hxnV
Aspergillus_oryzae_hxnV
Aspergillus_flavus_hxnV
Aspergillus_novoparasiticus_hxnV
Aspergillus_sojae_hxnV
Aspergillus_arachidicola_hxnV
Aspergillus_transmontanensis_hxnV
Neosartorya_udagawae_hxnV
Neosartorya_fischeri_hxnV
Aspergillus_lentulus_hxn
Aspergillus_sydowii_hxnV
Aspergillus_versicolor_hxnV
Aspergillus_unguis_hxnV
Aspergillus_mulundensis_hxnV
Aspergillus_spinulosporus_hxnV
Aspergillus_nidulans_hxnV
Aspergillus_carbonarius_hxnV
Aspergillus_aculeatus_hxnV
Aspergillus_brasiliensis_hxnV
Aspergillus_niger_ATCC_1015_hxnV
Aspergillus_awamori_hxnV
Aspergillus_luchuensis_hxnV
Aspergillus_kawachii_hxnV
Rasamsonia_byssochlamydoides_hxnV
Rasamsonia_emersonii_hxnV
Thermoascus_crustaceus_hxnV
Byssochlamys_nivea_hxnV
Byssochlamys_spectabilis_hxnV
Ajellomyces_dermatitidis_hxnV
Ajellomyces_capsulatus_G186AR_hxnV
Paracoccidioides_brasiliensis_hxnV
0.2

**Supplementary Figure S8. Phylogeny of the HxnV putative monooxygenase.** Orthologous HxnV proteins are 40-45 % identical to the *A. nidulans* protein and can be distinguish from other homologue sequences in the genome by the synteny of *hxn* genes and the conservation of intron positions. Certain taxa of *Sordariomycetes* have duplicated *hxnV* genes: in the smaller clade, the genes that encode the paralogues labeled “1” are associated with other *hxn* genes. In the larger clade, the copy of the *hxnV* gene for the paralogues labeled “2” are unlinked to resident *hxn* clusters. This larger clade also includes several species, token species like *Neurospora crassa*, *Magnaporthe oryzae* and *Trichoderma reesei*, that lack the *hxn* system and harbour only a lone copy of the *hxnV* gene, suggesting a loss of the nicotinate assimilation pathway and probably a novel function for *hxnV* in these *Sordariomycetes*. The tree is rooted in the monophyletic HxnV clade of the *Saccharomycotina*. Colour code: Blue: *Saccharomycotina*; Purple: *Pezizomycetes*; Magenta: *Leotiomycetes*; Brown: *Sordariomycetes*; Black lines *Xylonomycetes;* Orange: *Dothideomycetes*; Cyan: non-*Aspergillus*, non-*Penicillium Eurotiomycetes*; Darker Cyan: *Penicillium;* Green: *Aspergillus*, except Olive green: section *Flavi;* Red: sections *Nidulantes/Versicolores*. Species names in red, those mapping outside their cognate phylogenetic clade suggesting HGT events. Species names in blue: those showing a duplication of the *hxnV* gene.

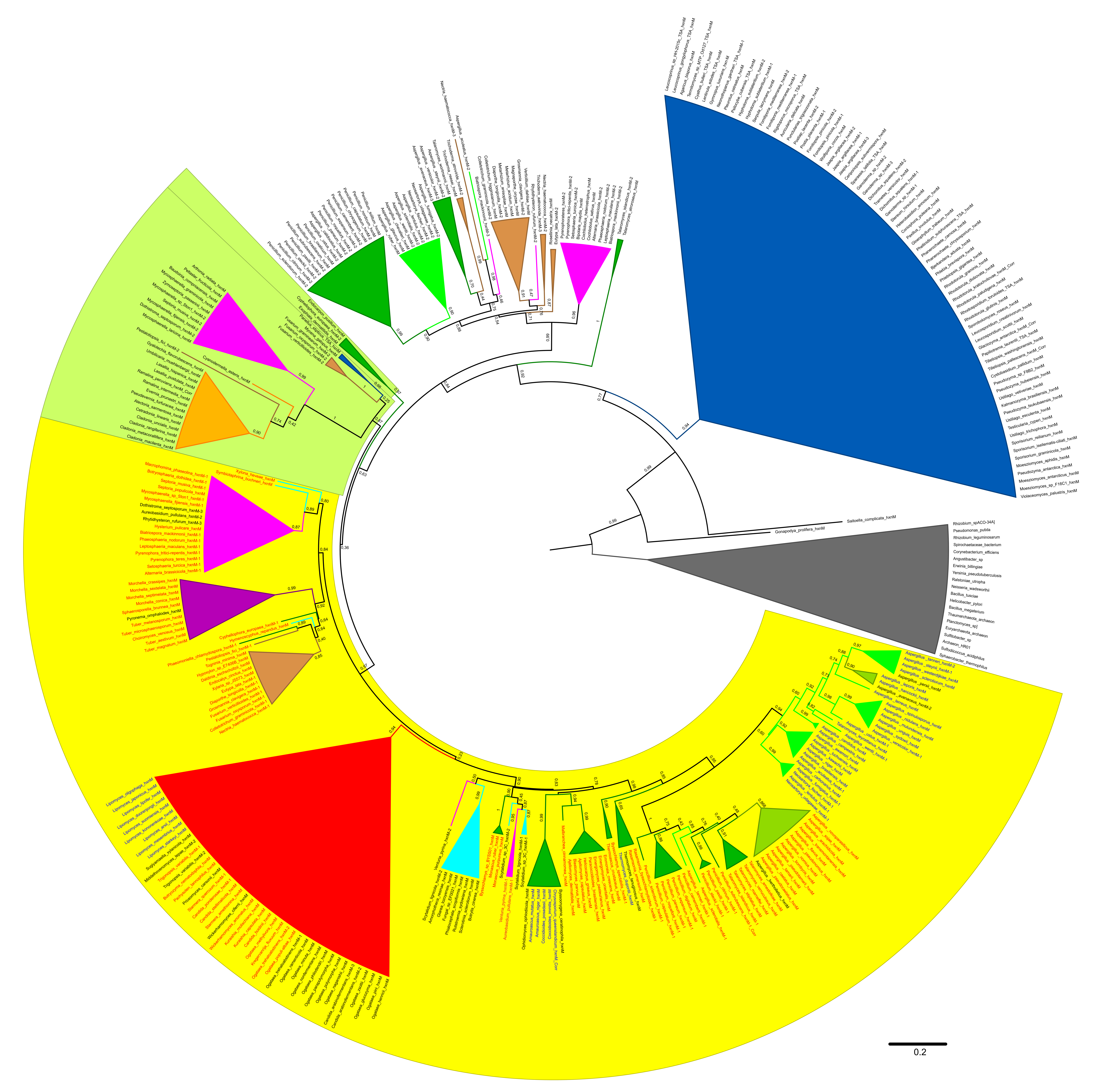
0.2

**Supplementary Figure S9. Maximum Likelihood Phylogeny of the HxnM putative imino-hydrolase protein.** All putative HxnM orthologues are >50% identical to the *A. nidulans* HxnM protein and >50% identical to the biochemically characterised cyclic imide hydrolase from *Pseudomonas putida* (strain YZ-26) (GenBank AAY98498). Methods used in the construction of the tree are detailed in the Materials and Methods section. Numbers in nodes are aLRTs (approximate Likelihood Ratio Tests). Names of species in black: Putative HxnM proteins encoded by genes (*hxnM*) not clustered with any other *hxn* gene. Names of species in blue: *hxnM* genes clustered only with *hxnN* genes (e.g., like in *A. nidulans,* see Figure 1 or Sup. Figure S5, or in *Lipomyces*). Names of species in red: *hxnM* included in a large cluster, encoding ≥ 4 genes. Highlighted in yellow: a monophyletic clade and putatively iso-functional clade comprising HxnMs from both clustered and un-clustered *hxnM* genes. Highlighted in light green: a clade comprising only proteins from un-clustered HxnM encoding genes, which appears as the outgroup to all clades with HxnMs from clustered genes detailed above. Colour code: Grey: prokaryotic outgroup (comprising both Bacterial and Archeal proteins); Blue: *Basidiomycota* (phylum); Red: *Saccharomycotina* (subphylum); Other colours: Pezizomycotina taxa; namely: Green: Eurotiomycetes; Olive green: *Aspergillus* section *Flavi* to highlight its anomalous position (HGT); Brown: *Sordariomycetes*; Magenta: *Dothideomycetes*; Orange: *Lecanoromycetes*; Cyan: *Leotiomycetes*; Purple: *Pezizomycetes*.

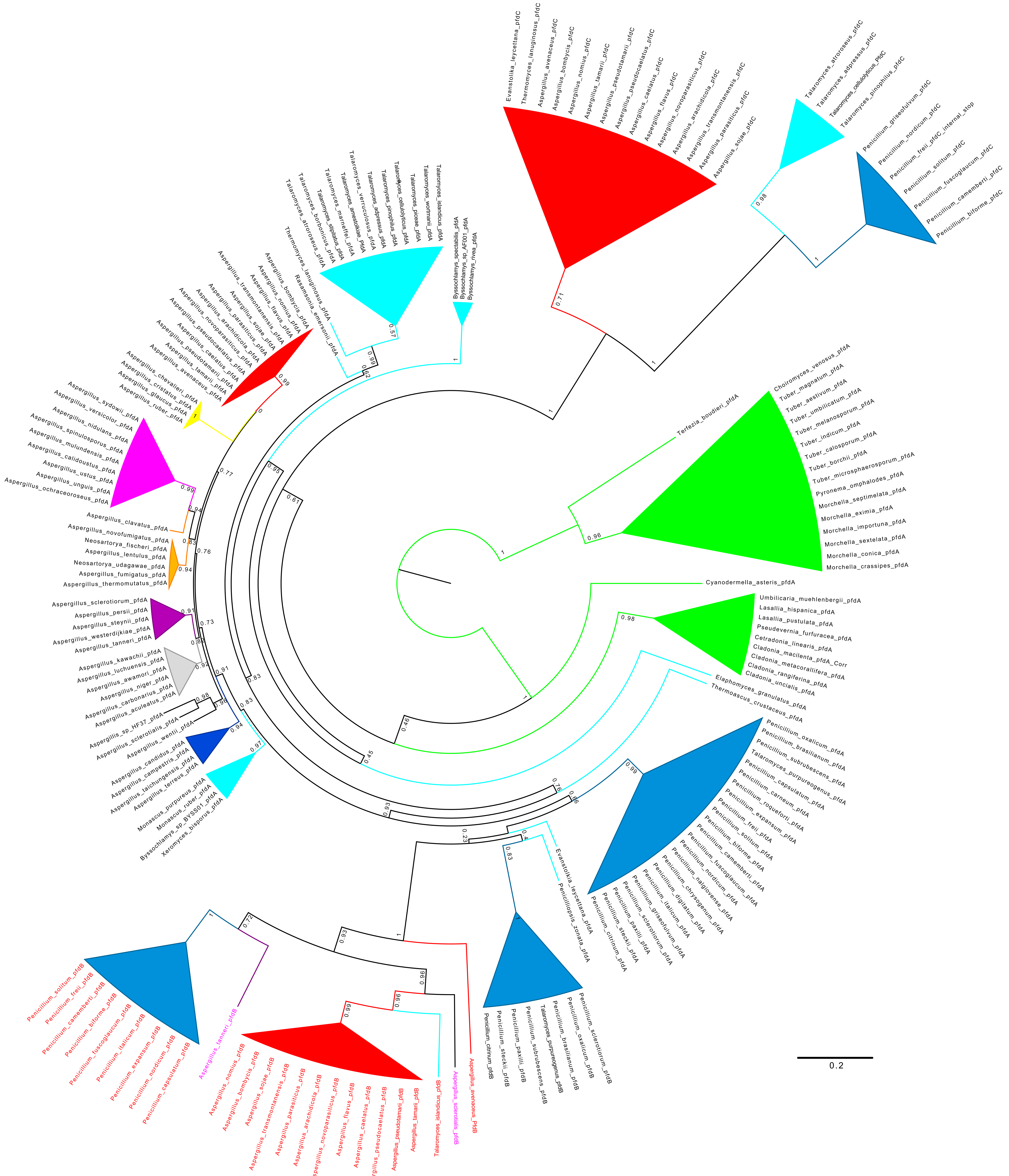

Cyanodermella_asteris_pfdA
Umbilicaria_muehlenbergii_pfdA
Lasallia_hispanica_pfdA
Lasallia_pustulata_pfdA
Pseudevernia_furfuracea_pfdA
Cetradonia_linearis_pfdA
Cladonia_macilenta_pfdA_Corr
Cladonia_metacorallifera_pfdA
Cladonia_rangiferina_pfdA
Cladonia_uncialis_pfdA
Elaphomyces_granulatus_pfdA
Thermoascus_crustaceus_pfdA
Choiromyces_venosus_pfdA
Tuber_magnatum_pfdA
Tuber_aestivum_pfdA
Tuber_umbilicatum_pfdA
Tuber_melanosporum_pfdA
Tuber_indicum_pfdA
Tuber_calosporum_pfdA
Tuber_borchii_pfdA
Tuber_microsphaerosporum_pfdA
Pyronema_omphalodes_pfdA
Morchella_septimelata_pfdA
Morchella_eximia_pfdA
Morchella_importuna_pfdA
Morchella_sextelata_pfdA
Morchella_conica_pfdA
Morchella_crassipes_pfdA
Terfezia_boudieri_pfdA
0.2

**Supplementary Figure S10. Phylogeny of the PfdA, B, and C paralogues in the *Eurotiales*.** The *pfdA* gene is ubiquitous in *Pezizomycotina* and encodes a well conserved protein of ~ 500 amino acids with a canonical peroxisome targeting sequence PTS-1 at its C-terminus. Generally, *Eurotiales* PfdAs are >72 % identical to the *A. flavus* protein. The *pfdB* and *pfdC* paralogues are restricted to specific taxa of the *Aspergillaceae* or *Trichocomaceae* families of the *Eurotiales* order; some species have two- while others have all three genes. The *pfdB* gene is regularly but not always, associated with the *hxn* gene cluster. The PfdB and C proteins are respectively, >60 % and >50 % identical to *A. flavus* PfdA. Typically, both paralogues are shorter than ubiquitous PfdA; multiple sequence alignments of the three paralogues (from one species) show a distinct gap in the middle of the alignment. All PfdAs and PfdBs feature a canonical PTS-1 (Neuberger et al., 2003) but some PdfCs appear to have lost the canonical signal sequence for peroxisome entry. All *Eurotiales* Pfd paralogue proteins included show the same domain organisation and their encoding genes have a conserved exon/intron structure with five exons (see Sup. Figure S12). In *Aspergillus*, the four introns in *pfdA* are confirmed by non-overlapping EST clones accessions DR703303 (introns 1 and 2 near the ATG) and CO136618 (introns 3 and 4 near the STOP). The exon 2 is always 59 nt long and exon 4 is always 74 nt long in nearly all Pfd paralogues. The size of the large central exon 3 distinguishes *pfdA* from its *B* and *C* paralogues. Tree construction was as detailed in Material and Methods, except that a Blosum 30 matrix was used for the BMGE alignment trimming. The species involved in the *hxn* cluster transfer (*Penicillium*, *Talaromyces*, *Aspergillus* section *Flavi*) are in red. Species names in magenta are *Aspergillu*s of sections other than *Flavi* which also include a *pfdB* gene in their *hxn* gene clusters. Colour code: Green: Outgroups *Pezizomycetes* and *Lecanoromycetes* (all PfdA; these taxa do not feature the *pfdBC* paralogues); Cyan: *Eurotiales* other than *Aspergillus* and *Penicillium*; Darker Cyan: *Penicillium;* Other colours: sections of the genus *Aspergillus.* Magenta: sections *Nidulantes*/*Versicolores*; Red: section *Flavi*; Orange: sections *Fumigati*/*Clavati*; Brown: sections *Terrei*/*Candidi*; Yellow: section *Aspergillus*; Grey: section *Nigri*; Purple: section *Circumdati*.

>*pfdA*
ATGGCCAAAGTGTTCGATGCGGCGGAAGgtatattgtttctcctctactccggcctgaattattgccagcctgatgtgctaattggcaaccaatcgt
tcagTCGCGAAACATAATACTCCAGACAGTTGCTGGGTGGTTTTATATGGAAAGTGTACGATgtacgaatctatccaagaattaaagcttcatttt
tcacatgttacacctcaaactaatccagtataacatattgtcatagGTCACCGACTTCCTCTCCGAGCACCCCGGTGGTTCCAAAATCATTCTGAAG
CTCGCTGGAAAGGATGCCACCGAAGAATACGACCCCATCCACCCCCCGGGCATATTAGAAGAAAACCTGAAGCCAGAAGCGATGCTTGGAACTGTCA
ACCCAGATACACTTCCAAAAGTGCAGGCAGAGCCAGTACCCTCGTCAAGCGAGGAGAATGAGGGTCCACCCCCAATGGAGTCTCTCCTCAACATGGA
TGACATCGAACAGGTGGCTACTAAGAACGTGAGCAAGAAGGCGTGGGCCTACTACTATTCTGCATCTGATGACAAAATCTCCAAACATTTCAACACC
GAAGTTTACCGCTCAATTCTCTTGCGGCCACGGGTTTTCATTGATTGCACACAATGTGACCTGGATACTACGCTGCTGGGGCATAAGCTAGGAATGC
CAATTTATGTGTCCCCTGCCGCGATGGCTCGATTGGGCCATCCTGCTGGTGAGGCAGGTATCGCCGAAGCTTGCCGCAGCTTTGGAGCCATGCAGGT
TATCTCAAACAACGCATCTATGACCCCGGAGCAAATTGTAAAAGATGCAGCCCCCGACCAAGTGTTTGGATGGCAGATCTATGTTCAAATTGATCGC
AAGAAGAGCGAAGCGATGCTGGCACGGATAAACAAACTCAAGCAAATCAAATTCATTGTTCTGACTCTGGATGCCCCAGTGCCAGGCAAACGAGAAG
ATGACGAGCGGGGAAATGCTGTCGGAGCATCAGCGCCAGTTCCAAGCGCGGCGAAGGTGGCCGATAGTGCTGAAGATGAAACTTCTCGGATCAATCA
ATCATCTGGCGGTGTTGGAAAGCAGCTCTTCGCTGGAACAGATCCCTCACTGACCTGGAAAGAAACTTTGCCTTGGTTGGCTGAACGTACTAACTTG
CCAATTGTCCTAAAGGGCCTGCAAACTCATGAGGACGCCTATGTTGCCTCTCTCCATACACCTCAAGTCAAAGGCATTATTCTTTCGAATCATGGTG
GTCGTGCGCTCGACACTGCTCCCCCAGCAGTGCATACGTTGATGGAAATCAGAAAATATTGCCCGGAAGTATTCGATAGATTGGACGTTTGGGTCGA
CGGCGGCATCCGACGCGGTACAGATGTTGTGAAAGCGCTATGCTTGGGCGCGAAGGCCGTGGGAATCGGGCGACCTGCCCTGTGGGGCCTGGGGGCT
GGCGGTGTCGAGGGCGTTAAGCGAACTCTGCAGAgtatgtttcggaaccctgataatcgagaaacaaacgactgactgaggcttagTACTGGCGGAC
GAGACCAAGACATGCATGCGATTACTTGGTGTAGATTCTGTGGATAAGCTGGGCCCCCAACATgtaagactccttcgttactgtctcaatgacagcg
aggttccccatcaaagatactccgatgcttacactatatagATCAACACCCGTCTTCTTGAGCAACAGATTTATAATGGTCCATCAGGCCTCGAATC
CATACGGAGCGTTTTCCGAGCGAAATTATAG

>*pfdB*
ATGAACAAGGTTTTCACAGGTGCTGAAGgtacatcaatagttcctgaaactcaagactgatctaactgcccaagTTGCCCAACACAACAACGAGAAC
AGCTGCTGGGTTGTTCTTTACGGAAAGGTATATGATgtgagtttgccctgtggcttgcatagatttccatcacaatgcgctgatcgggacgagGTAA
CGCATTTGCTGTCAAGCCATCCTGGCGGCGCCCAAGCCATTTTGCGTGTGGGGGGTATAGATGCCACAGACGACTTCGATCCCATTCATCCTCCTGA
GACCATGCAGAGTCTCCAATCGGCACAAATCGGATCAGTTGACCGCGACGAGGAGCCATTTTGTGCAAGTCATACTACGGGTCCCACGGACGAAATC
GACGTTAACACCCTCTTGAACCTGGACGAGATAGAACAGGCCGCCAGCAAGGTAATCAGCAAGAAGGCATGGGCCTACTATTACTCGGCATCCGACG
ACAAGATAACCAAGGACTTTAACACACAAGTTTACCGATCACTGTTGTTACGTCCCCGAGTCTTTGTCGACTGCCAGAAATGCGATGTGGAGACCGA
GCTTTTGGGATGGAAGCTTGGTCTACCGATATATATATCTCCCACCGCAATGGCTCGTTTAGGACATCCTAGCGGGGAGGCGGGACTCGCAGAGGCA
TGCAAAGCTTTTGGAGCGCTCCAGATCATCGCAAATAATTCGTCACTCTCCCCTGAACAGGTGGTGGCCAATGCAGCACCGACGCAAGTATTTGGAT
GGCAGCTGTATGTCCAACGAGACCGCAGTGCCAGCGAAGCGATGCTCGCACGGGTGAACAAGTTAGATGCGATCAAGTTTGTGGTACTCACCCTTGA
TGCGCCCGTCTCGGGAAAACGAGAAGATGATGAACGCATCAATATGAAGGCTCATCCAGCAGGCAGTGTCAGTGCCCAGCTTTTCGCTGGCACAGAT
CCGTCTCTCACCTGGCACGAGACTCTTGAGTGGCTGTCACGACACACGACGAAGCCTATCATCCTCAAGGGCCTGCAGACTCATGAGGATGTTGCTA
TTGCCGCCCGGCATACACCCCTCGTCCGAGCTGTTATTCTATCTAATCATGGTGGGAGGTCTCTGGACACGGCACCACCGGCCGTACACACCCTCCT
TGAAATACGGAAGTACTGTCCCTACGTTTTCATGAAGATGGAGGTGTGGGTAGATGGCGGTATTCGACGAGGTACCGACGTAGTTAAAGCGCTTTGC
CTTGGTGCGAAGGCTGTTGGCATTGGACGACCGGCTCTTTGGGGCCTTGGGGCCGGTGGAGTGAAGGGAGTTGAACGAACGCTACAAAgtgcgccgc
ctattctcttctgtgattgtcggtggccatctgctaatatcccgtcttcgtcttgtagTTCTGCTGGAGGAGACCAAGACCTGCATGCGACTCTTAG
GAGCGACCAGCATCAAGGAATTGGGTCCCCACTACgtaagggatgccttctttctctttcccgataagcactgactggattctgtgcagGTCAACTC
CAGCAGTGTGGAGCAGCAGATCTATCGGACGCCGCCGAATGTTGAGGATGAGGAAGCTGTAGCATACAAGGGGAAGCTCTAG

>*pfdC*
ATGATAGGGCGCCGGGAAAgtaagtcactgaacaggaaagtatgagggcaaaagtaattgatgtgacttagTTGCCAAACACAACTCCGCCGATAGT
TGCTGGGTTGTCCTCTATGGCAAAGTATACGACgtgagtagaacccgacaactttctcatcaggcgcattccatttaacaaatctagGTCACTAACT
TTCTCGAAAATCATCCCGGGGGCTCTGCGGCTATCCTGGCCTTGGCTGGGAAAGATGCAACACAGGAGTACGATACCCTACACCCCCCAGGTCTATT
AGAAGAGTACCTTGCTCCTGAGGCATGCTTGGGCGTCCTATCCGCATCCACGTCAGAGGCTGCTGAGCCAACAAGCATTTCCCAACACTCAGCATCT
AAAGAGACTTCCGAGGAAGCACCACTATCGTCACTGTTGAATCTCGCCGAGATCGAACAAGTTGCCAAGCGAAAACTCAGCCCAAAAGGGTGGGCGT
ATTACTCCTCTGCAACGGATGACGGCATCACAAAAGCGCACAACAATCTCATCTACCGTTCGATTTTACTAAGACCACGAGTTTTCATAGACTGCCG
CGAATGCGATCTTTCCATCGATTTTCTCGGCCTCAAGCTCGGACTACCGATCTACATTTCTCCAGCAGCAATGGCTCGTCTCGCCCACCCACAAGGC
GAAGCTGGCATTGCAGCAGCGTGCCGAAAGTTCGGGGCGATGCAAATTATCTCCCACAACGCCTCGATGGCGACACAGGAAATCGTCGCAAACGCCC
ACCCAGACCAAGTCTTCGGCTGGCAACTATATTGCCTAAAAGACGTGCGGCGTAGCGAGAAGCGGATCGCCGAAATCAACTCAATCAAAGCCATCAA
ATTCATCTGTCTCACGCTTGACGCTCCATTCCCCGGTAAACGCGAGATCGAGGAGCGACAAAAGATGGAAGAGCTGCGCATCGCTGGCGCGGCTTCG
TCGTCCCAGGTCTGGGGAACTGATGCCTCGCTCACGTGGGAGAGAACGCTAGATTGGCTCCGGATGCATACATCCCTGCCGATTATTCTAAAGGGTA
TTCAAACTTATGATGACGTTATACTCGCCGCTAAGCATGCTCCCCAGGTCCGGGGGATTGTACTTTCGAACCACGGTGGCCGCGCGTTGGACACCGT
ATCGACTCCTATGCATGTCCTACTGGAGATTAGGCGCTTCTGTCCAGAGGTCTTGGATCGATTGGATGTCATCGTTGACGGTGGCATCCAAAGAGGG
ACGGATGTCGCCAAGGCTCTGGCACTTGGTGCTAAGGCTGTGGGGATTGGACGAGCAGCGTTGTATGGTCTTGCAGCGGGTGGGCAAGAGGGTGTCG
AAAGGACGCTACAGAgtaggtttatcacaatcgctgaacatgtagcgagcttttcataactgatttgtccagTTCTTGCCGACGAAACGGCAACTGC
TATGAGACTTCTCGGTGTTCAGCGTGTTGACCAGCTGTCATTGCAGCATgtgggttgccatgtcctgaccgttatacaattactaaaatcactccag
GTAAATACCCAGCTCGTGGACTCACAGATCTTCAAGTCCGGAAGCTCGATGTCGTCAGTGGAGAGGGCATTTCCAGCCCGCGCCAAATTCTGA

**Supplementary Figure S11. Intron positions in *pfdA*, *pfdB*, and *pfdC* paralogues in *Aspergillus nomius*.** According to the data of *A. nomius* whole genome sequence project (JNOM00000000, https://www.ncbi.nlm.nih.gov/nuccore/JNOM00000000), all three *pfd* paralogues have four introns in conserved positions, two near the 5'end of the CDS and two near the 3'. Introns (lower case letters) are highlighted in blue (splice site consensus sequences: 5'-donor, lariat branch point sequence, and 3'-acceptor) and grey (other intronic sequences). The sizes of the small exons 2 and 4 are absolutely conserved throughout the orthologues. The size of the large central exon 3 distinguishes the three paralogues. In the ubiquitous *pfdA* gene exon 3 is 30 codons longer than the exon 3 of *pfdB* and 20 codons longer than exon 3 of *pfdC*. The canonical PTS-1 (Neuberger et al., 2003) has been lost in some but not all PfdC paralogues that supports a scenario of gene duplication of *pfdA* with simultaneous or subsequent cluster integration (mean similarity between *A* and *B* paralogues 65% compared with 88% of *A* orthologues among themselves). The third paralogue *pfdC* (mean identity between *A* and *C* paralogues 57%) present in section *Flavi*, and in a number of *Talaromyces* and *Penicilium* species and in a few species of other clades (supplementary fig. S11), which is consistent with a basal duplication followed by several episodes of loss completely unrelated to the evolution of the *hxn* cluster.

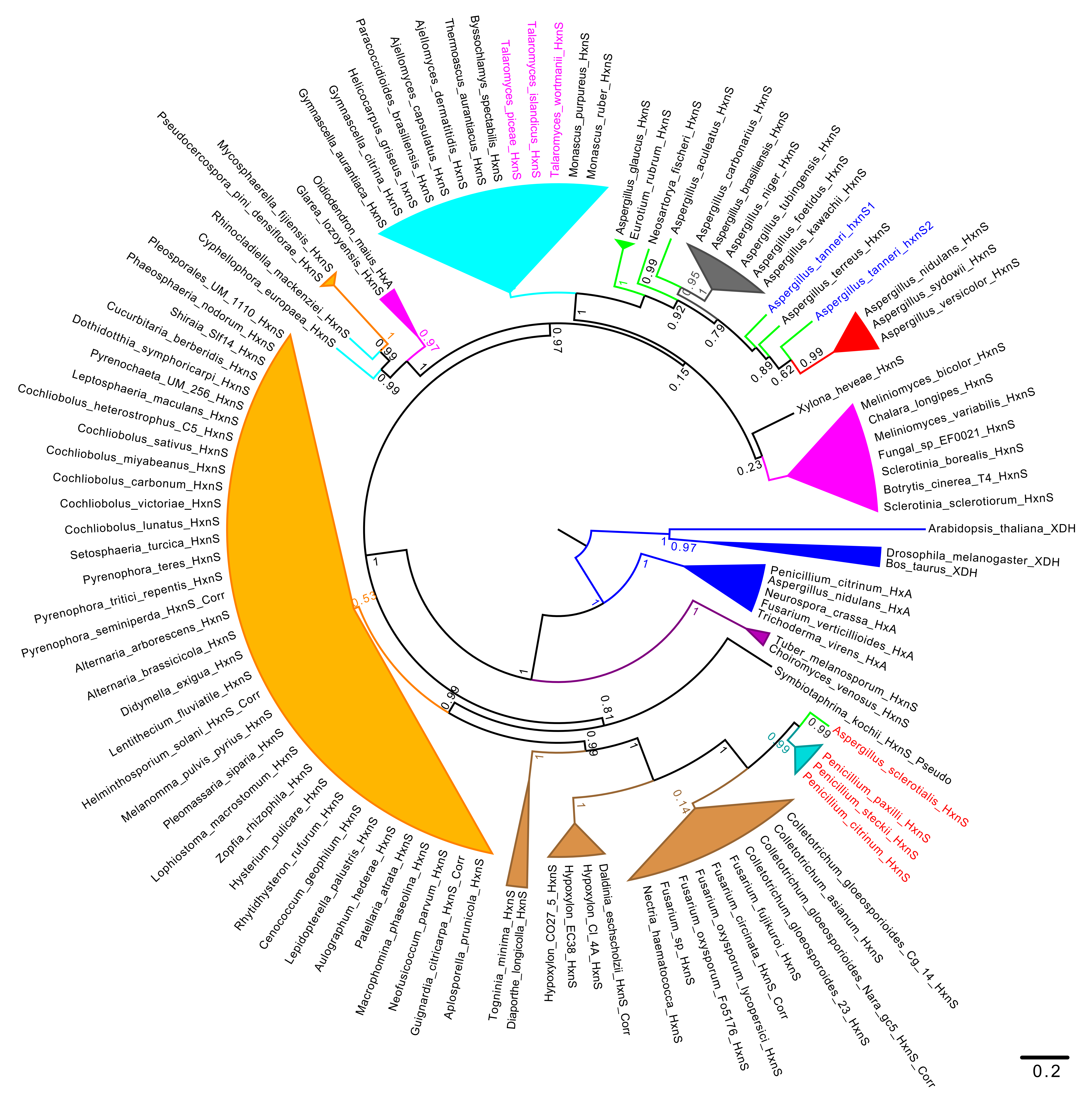

Monascus_ruber_HxnS
Monascus_purpureus_HxnS
Talaromyces_wortmanii_HxnS
Talaromyces_islandicus_HxnS
Talaromyces_piceae_HxnS
Byssochlamys_spectabilis_HxnS
Thermoascus_aurantiacus_HxnS
Ajellomyces_dermatitidis_HxnS
Ajellomyces_capsulatus_HxnS
Paracoccidioides_brasiliensis_HxnS
Helicocarpus_griseus_hxnS
Gymnascella_citrina_HxnS
Gymnascella_aurantiaca_HxnS
Oidiodendron_maius_HxA
Glarea_lozoyensis_HxnS
Mycosphaerella_fijiensis_HxnS
Pseudocercospora_pini_densiflorae_HxnS
Rhinocladiella_mackenziei_HxnS
Cyphellophora_europaea_HxnS
Pleosporales_UM_1110_HxnS
Phaeosphaeria_nodorum_HxnS
Shiraia_Slf14_HxnS
Cucurbitaria_berberidis_HxnS
Dothidotthia_symphoricarpi_HxnS
Pyrenochaeta_UM_256_HxnS
Leptosphaeria_maculans_HxnS
Cochliobolus_heterostrophus_C5_HxnS
Cochliobolus_sativus_HxnS
Cochliobolus_miyabeanus_HxnS
Cochliobolus_carbonum_HxnS
Cochliobolus_victoriae_HxnS
Cochliobolus_lunatus_HxnS
Setosphaeria_turcica_HxnS
Pyrenophora_teres_HxnS
Pyrenophora_tritici_repentis_HxnS
Pyrenophora_seminiperda_HxnS_Corr
Alternaria_arborescens_HxnS
Alternaria_brassicicola_HxnS
Didymella_exigua_HxnS
Lentithecium_fluviatile_HxnS
Helminthosporium_solani_HxnS_Corr
Melanomma_pulvis_pyrius_HxnS
Pleomassaria_siparia_HxnS
Lophiostoma_macrostomum_HxnS
Zopfia_rhizophila_HxnS
Hysterium_pulicare_HxnS
Rhytidhysteron_rufurum_HxnS
Cenococcum_geophilum_HxnS
Lepidopterella_palustris_HxnS
Aulographum_hederae_HxnS
Patellaria_atrata_HxnS
Macrophomina_phaseolina_HxnS
Neofusicoccum_parvum_HxnS_Corr
Guignardia_citricarpa_HxnS_Corr
Aplosporella_prunicola_HxnS
Togninia_minima_HxnS
Diaporthe_longicolla_HxnS
Hypoxylon_CO27_5_HxnS
Hypoxylon_EC38_HxnS
Hypoxylon_CI_4A_HxnS
Daldinia_eschscholzii_HxnS_Corr
Nectria_haematococca_HxnS
Fusarium_sp_HxnS
Fusarium_oxysporum_Fo5176_HxnS
Fusarium_oxysporum_lycopersici_HxnS
Fusarium_circinata_HxnS_Corr
Fusarium_fujikuroi_HxnS
Colletotrichum_gloeosporoides_23_HxnS
Colletotrichum_gloeosporioides_Nara_gc5_HxnS_Corr
Colletotrichum_asianum_HxnS
Colletotrichum_gloeosporioides_Cg_14_HxnS
Penicillium_citrinum_HxnS
Penicillium_steckii_HxnS
Penicillium_paxilli_HxnS
Aspergillus_sclerotialis_HxnS
Symbiotaphrina_kochii_HxnS_Pseudo
Choiromyces_venosus_HxnS
Tuber_melanosporum_HxnS
Trichoderma_virens_HxA
Fusarium_verticillioides_HxA
Neurospora_crassa_HxA
Aspergillus_nidulans_HxA
Penicillium_citrinum_HxA
Bos_taurus_XDH
Drosophila_melanogaster_XDH
Arabidopsis_thaliana_XDH
Sclerotinia_sclerotiorum_HxnS
Botrytis_cinerea_T4_HxnS
Sclerotinia_borealis_HxnS
Fungal_sp_EF0021_HxnS
Meliniomyces_variabilis_HxnS
Chalara_longipes_HxnS
Meliniomyces_bicolor_HxnS
Xylona_heveae_HxnS
Aspergillus_versicolor_HxnS
Aspergillus_sydowii_HxnS
Aspergillus_nidulans_HxnS
Aspergillus_tanneri_hxnS2
Aspergillus_terreus_HxnS
Aspergillus_tanneri_hxnS1
Aspergillus_kawachii_HxnS
Aspergillus_foetidus_HxnS
Aspergillus_tubingensis_HxnS
Aspergillus_niger_HxnS
Aspergillus_brasiliensis_HxnS
Aspergillus_carbonarius_HxnS
Aspergillus_aculeatus_HxnS
Neosartorya_fischeri_HxnS
Eurotium_rubrum_HxnS
Aspergillus_glaucus_HxnS
0.2

**Supplementary Figure S12. Phylogeny of HxnS in the *Pezizomycotina***. The tree is shown in a circular, cartoon form. Tree was constructed as indicated in Materials and Methods, except that MAFFT E-INS-i was used for the alignment. The HxnS sequences included are from Ámon et al. (Amon et al., 2017) to which orthologue proteins from some more recently published *Aspergillus*, *Penicillium* and *Talaromyces* species were added: *A. sclerotiales*, *A. tanneri*, *P. steckii*, *T. wortmannii*, and *T. picheae*. Colour code: Blue: Outgroups, including non-fungal xanthine hydroxylases and selected HxA orthologues; Purple: *Pezizomycetes*; Magenta: *Leotiomycetes*; Brown: *Sordariomycetes*; Black (lines): *Xylonomycetes*; Orange: *Dothideomycetes*; Cyan: non-*Aspergillus*, non-*Penicillium Eurotiales* (such as *Talaromyces*); Darker Cyan: *Penicillium* genus*;* Red: Sections *Nidulantes/Versicolores* (also called subgenus *Nidulantes* by Houbraken and Samson, 2011); Grey: Section *Nigri;* Green: all other *Aspergilli*. The three *Penicillium* species and *Aspergillus sclerotialis*, which cluster together within the *Sordariomycetes* are shown in red. The duplicated HxnS sequences of *A. tanneri* are shown in blue. The three *Talaromyces* species which conserve HxnS are shown in magenta.

**S1 Table. *A. nidulans* strains used in this work.** (All strains are *veA1* mutant)

| Strain | Genotype | Purpose | Reference |
|---|---|---|---|
| CS308 | *pyroA4 hxn6* | recipient strain of transformation experiment; sequencing | this work |
| CS3095 | *areA600 biA1 sb43* | mRNA expression analysis | [1] |
| FGSCA26 | *biA1* | mRNA expression analysis | [2] |
| FGSCA872/ CS51 | *hxnR*$^{c}$*7 biA1* | growth test; enzyme assay; mRNA expression analysis | [3] |
| HZS.135 | *hxB20 biA1* | growth test; Northern analysis; RT-PCR | provided by S. Amillis |
| HZS.136 | *hxnRΔ::zeo pantoB100* | mRNA expression analysis | [4] |
| HZS.145 | | mRNA expression analysis; sequencing | [4] |
| HZS.216 | *xprD1 biA1 pabaA1* | mRNA expression analysis | [1] provided by Vicky Sophianopoulou |
| HZS.697 | *hxn7 pabaA1* | Southern analysis | this work |

Explanation of mutant alleles, which are not described in the text: *sb43* is a non-functional allele of the sulphate transporter SB [5], *veA1* mutation in *velA* results in profuse conidiation regardless of the presence or absence of light [2]. Other gene symbols refer to auxotrophies: *biA1*, biotin; *pabaA1*, p-aminobenzoic acid; *pantoB100*, pantothenic acid;and *pyroA4*, pyridoxine.

### S2 Table. Primers used in this work

| | |
|---|---|
| **RT-qPCR** | |
| hxnP ReTi frw | 5'- tgtacttctttgactgcatgg -3' |
| hxnP ReTi rev | 5'- gagtattcgttgcccttgag -3' |
| hxnS ReTi frw | 5'- gagcatttctatcttgagacga -3' |
| hxnS ReTi rev | 5'- ccattgtgttctgggtactg -3' |
| hxnT ReTi frw | 5'- ctcgaccagtttctacacgac -3' |
| hxnT ReTi rev | 5'- ccgagatgatttcaaggacga -3' |
| hxnY ReTi frw | 5'- gtcattcttcgatcctctcacc -3' |
| hxnY ReTi rev | 5'- ggttgagtttctcgtcttcgt -3' |
| hxnZ ReTi frw | 5'- cgctgtatttcaactttctccc -3' |
| hxnZ ReTi rev | 5'- cagtagttgcggtaggtcag -3' |
| hxnR ReTi frw | 5'- cggcttctgttctactacagg -3' |
| hxnR ReTi rev | 5'- cagtctaggtctggaaagtctc -3' |
| actin ReTi frw | 5'- ggtatcatgatcggtatggg -3' |
| actin ReTi rev | 5'- tatctgagtgtgaggatacca -3' |
| hxnX ReTi frw | 5'- cttgtatcatctccacgacgg -3' |
| hxnX ReTi rev | 5'- ggctaaacactctccctctg -3' |
| hxnW ReTi frw | 5'- ggtatggcggtttatccctg -3' |
| hxnW ReTi rev | 5'- cctctcaggattccttgaaagac -3' |
| hxnV ReTi frw | 5'- gacccgtatattctcctcgtg -3' |
| hxnV ReTi rev | 5'- gaaatgaactattccgcgtaacag -3' |
| hxnM ReTi frw | 5'- aagacctaccgcatgattacag -3' |
| hxnM ReTi rev | 5'- cagcaattccgtcatctcct -3' |
| hxnN ReTi frw | 5'- cattgcatggttctatcttggg -3' |
| hxnN ReTi rev | 5'- atccatacaatcccagaatgct -3' |
| AN9159 ReTi frw | 5'- gatcagcaaaggtgggagag -3' |
| AN9159ReTi rev | 5'- cctccattacacataacaccga -3' |
| AN9162 ReTi frw | 5'- gaccaggagaacattccgag -3' |
| AN9162 ReTi rev | 5'- cattgattgcatccagacaagac -3' |
| AN10825 ReTi rev | 5'- gattattgttgccgccactg -3' |
| AN10825 ReTi rev | 5'- atatccatgttcgtgccctc -3' |
| AN6517 ReTi rev | 5'- gacaacttcaattctctgcctg -3' |
| AN6517 ReTi rev | 5'- tcatcactagcgttcaactcc -3' |
| **cDNA amplification** | |
| hxnX cDNA EcoRI frw | 5'- tttttttgaattcatgcccatcccagttgcagagaaac -3' |
| hxnX cDNA PstI rev | 5'- tttttttctgcagtcataaccgcgatgctacctgttc -3' |
| hxnM cDNA EcoRI frw | 5'- tttttttgaattcatgggaaagaagcgagttctcgtatc -3' |
| hxnM cDNA PstI rev | 5'- tttttttctgcagctatttcagcactgatgaaggagg -3' |
| hxnW ispan frw | 5'- atgtccaagttctccttgaaag -3' |
| hxnW 3UTR rev | 5'- gctcaacacagccggaatc -3' |
| hxnV cDNA EcoRI frw | 5'- tttttttgaattcatggctcgcagcgcagatcac -3' |
| hxnV cDNA PstI rev | 5'- tttttttctgcagtcacaagaagaatccaaaatactcctcc -3' |
| hxnN -69 5UTR frw | 5'- cgacaagaaatgaaaggctcctag -3' |
| hxnN 37 3UTR rev | 5'- cagagaggttcatttctcttctcaatc -3' |

| | |
|---|---|
| hxnT cDNA frw | 5'- agacagtcgaaaccatgggctc -3' |
| hxnT cDNA rev | 5'- cgtagtattcatgctctattgagcga -3' |
| hxnY cDNA frw | 5'- aagacgtacctaatggctccaac -3' |
| hxnY cDNA rev | 5'- ggctacgaaaccatagccgttg -3' |
| hxnZ cDNA frw | 5'- cgcactcctaaggcagtatgg -3' |
| hxnZ cDNA frw | 5'- gacacgactcctagatagcagca -3' |
| AN11197 XbaI frw | 5'- tttttttttctagacagctaatcctgcagtatagactcctc -3' |
| AN11197 KpnI rev | 5'- tttttttggtaccctcaatagagcatgaatactacgacac -3' |
| **sequencing** | |
| hxnX F366 | 5'- gtctgaggcgcagtatccag -3' |
| hxnX F716 | 5'- cacaggtcaactactggcttgg -3' |
| hxnX F1066 | 5'- gctgccatacttagcatctg -3' |
| hxnM F356 | 5'- aaattatcctcgacaagacctacc -3' |
| hxnM F709 | 5'- agagatgtggaggatatttgg -3' |
| hxnW seq 494F | 5'- gcagcgaagcagatgatagc -3' |
| hxnW seq 494R | 5'- gctatcatctgcttcgctgc -3' |
| hxnW seq 288F | 5'- cgtctcctctcaggattccttg -3' |
| hxnW seq 288R | 5'- caaggaatcctgagaggagacg -3' |
| hxnV seq 13F | 5'- cagatcacgaccaaggcttc -3' |
| hxnV seq 492R | 5'- gtaatgagtctggcgacccttc -3' |
| hxnV F513 | 5'- tcggacccgtatattctcct -3' |
| hxnV F1016 | 5'- ctaaggcaacaccgcaatatgtc -3' |
| hxnV F1811 | 5'- cgacgtccattctagcaatttctg -3' |
| hxnN seq 24F | 5'- cagacagtcgcaaagaaacg -3' |
| hxnN seq 483F | 5'- gcacggtgtaccagttacag -3' |
| hxnN seq 483R | 5'- ctgtaactggtacaccgtgc -3' |
| hxnN seq 832F | 5'- gtttggtacagacatcggcg -3' |
| hxnN seq 1401F | 5'- gtcgaagcattggtgaatcg -3' |
| hxnT frw | 5'- ctttgcgctgcagcaccgtgttgtccaag -3' |
| hxnT ReTi frw | 5'- ctcgaccagtttctacacgac -3' |
| hxnT ReTi rev | 5'- ccgagatgatttcaaggacga -3' |
| hxnY frw | 5'- caagcttgtacagcaagtacgcgaatcctg -3' |
| hxnY ReTi frw | 5'- gtcattcttcgatcctctcacc -3' |
| hxnY ReTi rev | 5'- ggttgagtttctcgtcttcgt -3' |
| hxnZ frw | 5'- cagatcatcatggaaatcgaactcgacaag -3' |
| hxnZ ReTi frw | 5'- cgctgtatttcaactttctccc -3' |
| hxnZ rev | 5'- ctgactggatgcagtagttgcggtaggtc -3' |
| hxnZ seq 507 frw | 5'- gggacgtcgaacttcatcgc -3' |
| hxnZ seq 507 rev | 5'- gcgatgaagttcgacgtccc -3' |
| AN11197 2F | 5'- gcacgcttatcgtctccactg -3' |
| AN11197 336F | 5'- gtcttcgggctctgtctctg -3' |
| AN11197 800F | 5'- gtatgatgccaatacagtaaagctacc -3' |
| AN11197 1375F | 5'- cttctcccgcttcaatactacatacc -3' |
| AN11197 1958F | 5'- agagatacagaacatgcatttcttccc -3' |
| AN11197 ReTi rev | 5'- cagtctaggtctggaaagtctc -3' |
